# Computational Primitives for Cost-Benefit Decision-Making

**DOI:** 10.1101/2024.12.09.627657

**Authors:** Lara I. Rakocevic, Luis D. Davila, Cory N. Heaton, Dirk W. Beck, Raquel J. Ibanez-Alcala, Safa B. Hossain, Neftali F. Reyes, Andrea Y. Macias, Alexis A. Salcido, Danil Tyulmankov, Serina A. Batson, Sabrina M. Drammis, Kenichiro Negishi, Paulina Vara, Atanu Giri, Sofia M. Gutierrez, Travis M. Moschak, Ki A. Goosens, Alexander Friedman

**Author notes:** Denotes authors who have equal second authorship (Luis D. Davila, Cory N. Heaton, Dirk W. Beck, Raquel J. Ibanez-Alcala, Safa B. Hossain, Neftali F. Reyes, Andrea Y. Macias, Alexis A. Salcido, Danil Tyulmankov, Serina A. Batson, Sabrina M. Drammis, Kenichiro Negishi, Paulina Vara, Atanu Giri). Denotes corresponding senior authors (Ki A. Goosens, Alexander Friedman).

## Abstract

Cost-benefit decision-making is a critical process performed by all organisms, including humans. Various factors, including risk^1,2^, uncertainty^3^, age^4^, sex^5^, and neuropsychiatric disorders^6^, can alter decision-making. To explore cost-benefit decision-making in humans, we developed a comprehensive task and analysis framework that presents participants with a series of approach-avoid trade-offs across a variety of contexts. With this system, we found that cost-benefit decisions in humans are made using a set of computational strategies that may be used for integrating costs and rewards, which we term ‘decision-making primitives’. We further show that these decision-making primitives are used by rodents performing a similar decision-making task^7^. We find that utilization of these primitives in both rodents and humans shifts based on factors like hunger and sex, and that individuals use primitives differently. We additionally demonstrate that using a naturally-inspired neural network architecture generates output that overlaps with human and rodent performance over a non-constrained neural network. This novel conceptual framework, by isolating discrete ‘decision-making primitives’, has potential to help us identify how different brain regions give rise to decision-making behavior, as well as to facilitate better diagnosis of neuropsychiatric disorders and development of naturally-inspired artificial intelligence systems of decision-making.

## Introduction

Decision-making is critical for daily functioning for nearly all organisms^8,9^, and can range from relatively inconsequential, like the estimated 200 food-related decisions an individual makes every day^10^, to life-altering decisions such as choosing a career to pursue. Pervasive and persistent abnormalities in decision-making are a trans-diagnostic symptom of various neuropsychiatric disorders^11,12^. Thus, understanding decision-making tendencies in normal and abnormal states could lead to methods for improving general decision-making^13,14^, aiding in neuropsychiatric disorder diagnosis^15^, and treatment^16^. In our work, we have identified discrete and heterogenous decision-making ”primitives”, computational strategies for valuation and integration of cost and reward, that shift in utilization depending on the task context and internal states in participants like hunger or tiredness.

A major challenge hindering our ability to understand decision-making processes is one applicable to most neuroscience research: discerning how different biological systems, like genes, cells, or neuronal circuits, lead to different behaviors^17^. To do so, quantifying the computational principles of the biological system of interest is a necessity; understanding these principles enables subsequent studies to accurately relate activity, ranging from gene expression, ionic conductance, or neuronal firing to distinct behaviors^18^. Many studies quantify principles in the context of “simple” well characterized organisms with a limited behavioral repertoire, such as rhythmic firing behavior in the pyloric circuit of crabs^19^, navigation in zebrafish^20^, or eating behavior in roundworms^21^. Recent advances have also started to break down rodent motor movements into simpler components that can be more specifically linked to its associated neural activity^22^. Perhaps there are primitives in decision-making as well, which may be used to identify the mechanisms which give rise to decision-making behaviors like with motor primitives in simpler organisms.

Intuitively, we like to think that human behavior is more complicated than that of simple organisms, yet many studies of human behavior do not explore the possibility that different strategies may be used across, and even within, individuals. However, understanding human behavioral heterogeneity is extremely important. Averaging behavior or other measures (e.g. neuronal or endocrine) across individuals to compare two or more groups may yield misleading results^23^. Heterogeneity is also important in clinical contexts because it may contribute to individual differences in the presentation of psychiatric disorders^24,25^, and perhaps differences in symptom presentation across days within single individuals^26^. We examined behavioral strategies for valuating and integrating costs and rewards both across and within individuals during cost-benefit decision-making, a cognitively complex computation that contributes to many human behaviors, and is also observed in most organisms^8,9^. Perturbations in cost-benefit decision-making are thought to contribute to many psychiatric and neurological conditions^11,12^, including substance-use disorder, post-traumatic stress disorder, Huntington’s Disease, Traumatic Brain Injury, and others. Thus, it is critically important to be able to measure and quantify decision-making on an individual level. Decision-making primitives may be a useful tool for quantifying changes in decision-making on an individual basis, ultimately aiding in diagnosis and treatment of neuropsychiatric disorders.

We developed a task framework where human participants rated how likely they were to pursue different combinations of rewards and costs of different values within natural decision-making narratives. We found that human participants use distinct behavioral strategies to integrate costs and rewards, suggesting each strategy is a behavioral ‘primitive’. We further demonstrate that behavioral decision-making primitives are observed in rodents performing a similar cost-benefit approach-avoid decision-making task. In addition, we found that individuals used these primitives in different proportions, and that this distribution shifted with states like hunger or tiredness. We finally show how a naturally-inspired neural network forms decision-making strategies more similar to human primitives as compared to a classic neural network.

## Results

### A task battery for context-dependent human decision-making

To better examine cost-benefit decision-making under different contexts, we created a battery of four decision-making tasks:1) an approach-avoid task which weighs situational costs and benefits, 2) a social task which weighs interpersonal costs and benefits, 3) a moral task which weighs ethical rights and wrongs, and 4) a probabilistic task which uses the same costs and benefits as in approach-avoid but introduces an element of chance when achieving the outcome (**Extended Data Fig. 1a; Methods: Decision-making application).** For each context, university students were presented with a list of story topics and asked to select those that they felt were most relevant to their own lives **(Extended Data Table 1: Participant Metadata);** each participant was then presented with 8-12 stories from the topic lists they identified as personally relevant. Each participant completed 3-4 stories per session, across 2-3 sessions held on separate days; participants were asked to provide a subjective rating of how hungry and tired they felt at the start of each session (on a scale of 0-100). Heart rate and pupillary diameter were tracked throughout all sessions. This was run on a web-app, developed as an open-access software suite written in Python, HTML, and CSS to be easily ported over and adapted by other labs for experimental use (https://github.com/lrakocev/dec-making-app/tree/raq_dev_utep, **Database User Guide** section in Supplemental Methods and Materials document).

Each story is split into three distinct phases: context setting (**Fig. 1a**), preference normalization (**Fig. 1b,c**), and response collection **(Fig. 1d)**. At the beginning of each session, participants are presented with a list of story topics from which they choose their themes of interest (**Fig. 1a**). From there, a story is presented to the participant, followed by a list of details related to the story, a subset of which is positive (i.e. getting a free ride somewhere) and a subset of which is negative (i.e. getting stuck in traffic on the way) (**Fig. 1b,c**). Once the participant has rated their preferences from 0 to 100 for each detail, the app chooses four preferences that are equally spaced and span over the full range of the point-space of ratings, for both costs and rewards (**Extended Data Fig. 1b-i; Methods: Normalization of preference ratings**). Using these four cost and four reward preferences, we create 16 trials where each cost and reward are paired, then ask the subject their likelihood of taking each cost-benefit offer (**Fig. 1d,e**). Thirty-five participants completed a total of 982 stories across the four contexts (**Fig. 1f**). In this way, we were able to leverage the human ability to imagine different scenarios and outcomes to ask about cost-benefit decision-making across varied contexts (**Fig. 1g, Extended Data Fig. 1j-ac; 2a-x**).

**Fig. 1.**
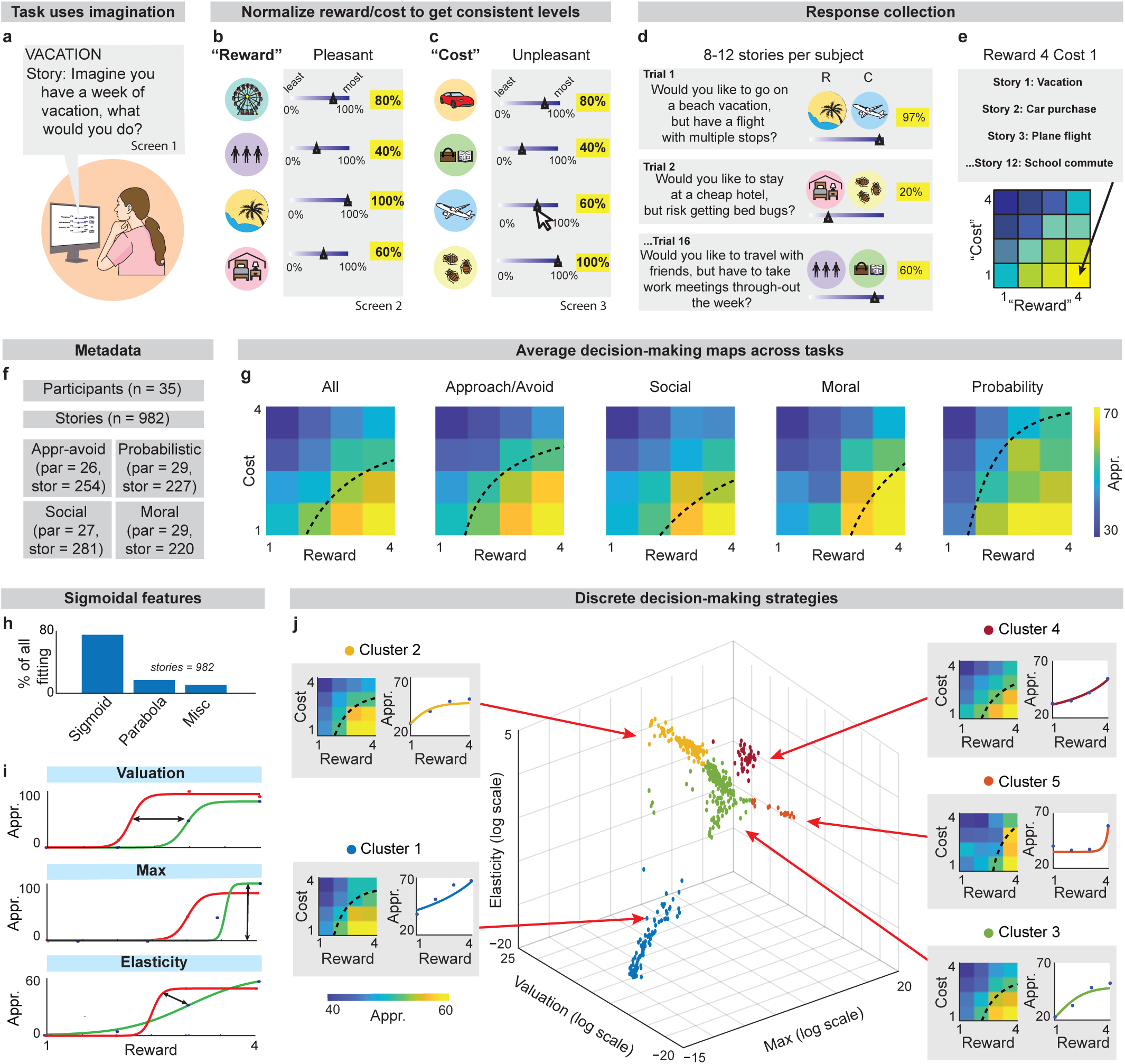
Constrained decision-making heterogeneity across neuroeconomic, social and moral tasks. **a.** Task begins by describing the context and setting of a scenario that will be used for the rest of the task. **b.** Participants are asked to rate how highly they value positive attributes about the context from 0 to 100. **c.** Participants then rate (from 0 to 100) how much they want to avoid negative attributes that can be associated with the task context. This sequence of steps allows for controlling for individual preferences **d.** The participant is then presented with a series of trade-offs where they rate their approach or avoidance on a scale from 0-100, enhancing the task’s statistical power. **e.** Participants repeat this procedure for 8-12 stories/sessions to gain statistical power for each combination of costs and rewards. **f.** Tasks metadata **(Extended Data Table 1: Participant Metadata). g.** Decision-making maps, with dashed lines representing where participants switch from preferring or avoiding trade-offs. Decision-making maps across tasks are significantly different (3-way ANOVA Matlab, reward: F(1766, 3) = 89.502, p=6.4718×10^-54^, cost: F(1766, 3) = 94.963, p=5.57×10^-57^, task type: F(1766, 3) = 44.936, p=5.5531×10^-28^). **h.** Data collected were fit to sigmoidal functions (72.8% of stories), parabolic functions (14.4%), or were considered miscellaneous (12.8%). **i.** Sigmoidal functions were differentiated by their “valuation”, “max”, and “elasticity”. **j.** Applying fuzzy clustering to sigmoidal data reveals distinct clusters of behavioral strategies. Each point represents a psychometric curve of a single session while different colors delineate unique clusters (MPC=0.63697, n=35 participants, 693 sessions). Bhattacharya distance indicates clusters are separate from each other (**Extended Data Fig. 4c**). The average psychometric function for each primitive is shown next to the primitive itself and the decision-making map. Log-scale was used for plotting these points, as well as for the rest of the manuscript.

### Behavioral primitives in human decision-making

We first compared the behavioral strategies used in different decision-making contexts by comparing the average decision-making map for each context. This commonly used analytic approach^27–29^ suggests that different behavioral strategies are used in different decision-making contexts. We found that participants were significantly more cost-averse when making social or moral decisions than for approach-avoid decisions (**Fig. 1g**, Social or Moral vs App-Av, F(6,8693) = 8.6, p=2.39×10^-9^). Participants were also significantly less cost-averse when rewards and costs were probabilistic (**Fig. 1g**, Prob vs App-Av, Cost x Task interaction: F(3,5788) = 13.13, p=1.519×10^-5^). It may be the case that participants feel costs more heavily in social and moral contexts, due to the interpersonal element of such tasks. Similarly, they may feel it less heavily in the probabilistic task since the outcome is uncertain: they may or may not have to actually contend with the cost-benefit tradeoff due to the probabilistic element built in.

Because averaging across all participants can obscure individual differences, we developed a second more granular analytic pipeline in which we analyzed the behavioral strategy for every story completed by every participant. We found that 72% of data was best fit with the sigmoidal functions that are expected when rewards and costs span large ranges (**Fig. 1h; Methods: Psychometric function fitting**). We then put three features (“valuation”, “max”, and “elasticity”) from each sigmoidal curve (**Fig. 1i; Methods: Primitive Identification)** into a 3D vector and plotted these points in a 3-D ‘parameter space’ to compare variability in behavioral strategies across and within individuals. To our surprise, we observed that the behavioral strategies occupied a very limited portion of the parameter space that described all possible strategies. In addition, they form distinct clusters which we quantified using fuzzy c-means clustering coupled with the modified partition coefficient, used to find the optimal number of clusters (MPC=0.63697, n=693 sessions, **Fig. 1j, Methods: Identifying Clusters).** Even using an alternative method of calculating subjects’ strategies using decision-making boundaries, which included all data and not just sigmoidal functions, we found discrete clusters of decision-making strategies (MPC=0.73084, n=982 sessions, **Extended Data Fig. 3a, Methods; Decision-Map Based Clustering)**. Both techniques produce discrete clusters of sessions with similar psychometric summaries. Discreteness in this space implies a finite and quantifiable number of decision-making behaviors which allows for comparison across tasks, individuals, and potentially species. We define each cluster as a decision-making “primitive”: a building block behavior that can be combined to produce more complex behaviors.

To further demonstrate that the five clusters represent distinct behavioral primitives, we first compared them using features of pupil diameter and heart rate (average pupil diameter, number of local maxima in pupil diameter, and maximum heart rate; **Extended Data Fig. 3b-o, Methods: Physiological Features**). In the literature, pupil diameter has been linked to interest^30–32^ and elevated heart rate has been linked to arousal^33,34^ so examining features derived from this data might allow for additional insight about subjects’ state while completing the tasks. We found that each primitive has unique physiological correlates which differ from other those of other primitives (F(8,13378) = 8.87, p=3.44×10^-12^, 2-way ANOVA, **Fig. 2a, Methods; Physiological Features)**. For example, primitive two had the highest average pupil diameter suggesting greater possible attention/engagement, while primitive five had the highest number of maxima in pupil diameter, suggesting more shifts in attention/engagement across a trial.

**Fig. 2.**
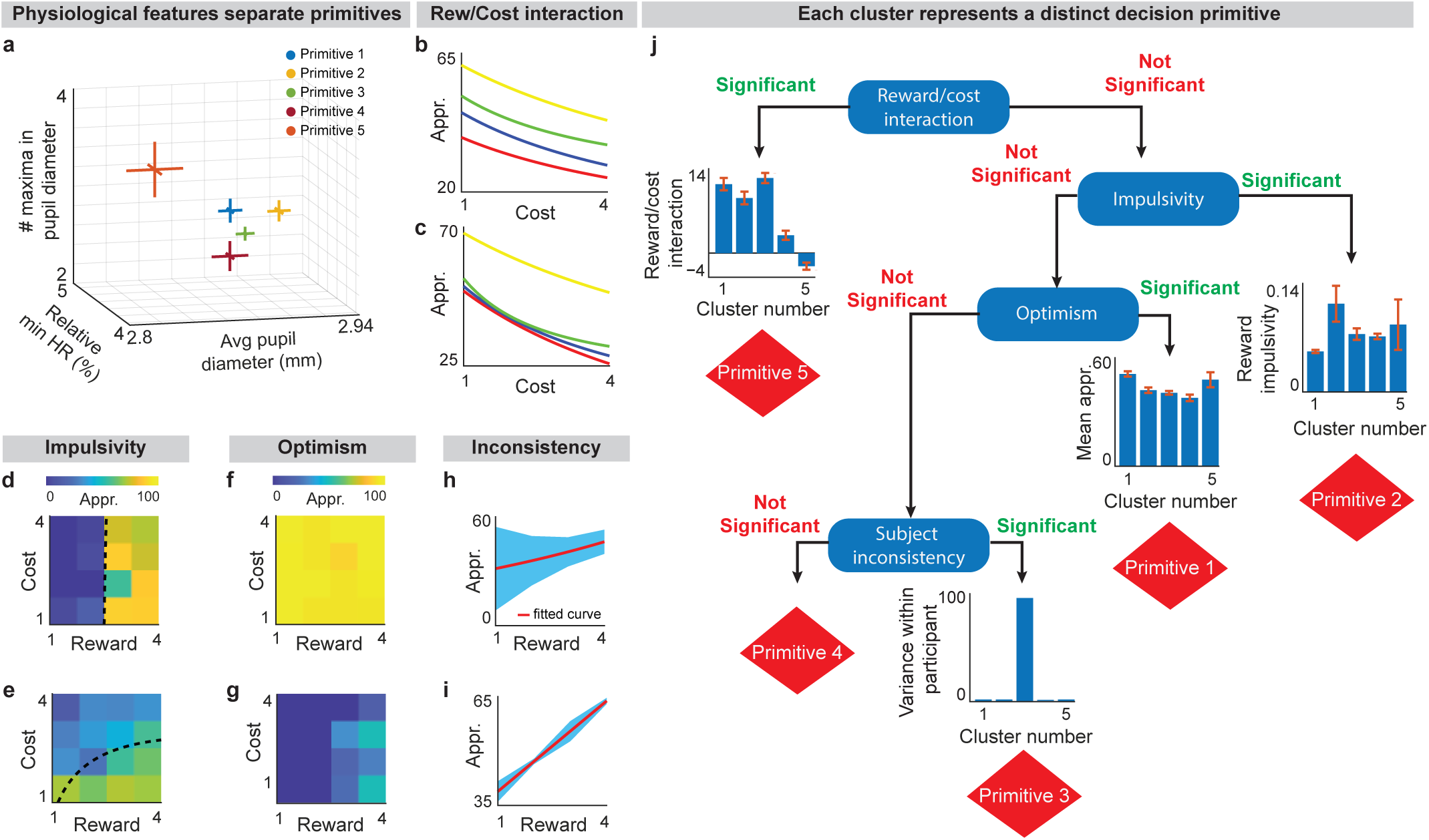
Behavioral strategies define clusters. **a.** Features derived from eye tracking and heart rate data show that each primitive has a unique physiological profile (F(8, 13378) = 8.87, p=3.44×10^-12^, 2-way ANOVA). **b,c.** Examples of high and low “reward/cost interaction”. **d,e.** Examples of high and low “impulsivity”. **f,g.** Examples of “optimism” and “pessimism”. **h,i.** Examples of high and low “inconsistency”. **j.** Each primitive has a representative feature or unique set of features which differentiates it from other primitives. For example, Primitive 5 has low reward/cost interaction while Primitive 3 has high participant inconsistency (reward/cost interaction: F(4, 699) = 13.67, p=9.66×10^-11^ ANOVA, impulsivity: F(4, 699) = 2.5, p=0.036, ANOVA; optimism/pessimism: F(4, 699) = 10.47, p=3.06×10^-8^ ANOVA, inconsistency: p<1×10^-63^ F-test).

We also identified characteristics of decision-making that significantly differentiate primitives from each other. We defined a set of behavioral features: “interaction”, “impulsivity”, “optimism”, ”pessimism”, and “consistency”. We calculated these features for each session completed by a participant, then grouped the features by cluster and calculated summary statistics for each cluster (**Methods; Features in tree diagram**). We found that each primitive corresponded to a unique set of behavioral features. For example, participants mostly consider reward and cost simultaneously when making decisions (**Fig. 2b**), but in one primitive, “interaction”, rewards and costs do not interact – participants do not consider both together when deciding (F(4, 699) = 13.67, p=9.66×10^-11^, ANOVA, **Fig. 2c**). Some primitives show “impulsivity”. Participants that are highly impulsive switch decisions from “no” to “yes” drastically and quickly, (F(4, 699) = 2.5, p=0.036, ANOVA, **Fig. 2d**) while unimpulsive participants change their decisions more gradually (**Fig. 2e**). Other primitives demonstrate “optimism”, a tendency to accept offers with high likelihood. A participant may be optimistic (F(4, 699) = 10.47, p=3.06×10^-8^, ANOVA, **Fig. 2f**) or be “pessimistic” (**Fig. 2g**). Finally, there may be “inconsistency” across a participant’s sessions, denoted by high variation, (p<1×10^-68^, F-tests, **Fig. 2h)** or “consistency”, when participants act with little variation across sessions (**Fig. 2i)**. While primitives are defined without using these features, we find that each primitive can be significantly linked to a different feature or unique combination of features. We use a tree diagram as an example to show how primitives can be separated by combinations of features, though other orderings may yield separations across primitives as well **(Fig. 2j)**.

### Context-specific strategies arise from differential distribution of behavior across primitives

Using an approach that calculates a single average of behavior for each decision-making context, it appears that unique behavioral strategies are used in the four contexts (**Fig. 1g, 3a**). We wanted to understand why these differences in strategy arose, and if they could be related to the primitives identified above. Differences across contexts could arise if each context relied on a different decision-making primitive, but it could also arise if each context utilized a different distribution of behavior across primitives. We conducted several analyses to differentiate between these two hypotheses.

**Fig. 3.**
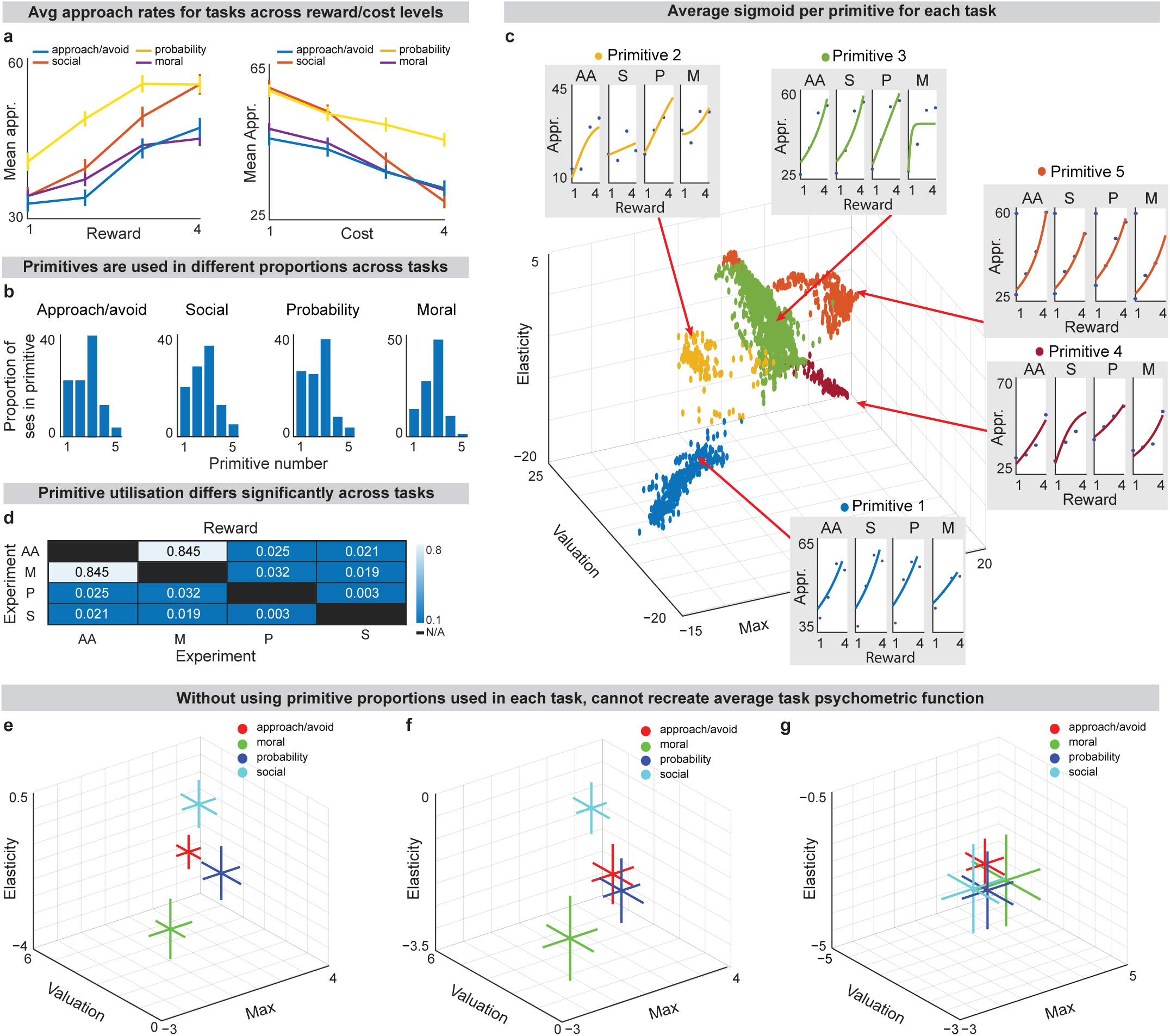
Primitives are used differently across tasks. **a.** Average approach rate uniformly increases with reward and decreases with costs across all 4 tasks, with significant difference between tasks when keeping cost constant and varying reward (F(9, 1760) = 2.11, p=5×10^-6^, 2-way ANOVA) and keeping reward constant and varying cost (F(9, 1760) = 4.6, p=0.02, 2-way ANOVA). **b.** Primitives are used differently across tasks. We abbreviate the tasks with the following: AA=approach-avoid task, S=social task, P=probability task M=moral task. **c.** All primitives appear in every task, with similar psychometric functions within primitives across tasks. **d.** Primitive proportions differ significantly across all tasks, except for moral and approach-avoid tasks, using data split by cost and reward levels (p<0.01, chi-squared, AA n=25 participants, 683 points; S: n=27 participants, 617 points; P: n=29 participants, 623 points; M: n=29 participants, 577 points). This table shows the p-values of chi-squared tests between the proportions used for each task pair. **e.** Plotting the average psychometric function for each task demonstrates that strategies for each task are distinctly different. **f.** This discrete clustering across task types can be replicated after shuffling the psychometric data across tasks using known primitive probabilities. **g.** We shuffle the psychometric data across tasks and recombine without the use of the primitive probabilities for specific tasks and can no longer re-create the same average psychometric functions, unlike in **(f).**

First, for each of the four contexts, we examined the proportion of stories using each behavioral primitive (**Fig. 3b**). We found that each primitive was observed in every context, but the distribution of behavior across the primitives differed significantly across contexts (p<0.01, chi-squared, **Fig. 3c,d, Methods: Task Differences**). Furthermore, using a weighted average of the proportions of primitives multiplied by the shapes of the psychometric functions of each primitive, we reproduced the average psychometric function for each task **(Fig. 3e,f).** When the proportions were randomly shuffled, all tasks collapsed to the same average function **(Fig. 3g, Methods: Task Differences)**. These two analyses show that the distribution of behavior across the primitives is key to defining task differences, and demonstrate that averaging behavior within each of the four decision-making contexts obscures the shared computational strategies for relating cost to reward. The moral task, for example, yielded a noticeably different pattern than the others, with a comparatively sharp drop in use of primitive 5. This suggests near-ubiquitous presence of reward-cost interaction and we speculate that perhaps moral choices bias towards more thoughtful weighing of rewards and costs. The social task, however, shows more use of primitive 2 which relates to impulsivity. Perhaps social aspects of choice are more non-negotiable and past a certain threshold, decision-making becomes less nuanced.

### Behavioral heterogeneity across individuals arises from differential distribution of behavior across primitives

We next sought to determine whether participants exhibited similar preferences for one or more decision-making primitives and to compare those preferences across individuals. The distribution of behavioral strategies varied greatly across individual participants: some participants routinely used one primitive while others were more flexible in primitive use (**Fig. 4a-c, Extended Data Fig. 5a-d**). In some cases, an individual’s behavior is linked across multiple task contexts. We find that the primitive used in one task can be predictive of the primitive used in another using a Bayesian analysis (**Fig. 4d-f, Extended Data Fig. 5e-h; Methods: Conditional Primitive Membership**). In general, we find that primitive 3 is used most often across all subjects and tasks. We speculate that it may be a sort of default strategy while others could reflect momentary aberrations based on other factors such as mood. This speculation may be additionally supported by the physiological profile of primitive 3 which is very central in the physiological feature-space, suggesting it may sit in the “neutral mood” state, as compared to states of high arousal or interest, based on pupil dilation and heart rate (**Fig. 2a).**

**Fig. 4.**
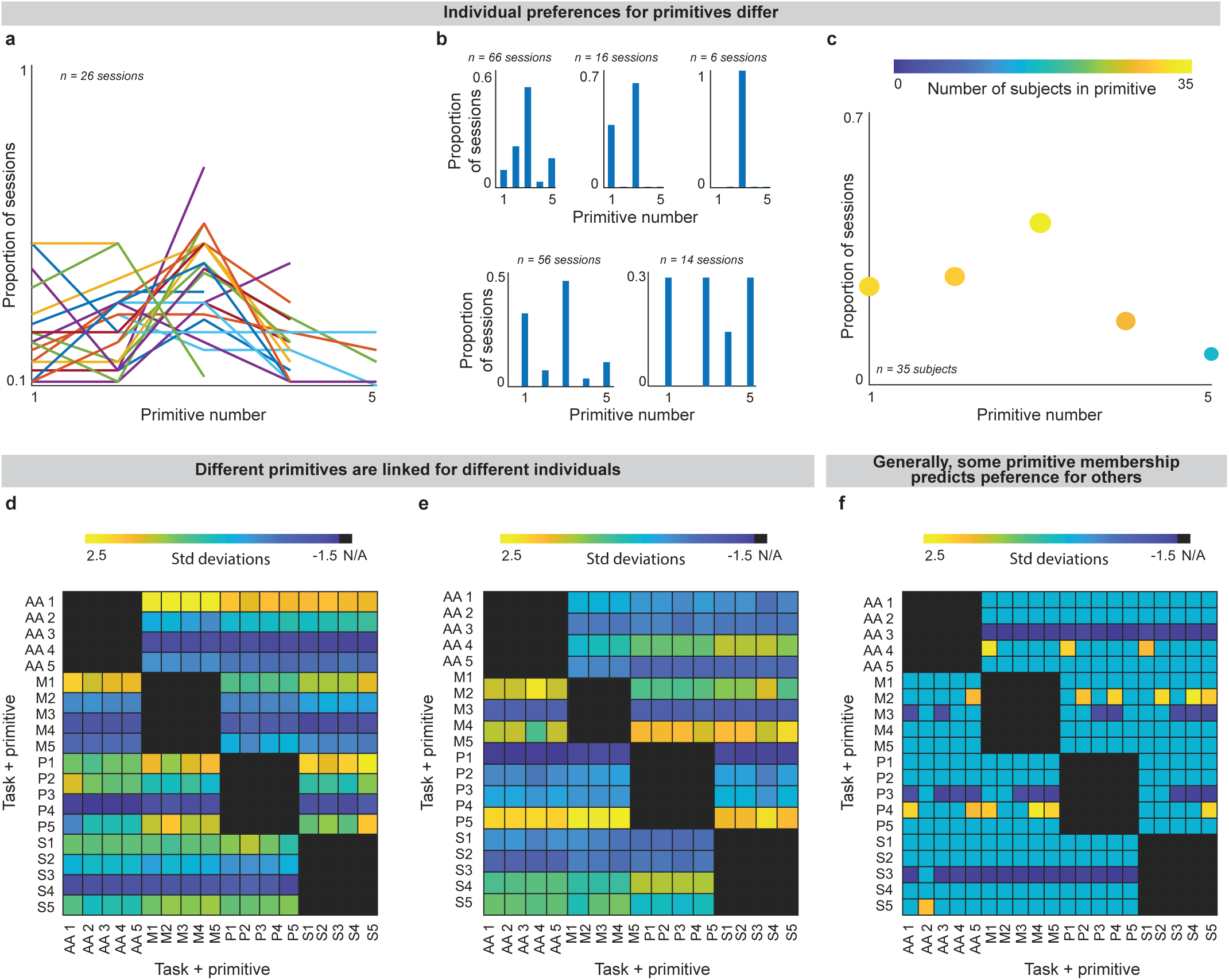
Use of decision-making primitives differs across individuals, and primitives across tasks can be linked. **a.** Different participants exhibit different decision-making strategies across sessions in a task: while some switch between several different primitives, others use a single primitive repeatedly. Each line represents an individual participant completing the approach-avoid task (see **Extended Data Fig. 4a-c** for other tasks). **b.** Examples of individual participant’s use of primitives. Each bar chart represents the proportion of sessions in each primitive for a participant in a given task. **c.** Some primitives are used across more individuals (denoted by color, yellow points have more participants than blue points). In addition, for each individual some primitives are used in more sessions (denoted by size and on the y-axis, n=35 participants). **d-f.** Each square in the heatmap demonstrates the probability of a primitive being used in a task given what primitive was used in another task. Yellow squares indicate a high likelihood of using both primitives while a dark blue square indicates that it is highly unlikely a participant uses both primitives (**Methods: Conditional Primitive Membership).** This indicates that sometimes primitive use is linked across tasks. **(d,e)** Show individual examples, while **(f)** shows the average likelihoods across all subjects.

### Rodents also use cost-benefit decision-making primitives

We wondered if these primitives may exist across species. If primitives also appeared in rat data, this would be additional confirmation of our findings by verifying their existence in a second organism. To compare rats and humans, we used data from a previously developed high-throughput system for measuring cost-benefit decision-making in rats^7^. This system uses a similar set-up by measuring approach rates in rats for differing levels of trade-offs between costs and rewards, both in a baseline state as well as in a food-deprived state. Rats were sensitive to changing rewards and costs, acting in line with neuroeconomic expectations **(Fig. 5a,b).** Most importantly, discrete primitives emerged from the rat data. We again used fuzzy c-means clustering to quantify these distinct clusters (MPC = 0.7061, **Fig. 5c).**

**Fig. 5.**
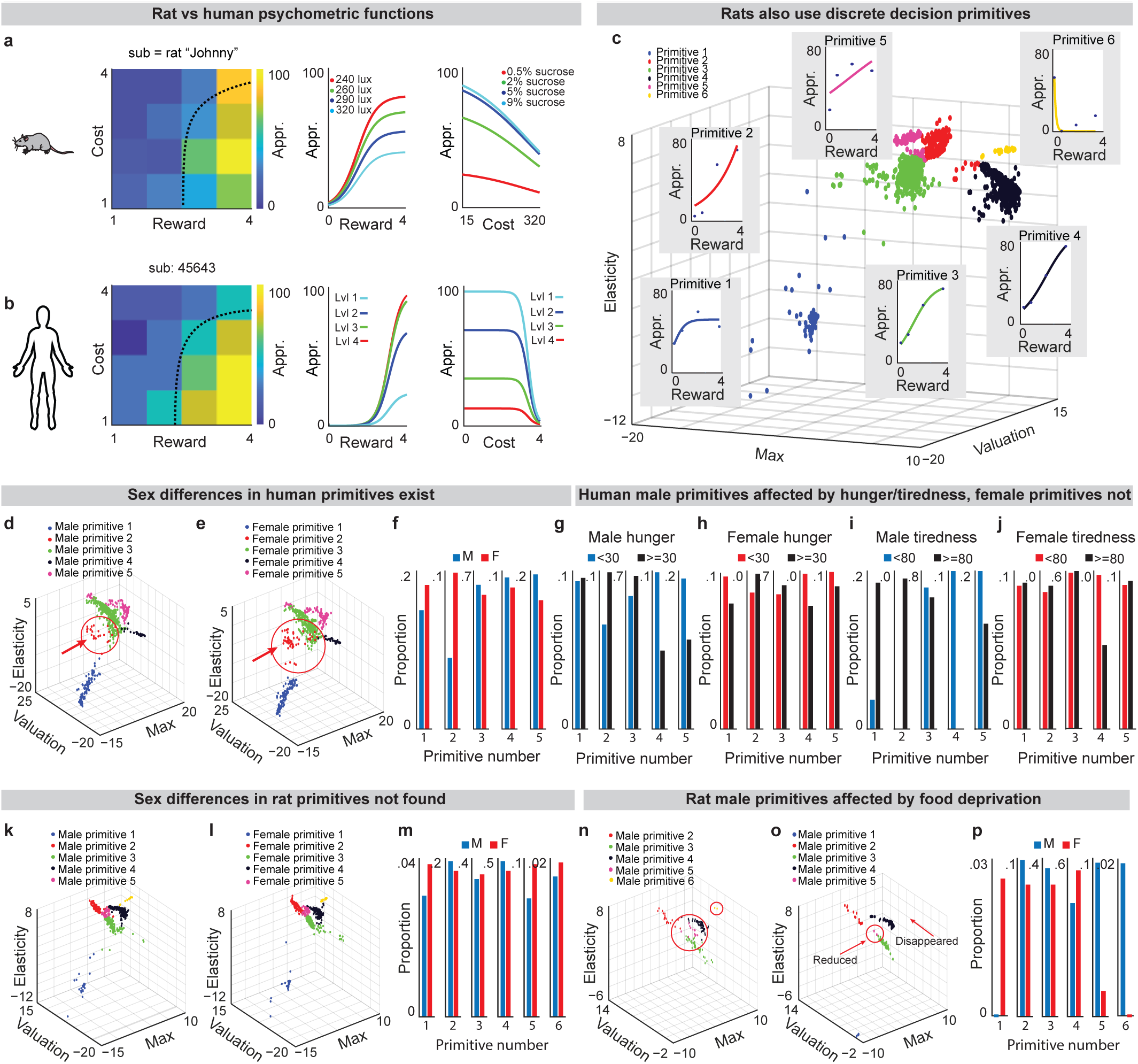
Primitives in rats; sex, hunger, tiredness effects primitives. **a,b.** Example of a rat’s (**a**) and human’s (**b**) decision-making maps (left) and psychometric functions during cost-benefit trade-off decision-making. **c.** Sigmoidal data from rats can be clustered similarly to human data and reveals the existence of discrete behavioral primitives in rats (MPC=0.7061, n=42 rats, 1616 points). **d-f.** Differences in primitives between male (n=11 subjects, 695 points) and female (n=24 subjects, 695 points) humans do exist (p=0.003, chi-squared). Female data was cut to match size of male data set (**Extended Data Fig. 6c,d**). **g,h.** Hunger changes the proportions of primitives used in human males (p=0.0108, threshold=30, chi-squared), but effects were not found in females. **i,j.** Tiredness changes the proportions of primitives used in human males (p=0.024, threshold=80, chi-squared), but effects were not found in females. These effects hold across thresholds as shown in **Extended Data Fig. 6g,h**. **k,l,m.** Differences in primitives between male (n=19 subjects, 751 points) and female (n=23 subjects, 751 points) rats were not found. **n,o,p.** Primitive proportions differ significantly for male rats experiencing food deprivation as compared to ad libitum male rats (p=0.011, chi-squared).

### Internal states shift primitive use

We wanted to examine if these primitives were affected by biological states. Decision-making is often impacted by physiological changes^35^; therefore, we reasoned that the decision primitives we found would be as well. We examined the effects of hunger, and sex on rat and human decision primitives. We found a significant difference between male and female primitive use in humans, suggesting preference for different decision primitives by each sex (p=0.003, chi-squared, **Fig. 5d-f, Extended Data Fig. 6c,d**). In our experiments, we also found sex-specific effects of self-reported hunger and tiredness: the proportions of male decision primitives differ significantly across several thresholds for both hunger and tiredness (hunger: p=0.0108, threshold=30, chi-squared; tiredness: p=0.024, threshold=80, chi-squared) whereas proportions of female decision primitives used do not differ significantly (hunger: p=0.3897, threshold=30, chi-squared; tiredness: p=0.8666, threshold=80, chi-squared; **Fig. 5g-j; Methods: Rat Primitive Analysis, Metadata Collection**). In particular, male primitives 2, 4, and 5 differed greatly with hunger, suggesting higher impulsivity and lower reward-cost interaction when hungry. Tiredness similarly showed increases in use of primitive 2, indicating increase impulsivity, but also showed an increase in use of primitive 1, suggesting higher approach rates as well. This aligns with research that indicates that both tiredness and hunger (generally induced using ghrelin) have been found to increase impulsivity^36,37^. For ad libitum rats, sex difference in rats were not observed (p=0.63406, chi-squared, **Fig. 5k-m**). After the rats underwent challenge like food deprivation, the primitives used by each sex differed significantly (p=0.0256, chi-squared, **Extended Data Fig. 6k-m**). Furthermore, we found sex-specific effects of hunger in rats: males used primitives in different proportions after food deprivation (p=0.011, chi-squared, **Fig. 5n-p**) but females did not (p=0.116, n=12, chi-squared, **Extended Data Fig. 6n-p**). Thus, decision primitives for rats and humans differ with different biological variables such as sex, hunger, and tiredness, which again serves as further confirmation of the validity of these primitives. We additionally examined primitives in humans and rats and found that there were some primitives that were shared across the two species **(Extended Data Fig. 7)**.

### Neural network with natural cortico-striatal-dopaminergic architecture also shows decision primitives

Given increasing interest in using artificial intelligence (AI) tools to supplement human decision-making^38^, we were interested in the ability of artificial intelligence (AI) to reproduce human decision primitives. A feedforward network trained on the full potential space of sigmoidal psychometric functions (**Methods: Construction of AI)** yielded no discrete primitives in psychometric space **(Fig. 6a, Extended Data Fig. 8a)**. We predict that a neural net without restrictions creates every possible strategy instead of filtering to only adaptive/useful strategies that reliably yield good outcomes, unlike rats and humans.

**Fig. 6.**
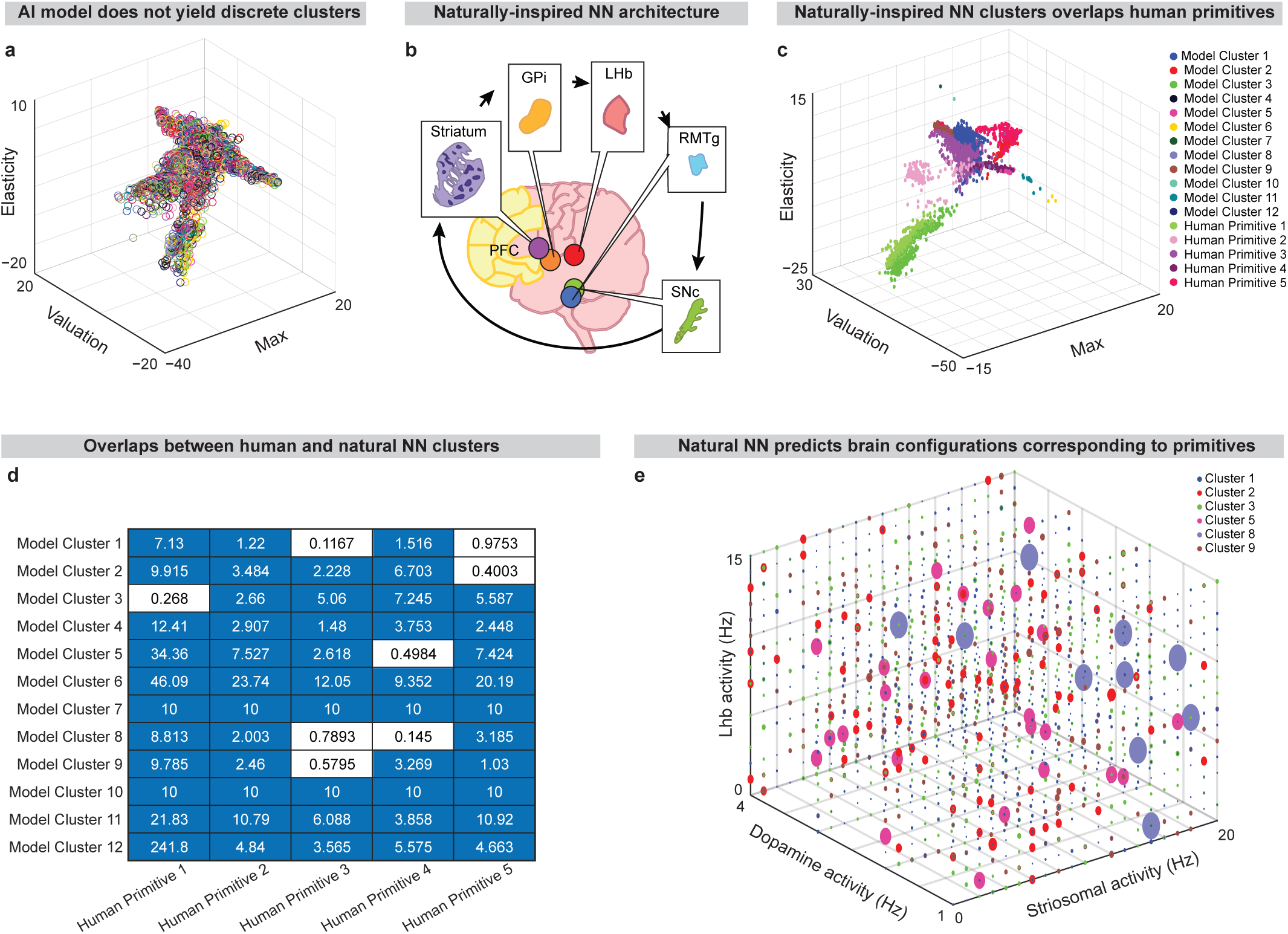
Naturally-inspired network reproduces human clusters while AI does not. **a.** The collection of artificial neural networks trained to different portions of the possible space do not produce discrete clusters (**Extended Data Fig. 8a)**. **b.** Depiction of “naturally-inspired” neural network architecture based on decision-making circuit**. c.** Naturally-inspired NN model produces clusters similar to experimentally found primitives. **d.** Bhattacharya distances between experimentally found human primitives and clusters generated by our naturally inspired neural network show that some clusters overlap. Blue indicates that clusters do not overlap, white indicates that clusters do overlap. **e.** Using our naturally inspired neural network, we predict the circuit activity that corresponds to certain primitives. In some cases, there are many potential configurations that may give rise to the same behavior. Color corresponds to cluster membership and we only show clusters that overlapped in location with human primitives. (**Method: Naturally-inspired Neural Network).**

In other modeling work, we developed a computational model based on the role of the cortico-striatal-dopaminergic circuit in decision-making^39^. We focus on this circuit for several reasons. The cortex calculates information relevant for decision-making^40–43^. Striosomes serve to gate cortical information ahead of dopaminergic decision-making^44^. Dopamine and the lateral habenula (LHb) calculate critically important information related to reward, cost, and reward-prediction error^45–48^, making them interesting regions to model for decision-making. We chose this motif since, in our view, it is necessary for defining the state of decision-making which may be the underlying mechanism of the decision primitives identified here.

The architecture of this network is restricted by real-world connectivity patterns of several key brain regions in the circuit (**Fig. 6b**). We wondered if restrictions such as these were sufficient to reproduce the primitives we found in our human data (**Fig. 1j**). We found that while the NN produced more clusters than in human experiments, several of these clusters overlapped with the human-like decision primitives found (**Fig. 6c; Methods: Naturally-inspired neural network**). Comparing modeled primitives to human primitives, we find that several of them overlap in location using Bhattacharya distance to measure closeness (**Fig. 6d**). This network additionally gives us the ability to predict potential activity of brain regions underlying each commonly employed decision-making primitive (**Fig. 6e**). We identify several patterns of activity in the striatum, of dopaminergic neurons (DA), and Lateral Habenula (LH) that could be validated in humans performing cost-benefit decision-making in a functional magnetic resonance imaging machine in future work (**Extended Data Fig. 8f-j)**. This gives us the potential to find combinations of brain activity that may lead to different decision-making primitives.

## Discussion

Our novel decision-making task framework and tailored analytical approaches enabled us to identify a set of decision-making primitives, which are correlated with physiological responses like heart rate, eye movement, and shift in utilization across different task contexts and internal states. In addition, these primitives were observed in multiple species. Discrete principles have broadly been observed across fields of science, with similar findings in motor movement^22,49,50^, neural patterns of simpler organisms^19^, protein and gene families^51–53^, and even in the quantum states of chemistry and physics^54,55^. The repetition of this pattern suggests the use of discrete organizational principles across the natural world.

Though participants generally follow neuroeconomic expectations in approaching more as reward increases and vice versa for costs, we find that different strategies are used to accomplish this: for example, using increased “impulsivity” or acting with “optimism” and “pessimism”. Changes in strategy may be linked to internal states and the related neuronal activities^56^. Supporting this idea, we observed shifts in decision-making primitive utilization based on hunger, tiredness, and sex. Thus, future research could explore decision-making primitives as a measure for state-dependent variations in traditional neuroeconomics including prospect theory^57^ or utility theory^58^ as well as providing an important lever for understanding neuropsychiatric disorders affected by internal state.

Shifts across decision-making primitives may explain many psychiatric disorder phenomena. Beyond decision-making as a symptom, there are several phenomena where not all individuals show impaired decision-making^59–61^, and that the severity of decision-making symptoms during disorders, can vary across days^62–64^ (for example, when a patient decides to take medication willingly one day, but not the next). In addition, aberrant decision-making may reflect new decision-making primitives that are not typically observed in healthy decision-makers.

We borrowed the term ‘primitives’ from established literature on motor primitives; compositional elements of motor behaviors which were uncovered by decomposing complex motor behaviors^49^. This approach of breaking down complex movements into their components, termed movement ‘syllables’, has recently been applied to rodents to facilitate connection between neural recordings to the correlated behavior/movement^22^; emphasizing the importance of precise characterization of behaviors^65–67^ and robust neural recording systems^68,69^. By examining more fundamental aspects of movement rather than compositions of multiple components, it is easier to identify a one-to-one mapping onto neural activity. Our research, and the resulting decision-making primitives, reflect a similar approach of simplifying decision-making which could potentially increase the accuracy and granularity of simultaneously recorded neural activity.

Further exploration of decision-making primitives could be of interest to various fields including neuroscience, psychiatry/psychology, and even computer science. Decision-making primitives could be useful for linking the activity of specific brain regions or circuits, including but not limited to, the cortex^40^, striatum^40,70,71^, or basal ganglia^72^ to specific decisions, and decision-making related behaviors. Decision-making primitives may also aid in developing or modifying more “naturally-inspired” artificial intelligence systems, including deep learning tools that seek to explain brain function computationally^73^. Our decision-making task framework and all the analytical tools used in this manuscript are software based, thus any lab with a computer could adapt and implement our tools to examine approach-avoid decision-making in different environments.

## Limitations

Our framework for reducing the dimensionality of complex human decision-making is only one of many potential approaches and the primitives that we describe here are not the only strategies guiding decision-making behavior. Our behavioral battery does not explore certain important aspects of decision-making (e.g. temporal discounting), and it is possible that other task features might drive participants to use strategies occupying other parts of the parameter manifold that were not used during our behavioral battery. There could also be uncommon behavioral strategies that were not represented in the psychometric functions we collected here, that may be apparent if psychometric functions were collected for other populations of participants. In addition, the behavioral clusters we identified could be further stratified if we considered additional dimensions (e.g. decision time). Despite these limitations, human psychometric functions represent an appealing approach for dissecting cost-benefit decision-making because it is fundamentally quantitative, and it explores behavior over a wide range of cost and reward values. Further, by normalizing subjective value across participants and contexts, our representation of the decision-making boundary in parameter space is tailored measurement for cost-reward integration. Using these tools, we have been able to identify discrete primitives of decision-making that have exciting implications for both theoretical and practical work in the fields of neuroscience, neuroeconomics, psychiatry, psychology, and computer science.

## Acknowledgements

We greatly appreciate G. Schoenbaum, and Y. Shaham for their constructive criticism and discussions during model development. This project was supported by the NSF/CAREER (#2235858), NIH/NIDA (#R01DA058653), U-RISE T34GM145529 and G-RISE T32GM144919.

## Methods

### Research Ethics Statement

Human data collection for this project received approval for all protocols from the University of Texas at El Paso IRB on June 19, 2024 (FWA No. 00001224).

Animal data collection for this project received approval for all protocols from the University of Texas at El Paso Institutional Animal Care and Use Committee and followed the Guide for Care and Use of Laboratory Animals (IACUC reference number: A-202009-1).

### Demographic Data

We collected data from 35 participants, all members of the University of Texas at El Paso community. Further details of demographic data can be found in Extended Data Table 1. All participants provided written informed consent before the start of the experiment. In all analyses involving sex differences, we trimmed the number of participants to be equal for both groups, as described in the section **Statistical methods of data trimming** below.

This publication is dedicated to open science. From the data collection all the way through to the figure creation and statistics, all processes are done automatically and without subjective human interference. As participants completed the task, the application described below uploaded data automatically to a database. This data without exclusion is submitted under https://doi.org/10.7910/DVN/OZARPL. Run-me codes for each figure, analysis, and statistics can be found in the **Supplemental Methods and Materials Document,** along with instructions for replication.

#### 1. Decision-making application

Depending on context, approaches to decision-making can vary wildly: one reacts very differently to an impending car accident than when choosing what to eat for dinner. To understand decision-making holistically, we developed an application with 4 story-based decision-making tasks to measure the effect of context on decision-making primitives (https://github.com/lrakocev/dec-making-app/blob/raq_dev_utep/app.py).

Our application is set up so that a participant encounters a story, chooses rewards and costs relevant to them within that context, and then is asked to evaluate various cost-reward pairings in context. In a session, a participant first reads a story that gives them a basis from which to consider costs and rewards. The value of costs and rewards are participant dependent: what one participant might value another finds meaningless. This is an ongoing problem in the field: previous experiments have tried many vehicles for administering rewards and costs, including money^74^, points^75^, air puffs^76^. In each of these situations, it is difficult to understand the exact effect of each reward or each cost on the participant and how participants weigh these rewards and costs in their decision-making. To this end, we chose to use stories to deliver different levels of rewards and costs. These can be crafted for different populations of participants and participants themselves can decide what importance these rewards and costs have (https://github.com/lrakocev/dec-making-app/tree/raq_dev_utep/stories/task_types).

In the remainder of the methods, details on the development and deployment of the tasks can be found under **Decision-making application** including physiological tracking found under **Physiological Measurements,** an explanation of the mathematics behind the task can be found in **Mathematical principles to make application quantifiable,** discussion of primitive analysis can be found under **Primitive identification** with additional clustering methods described in **Decision-Map Based Clustering**, details on task and individual differences can be found under **Task Differences** and **Conditional Primitive Membership** respectively, a description of rat-human and metadata analyses can be found under **Rat Primitive Analysis** and efforts involving AI and naturally-inspired NN are detailed under **Modeling Decision Primitives.** The repositories for all code for all analyses are as follows: https://github.com/lrakocev/human_dm, https://github.com/lrakocev/dec-making-app, and https://github.com/lddavila/human_dec_making. For details on any given section or figure panel and related analyses, look to the corresponding section in the Supplemental Methods and Materials.

#### 2. Stories used in decision-making tasks

For each task, we created 24 potential stories that could be presented to the subject. The reasoning behind the number of stories is described above in the **Mathematical principles to make application quantifiable** section. The participant chooses topics which are relevant to them, and from there stories are presented one by one. The story is presented on a screen and gives the participant background before they choose their preferences.

An abbreviated example of a story context would be the following: you are going on a vacation and must take a long flight. What kind of seat would you like to sit in? Once the participant has the necessary context, we present a list of 6 reward and 6 cost preferences which participants rate from 0 to 100 on a sliding scale of how pleasant (or unpleasant) each item is in the given scenario. Participants must rate each item differently and are encouraged to rate items on an absolute scale. Potential rewards might include having a window seat, a row to yourself, sitting next to a friend (**https://github.com/lrakocev/dec-making-app/blob/raq_dev_utep/stories/task_types/approach_avoid/story_2/pref_reward.txt**) while potential costs might include sitting close to a screaming child, in a cramped seat, or having to get up often (**https://github.com/lrakocev/dec-making-app/blob/raq_dev_utep/stories/task_types/approach_avoid/story_2/pref_cost.txt**). Once the participant has completed the reward and cost preference submission, the app chooses 4 of these preferences each (outlined in the **Normalization** methods) and builds 16 questions with them, pairing each cost with each reward. These are presented in a random order, and each time the participant rates their likelihood of accepting the offer from 0 to 100 (0 being never accept, 100 being always accept) on a sliding scale. These offers combine the rewards and costs, for example: sitting next to a friend, but having to get up often (**https://github.com/lrakocev/dec-making-app/blob/raq_dev_utep/stories/task_types/approach_avoid/story_2/questions.txt**). Additional story descriptions and links to all of the stories can be found in https://github.com/lrakocev/dec-making-app/tree/lr_analysis/stories.

#### 3. Mathematical principles to make application quantifiable

##### 3a.#Number of stories + trials

We aimed to make decision-making analyzable. In theory, to best understand a subject’s decision-making, we would examine decisions at every possible combination of rewards and costs with multiple samples of each combination to ensure that we have an accurate understanding of their decision at those levels. However, for each reward-cost combination, you need to present a question and record a response. This is unfortunately time-consuming, and participants start to weary after many responses and begin to respond less accurately.

To ensure accurate data, it is necessary to balance the human element of collecting valid data with the need to collect enough data. We found that to best capture a subject’s decision-making, 8-12 stories are needed. This means that for each participant in each task, we have 8-12 data points for each reward-cost combination. This allows us to correct for variability across an individual’s decision-making. Details on why this number of stories is necessary can be found below in the section **Story number criteria.**

Each story produces 16 responses corresponding to 4 reward levels and 4 cost levels. To read the story and to decide on preferences takes 5-10 minutes, and to respond to 16 questions takes between 15-20. Therefore, each session takes 20-30 minutes to complete in total. Participants usually cannot complete more than 3 stories per session otherwise they will begin to answer without full concentration. Considering this timing and the necessity for 8-12 stories per task, we spread the data collection across 3-4 days per task. Thus, to complete this for all 4 tasks takes a total of 12-16 days.

##### 3b.#Story number criteria

While sometimes fewer stories can yield reasonable results, we find that using 8-12 stories per participant guarantees a successful decision-making map. There are several criteria used for evaluating the quality of a subject’s decision-making map. We expect that approach rate increases with reward and decreases with cost, and that both reward and cost are integrated in decision-making. We use ANOVA to detect the significance of the effects of reward and cost (https://github.com/lrakocev/human_dm/blob/permutations/psychometric%20analysis/psychometric_anova.m). This generally creates a decision-making boundary that is diagonal in shape, with approach being more likely below the boundary and avoid being more likely above (**Extended Data Fig. 2e,f,k,l,q,r,w,x**).

##### 3c.#Normalization of preference ratings

It is crucial to get each individual’s preferences in order to accurately present rewards and costs to them that correspond to their perceived value of the reward and cost. In order to make the levels of reward and cost comparable both across individual subject’s sessions as well as across participants in general, it is necessary to create bins for the reward/cost levels. Essentially, we are trying to compare like-valued rewards and like-valued costs. We choose to create 4 such bins because this is the minimum necessary to fit a sigmoid.

To arrive at these 4 bins, we initially provide 6 preferences for the participant to choose from and then we select 4 to use in the trials. We start with 6 preferences to increase the likelihood of getting 4 equally spaced rewards for each person. For example, if we start with only 4 rewards and 2 out of 4 are meaningless to a subject, then we have no alternatives to choose from. Starting with more preferences allows flexibility. To select preferences, we compare the difference that each rating has with every other rating, then choose the configuration of four out of the six ratings that create the most equal differences between consecutively ranked ratings. This is essential for creating accurate psychometric functions. Reward/cost levels must be similarly spaced, or the meaning of each level is rendered meaningless. For example, a reasonable spread of levels would be: 22, 47, 72, 97 (**Extended Data Fig. 1b-i**) If a participant is choosing between preferences with too similar of a rating, we will not be able to capture the effect of raising the reward/cost since the participant does not experience a tangible increase in the reward/cost (https://github.com/lrakocev/human_dm/blob/permutations/psychometric%20analysis/get_ratings_by_subject.m).

##### 3d.#Granularity of response

We give the participant a sliding scale from 0 to 100 to mark their response instead of measuring discrete responses: if the participant slides the scale to 78%, they are indicating that they are 78% likely to take that offer. This allows us more granularity than a participant responding with a binary yes or no. For example, 8 yes responses and 2 nos would be an approach rate of 80% when using Boolean values, but using numerical values yields a more detailed picture of the approach rate. In addition, the accuracy of the responses increases-it may be difficult for the participant to concretely respond yes or no and a numerical value allows for a more nuanced response. In this way, we make it easier for the participant to respond while gaining more statistical power from their responses.

##### 3e.#Data Organization

When running tasks, often multiple participants would be completing the task at the same time across several different computers. Therefore, it was necessary to coordinate the organization and upload the data to the same database. We use a PostgreSQL database hosted at Utep to upload our data continuously and automatically from the application while the participant is completing the task (https://github.com/lrakocev/dec-making-app/blob/raq_dev_utep/app.py). Upon final collection, all data was uploaded to the Harvard database (https://doi.org/10.7910/DVN/OZARPL).

In the initial session, the participant is given a subject-id number and from then on is referred to in the database only by that id number and is otherwise completely anonymized. The participant metadata is recorded only as reported by the participant (as described in the **Metadata** section). In addition to this metadata, the following information is recorded and uploaded continuously: current task type, story number, trial number, reward and cost level, reward and cost ratings, approach rates, eye-tracking data, heart-rate data, story relevance. Participants use the same ID number to link their data across several sessions. Further information can be found in the **Database User Guide** section in the Supplemental Materials and Methods.

##### 3f.#Physiological Measurements

We use the Tobii Pro Spark eye tracker and the OpenANT and pulse monitor to track eye movement and heart rate respectively. We measure heart rate and eye movement on a trial level and continuously write this data to the database while the participant is completing their trials. In order to make this cooperate with our decision-making app, we wrote custom libraries to interface between the externally bought hardware and our application software (https://github.com/lrakocev/dec-making-app/blob/raq_dev_utep/eyetracker_lib.py, https://github.com/lrakocev/dec-making-app/blob/raq_dev_utep/heartrate_lib.py). Further information about the app (including the heart rate and eye tracker) and data collection can be found in **Decision-Making App User Guide** in the Supplemental Materials and Methods.

#### 4. Primitive Identification

##### 4a.#Psychometric function fitting

We fit a psychometric function to each grid of approach rates corresponding to cost-reward offers within a single story/session. We averaged the approach rates across cost levels so that each grid is distilled into 4 approach rates corresponding to the 4 reward levels. Using Matlab’s built-in fitting function (https://www.mathworks.com/help/curvefit/fit.html), we fit the following functions to the approach rate data: 2 parameter sigmoid, 3 parameter sigmoid, 4 parameter sigmoid, parabola, line. Since the fitting function uses a random starting point, we run the fit function up to 20 times to ensure that the best fit is not missed due to an incorrect starting point. We then compare the R-squared value from each of these fit types. If the fit is above the threshold of 0.4, that function is assigned to the data with preference given in the following order: 3 param sigmoid, 4 param sigmoid, 2 param sigmoid, parabola, line. (https://github.com/lrakocev/human_dm/blob/permutations/clustering/fit%20psychometric%20functions/sigmoid_analysis_updated.m). Ultimately, we use only the sigmoidal functions (72% of all data). We include 35 participants in our analysis, totaling 693 sessions across all subjects. (https://github.com/lrakocev/human_dm/blob/permutations/clustering/fit%20psychometric%20functions/cluster_runme.m).

We take the parameters of the fitted sigmoidal psychometric function to use as features describing the overall strategy of the session. The parameters we use, “max”, “valuation”, and “elasticity”, are defined as the following:

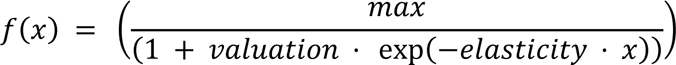

In the case of a 2-parameter sigmoid, max is set to 1. When plotting these parameters, to see the small differences between them we use a log-scale. (https://github.com/lrakocev/human_dm/blob/permutations/clustering/create%20clusters/getTable.m).

##### 4b.#Decision-Map Based Clustering

We wanted to see if discrete clusters in decision-making behavior could be found using another method. Our goal was to capture the decision-making maps in function form so that we could find patterns among these maps. We used a 2D sigmoid function which could be fit to any map and would describe the “decision boundary” between the participant approaching more and avoiding more. The function we fit to was:

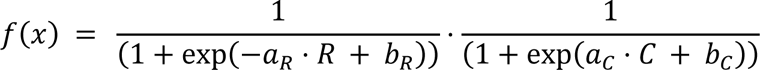

where a_R and a_C are reward and cost elasticity respectively, and b_R and b_C are reward and cost valuation respectively. This can be fit directly onto each session, represented by a 4×4 grid of approach rate data across changing reward and cost. We again took the parameters of this function and identified clusters when plotting these parameters. We chose to plot a_R, b_R, and b_C in our 3D representation, though all combination of the parameters clustered (https://github.com/lrakocev/human_dm/blob/permutations/clustering/alternative%20clustering/alt_clustering_runme.m).

##### 4c.#Identifying Clusters

We tried multiple built-in Matlab methods of identifying clusters in our data including spectral clustering, gaussian mixture models, and fuzzy c-means clustering. We found that each of these methods produces similar results, therefore we chose fuzzy c-means clustering^77^ because it is computationally efficient and provides a probability for each point belonging to a cluster https://www.mathworks.com/help/fuzzy/fcm.html, https://github.com/lrakocev/human_dm/blob/permutations/clustering/create%20clusters/runme.m). In order to determine the number of clusters and the goodness of fit, we use modified partition coefficient (MPC) method^78^ (https://github.com/lddavila/human_dec_making/blob/main/Utility%20Functions/calculate_mpc.m).

After identifying the clusters, we verify that they significantly differ from each other in location, in their means, and in their variances. To compare cluster locations, we use Bhattacharya distance on each parameter. To compare cluster means, we use a two sample t-test (https://www.mathworks.com/help/stats/ttest2.html) across clusters for each parameter. To compare variances, we use two-sample F-test (https://www.mathworks.com/help/stats/vartest2.html) across clusters for each parameter.

##### 4d.#Average psychometric functions per primitive

The average psychometric function of a primitive s calculated by collecting all the initial approach-rate data for the points in that cluster, then averaging all this data for each reward level and using our initial fitting method to fit a new psychometric function to these averaged approach rates. (https://github.com/lrakocev/human_dm/blob/permutations/clustering/per%20cluster%20analysis/per_cluster_runme.m).

##### 4e.#Features in tree diagram

To understand the meaning of each cluster, we calculated a set of features over all data and grouped them by cluster to see if there were significant differences between clusters. We calculated reward/cost interaction, impulsivity, mean approach rate, and internal participant variance. To determine significance, we used ANOVA for differences in means across groups, while for variance related measures, we used F-tests comparing each pair and reporting the largest p-value from these pairs.

For reward/cost “interaction”, if a participant weighs both cost and reward, then their level of approach should be overall higher for a higher reward or for a lower cost. We calculated this feature by taking the average approach rate at each level of reward respectively, then finding the median difference between average levels for the first three levels of reward. “Impulsivity” can be examined by the rate at which a participant changes their approach rate based on changes in cost/reward: sudden jumps in approach are impulsive while gradual changes are not. We calculated impulsivity as the slope of the best-fit line through approach rates. We determine “inconsistency” by examining the variance in the functions used by a participant across all their sessions. This was calculated by comparing all psychometric functions used by that participant to their average psychometric function and finding the variance. “Optimism” is calculated by calculating the mean approach rate for each session (https://github.com/lrakocev/human_dm/blob/permutations/primitive%20building/prims_run_me.m).

##### 4f.#Physiological Features

From our physiological measurements (see above **Methods: Physiological measurements),** we calculated several physiological features across all the data and then grouped these features based on primitive (https://github.com/lrakocev/human_dm/blob/permutations/physio/run_corr_to_clusters.m). The eye-tracking features included: average pupil diameter, number of local maxima in pupil diameter, number of local minima in pupil diameter (https://github.com/lrakocev/dec-making-app/blob/lr_dev_0817/eye_tracker_analysis.py). The heart-rate features included: relative maximum heart rate, relative minimum heart rate, and direction of change in heart rate through the trial (https://github.com/lrakocev/dec-making-app/blob/lr_dev_0817/heart_rate_analysis.py). Measurement is collected on a trial-by-trial basis, so for each session there are 16 trials whose data are averaged. To calculate the number of local maxima/minima in pupil diameter, we first applied a Savitzky-Golay filter (https://docs.scipy.org/doc/scipy/reference/generated/scipy.signal.savgol_filter.html) to smooth the data then found the extrema of the smoothed data using built-in scipy function (https://docs.scipy.org/doc/scipy/reference/generated/scipy.signal.argrelextrema.html).

To calculate the relative maximum/minimum of the heart rate data, we took the highest and lowest point in the trial, then found the difference from the average heart rate of the trial in percentage terms. We found these in relative terms since average heart rate differs for each individual and cannot be directly compared. We calculated direction of change in heart rate by comparing when the maximum HR occurred relative to the minimum HR in the trial. If the maximum occurred before the minimum, the direction decreased and had a value of -1, and vice versa if the maximum occurred after the minimum. If there was no change, direction has a value of 0.

To compare the significance of the features on a per-primitive basis, we used a multi-way ANOVA.

##### 4g.#Features using Psychometric Functions

Beyond the features described in the **Features in tree diagram** section above (“interaction”, “impulsivity”, “optimism”, “inconsistency”), we used several other features to compare primitives (**Extended Data Fig. 3B, 3C, 7C, 7D)**. “Valuation” and “elasticity” refer to the parameters of the psychometric functions as described in the **Psychometric function fitting** section above. Unlike when plotting these parameters, in this case we use the raw values rather than putting them in a log-scale. To calculate “variance in cluster”, we found the average mean squared error of each psychometric function to the average psychometric function of the primitive. To calculate “session variance”, we calculated the variance of all of the choices made in each session (for each reward-cost combination, the participant gives a value from 0-100 for how likely they are to take the offer), then averaged these variances for all the sessions within a cluster. Further details can be found in **Supplemental Materials and Methods: Extended Data Fig. 7C, 7D**, https://github.com/lrakocev/human_dm/blob/permutations/primitive%20building/rat_v_human_runme.m).

#### 5. Task Differences

Based on the unique proportions of points per primitive, each task has a unique average psychometric function which describes the average strategy for that task across all participants. We extracted coefficients from this function to show us where the task exists in psychometric space (https://github.com/lddavila/human_dec_making/blob/main/runMe.m).

#### 6. Conditional Primitive Membership

We calculated the conditional probability of a participant having points in primitive X from task 1 given their membership in primitive Y from task 2 (**Fig. 4d-f)** using Bayes’ rule. To make a statistical claim about this data, we performed bootstrapping by randomly selecting different subsets of the data using Matlab’s builtin function bootstrp (https://www.mathworks.com/help/stats/bootstrp.html) and performing the same analysis repeatedly. We then averaged the value for each primitive pair across these different iterations and subsets of data. We found the average likelihood of belonging to one primitive based on membership of another in another task, then found the standard deviation of each specific pair’s likelihood compared to the mean likelihood. We then used standard deviation to denote significance. (https://github.com/lrakocev/human_dm/blob/permutations/clustering/chaining/chain_behavior_runme.m).

#### 7. Rat Primitive Analysis

For our rat-human comparisons, we use rat data collected from a maze-like open field system created in RECORD (https://www.nature.com/articles/s42003-024-06489-8?utm_source=rct_congratemailt&utm_medium=email&utm_campaign=oa_20240706&utm_content=10.1038/s42003-024-06489-8). In this system, rats are given offers with 4 levels of reward and 4 levels of cost. This creates the capability of building psychometric functions along a reward or cost axis per animal. We include 42 rats in our analysis, totaling 1616 sessions split across 19 male and 23 female rats. In addition to the baseline experiments, experiments involving food-deprived rats were also run, involving 22 rats.

In the rat data, not every reward and cost level are paired together as in the human data for each rat, therefore we cannot consistently produce decision-making maps for each rat session as we do for the human sessions. To compare the rat and human data, we must use psychometric functions along either a reward or cost axis, rather than the 2D sigmoidal functions that include both. Therefore, we use our initial clustering approach as described in the **Clustering** section above to analyze the rat data (**Fig.5a-c**). (https://github.com/lddavila/human_dec_making/blob/main/runMe.m).

##### 7a.#Metadata collection

At the beginning of each session, we collect demographic information as well as additional metadata including sex, gender identity, age, race, ethnicity, hunger, tiredness, pain, stress, menstruation, weight, relationship status, sexual orientation, education, story relevance. For metadata entries, participants rate on a scale of 0 to 100 how much they are experiencing the attribute where 0 would be not at all and 100 would be maximally. (https://github.com/lrakocev/dec-making-app/blob/raq_dev_utep/app.py).

Because we are working with human participants and not lab animals, we cannot control the experience of the participant - i.e., we cannot deprive a participant of food to ensure hunger. Therefore, when a person reports their hunger, they are reporting on a perceived experience which differs from person to person. Reporting is subjective which makes finding a threshold that splits the data into “has” and “has not” difficult. Because of this fact, in our analyses that involve metadata (such as hunger and tiredness), we compare across a range of thresholds and check the validity of the claim across all these thresholds **(Fig. 5d-j, Extended Data Fig. 6e-h, https://github.com/lddavila/human_dec_making/blob/main/session_cost_clustering_figures_threshold_0.m**).

##### 7b.#Statistical methods of data trimming

As noted above, we cannot control the state of a participant before the experiment begins. As such, there may be many participants that come hungry and few that come having eaten for example. Because of these varying outside states, the number of data points in two groups being compared could differ vastly. In analyses that involve comparing different groups, we use trimming methods to equalize the number of data points in each of the groups. To do so, we use built-in Matlab functions, datasample (https://www.mathworks.com/help/stats/datasample.html) and randstream (https://www.mathworks.com/help/matlab/ref/randstream.html), to sample the data without replacement.

##### 7c.#Similar Primitives between Rats and Humans

We wanted to see if any decision primitives were shared among rats and humans. To do so, we compared both the locations of primitives as well as behavioral features exhibited in each cluster. We used Euclidean distance to identify primitives that were overlapping in location. We used a variety of behavioral features, which is described in more detail in the **Supplemental Methods and Materials Document**. To identify similar primitives, we first narrowed the potential pairings to primitives that had overlapping location. Then, we compared the features that each primitive exhibited. We counted both features that were exhibited significantly and were not exhibited significantly in our matching. For example, rat primitive 1 and human primitive 1 both do not exhibit high valuation, but they both do exhibit high reward interaction, therefore they match on both accounts (**Extended Data Fig. 7b-d, https://github.com/lrakocev/human_dm/blob/permutations/primitive%20building/rat_v_human_runme.m**).

#### 8. Modeling Decision Primitives 8a. Identifying Sigmoidal Space

The clusters are situated in the “sigmoidal domain” so we sought to visualize this space. We start with the full range of transformed coefficients appearing among the human clusters as plotted, and create all possible combinations of these coefficients. We then apply the reverse transformation that is applied for plotting to get the raw coefficients. From there, we use these as the coefficients in a sigmoid function for a 2, 3, and 4 parameter sigmoid and use x-values 1-4 to produce corresponding y-values. Then, we filter this set based on these y-values to the set of plausible values for our experiment by checking that values are within 0 and 100 (the range of possible approach rates participants could choose from). (hhttps://github.com/lrakocev/human_dm/blob/permutations/ai%20primitives/create_sigmoidal_space.m)

Once we have identified the sigmoidal space, we perform the same steps on this space as in the **Psychometric function fitting** section above (https://github.com/lrakocev/human_dm/blob/permutations/ai%20primitives/testing_bias_sig_fit.m). For each set of y-values, we fit a psychometric function using the built-in Matlab fit function (https://www.mathworks.com/help/curvefit/fit.html). We then get the parameters of this fitted function and plot these parameters using the log-scale as described in our initial plotting methods above (**Extended Data Fig. 8a**). While in theory all combinations of parameters can yield a sigmoidal function, only certain combinations of parameters can yield psychometric functions that could be plausibly achieved using our experiment because the outputs of these functions must be between 0 and 100.

We use the built-in Matlab functions boundary (https://www.mathworks.com/help/matlab/ref/boundary.html) and trisurf (https://www.mathworks.com/help/matlab/ref/trisurf.html) to identify the borders and volume of the final space and experimentally found primitives. We find that the experimentally found human primitives occupy 9.7% of the possible space (**Extended Data Fig. 8b**).

##### 8b.#Comparison of Primitives to Sigmoidal Space

We sought to compare the sigmoidal space described above to the human primitives found experimentally. We wanted to identify where the experimental primitives were located in this space, as well as compare the density of the primitives (i.e. density of the cluster) relative to the space. To do so, we applied fuzzy c-means clustering on the sigmoidal space using a high designated number of clusters in order to over-cluster the space. This would give some sense of the densities of different portions of the space and by over-clustering, we could isolate more granular pieces of space (**Extended Data Fig. 8c**). Then, we found which theoretical clusters overlapped with the human primitives using Bhattacharya distance. We calculated the normalized density of these overlapping clusters and compared them to the normalized densities of the primitives. In the case that the primitives overlapped with multiple theoretical clusters, or vice versa, we summed the normalized densities together. We found that the normalized densities of the primitives differed from the normalized density of the pieces of sigmoidal space they overlapped (**Extended Data Fig. 8d, https://github.com/lrakocev/human_dm/blob/permutations/ai%20primitives/space_vs_hum_runme.m**).

##### 8c.#Construction of AI

We chose to use the simplest possible model to see if this model could produce all psychometric functions within the sigmoidal space described above. We use the built-in Matlab model feedforwardnet (https://www.mathworks.com/help/deeplearning/ref/feedforwardnet.html), which consists of a series of layers, the first of which has a connection from the network input and the last of which produces the network’s output. In this case, we specify one hidden layer with 10 neurons. The input of this network is a 100×4 array where each row is a set of four reward values 1-4, and the output of the network is a 100×4 array of corresponding approach rates at each of those reward values. We use the default training method of “trainlm” and use the built-in training and validation criteria to check convergence of training.

We start with the full possible space of data as described in the **Identifying Sigmoidal space** section, then split this data into 12 evenly sized sections corresponding to the number of models we want to use. Within each section, we randomly select a set of 100 rows of data with 100 sets of approach rates (which come in groups of 4, corresponding to the 4 reward levels). We then train a model on this set of data and yield a new set of 100 rows of approach rates. We fit a sigmoid to these approach rates as before using the built-in Matlab fitting function and record the coefficients for this data. We repeat this for 5 iterations per section. (https://github.com/lrakocev/human_dm/blob/permutations/ai%20primitives/ai_runme.m) We found that these models fill all the space described above.

##### 8d.#Naturally-inspired neural network

We use our previously developed naturally-inspired neural network^79^ that provides a potential explanation for the linkages between several brain regions and their unique roles in decision-making. This toolbox, focusing on the cortico-striatal-dopaminergic circuit, explains mechanistically how each of the neuronal elements of this circuit contribute to how choices are made, and how the activity of this circuit can create decision-making maps (https://github.com/dirkbeck/DM_space_model/tree/main).

This model is inherently restricted since it is based on the biology of the brain (**Fig. 6b, Extended Data Fig. 8e**). Only certain connections exist among brain regions unlike in an AI network where any connection can exist. In addition, the range of values for activity in a region is limited. For example, striosomes have an average firing rate range between 0 to 20 hz - values significantly above that would be biologically unlikely^80^.

We use this model to find behavior based on different brain configurations. To this end, we simulate the full possible range of brain configurations based on the likely physiological values for each region and plug these combinations into the model. These values are derived from a review of the existing literature (https://github.com/lrakocev/human_dm/blob/permutations/modeling/dirk_model_sim.m).

Once these values have been plugged in, the model produces a behavioral output and approach rate. Using these approach rates at different reward levels, we can fit a sigmoid to this data in the same way as in the experimental data. From here, we plot these sigmoidal coefficients and find a finite number of clusters, some of which include clusters seen in human data (**Fig. 6c**, https://github.com/lrakocev/human_dm/blob/permutations/ai%20primitives/dirk_helper_runme.m). We compare these clusters using Bhattacharya distance and see that there are overlaps between the two sets (**Fig.6d**, https://github.com/lrakocev/human_dm/blob/permutations/ai%20primitives/dirk_helper_runme.m). Additional clusters produced by the model which do not overlap with experimental clusters may be found in future experiments or may not exist experimentally.

Since the model produces behavior based on possible brain configurations, this model gives us the ability to predict potential configurations of activity that might lead to different behaviors. For example, we link levels of activity in striosomes, dopaminergic neurons, and in the lateral habenula to our experimentally found primitives (https://github.com/lrakocev/human_dm/blob/permutations/ai%20primitives/clusters_to_configs.m). We could confirm these levels of activity with fMRI in future experiments.

## Extended Data Tables

**Extended Data Table 1:**
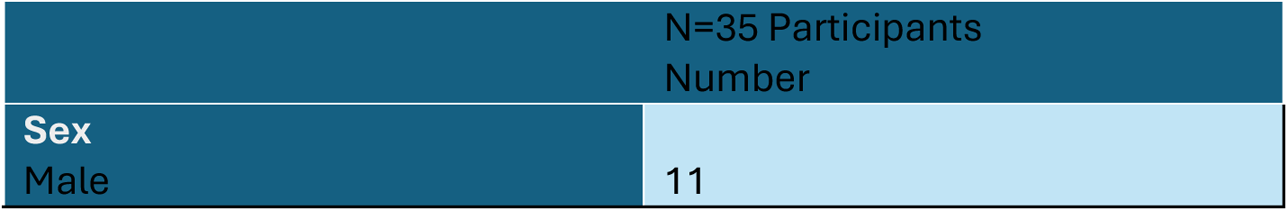
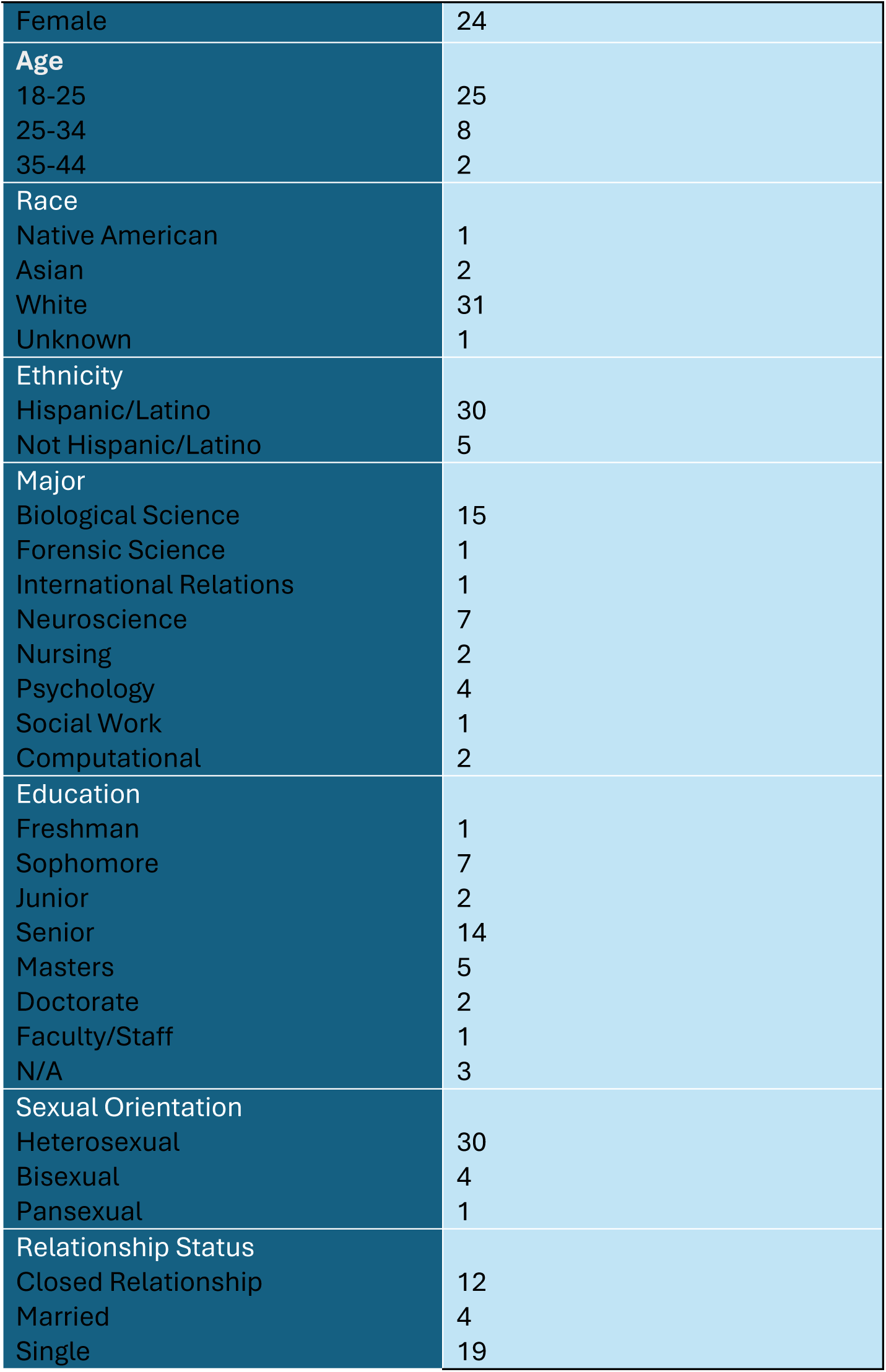
Participant Metadata.

## Notes

### Competing Interest Statement

The authors have declared no competing interest.

## References

1. Colautti, L., Antonietti, A. & Iannello, P. Executive Functions in Decision Making under Ambiguity and Risk in Healthy Adults: A Scoping Review Adopting the Hot and Cold Executive Functions Perspective. Brain Sciences 12, 1335 (2022).

2. Platt, M. L. & Huettel, S. A. Risky business: the neuroeconomics of decision making under uncertainty. Nat Neurosci 11, 398–403 (2008).

3. Trimmer, P. C. et al. Decision-making under uncertainty: biases and Bayesians. Anim Cogn 14, 465–476 (2011).

4. Berry, A. S., Jagust, W. J. & Hsu, M. Age-related variability in decision-making: Insights from neurochemistry. Cogn Affect Behav Neurosci 19, 415–434 (2019).

5. Orsini, C. A., Truckenbrod, L. M. & Wheeler, A.-R. Regulation of sex differences in risk-based decision making by gonadal hormones: Insights from rodent models. Behavioural Processes 200, 104663 (2022).

6. Aupperle, R. L. & Paulus, M. P. Neural systems underlying approach and avoidance in anxiety disorders. Dialogues in clinical neuroscience 12, 517–31 (2010).

7. Ibáñez Alcalá, R. J., et al. RECORD, a high-throughput, customizable system that unveils behavioral strategies leveraged by rodents during foraging-like decision-making. Commun Biol 7, 822 (2024).

8. Dexter, J. P., Prabakaran, S. & Gunawardena, J. A Complex Hierarchy of Avoidance Behaviors in a Single-Cell Eukaryote. Current Biology 29, 4323–4329.e2 (2019).

9. Adams, G. K., Watson, K. K., Pearson, J. & Platt, M. L. Neuroethology of decision-making. Current Opinion in Neurobiology 22, 982–989 (2012).

10. Wansink, B. & Sobal, J. Mindless Eating: The 200 Daily Food Decisions We Overlook. Environment and Behavior 39, 106–123 (2007).

11. Endrass, T. & Ullsperger, M. Decision-making as transdiagnostic construct for mental health research. Neuron 109, 1912–1914 (2021).

12. Goschke, T. Dysfunctions of decision-making and cognitive control as transdiagnostic mechanisms of mental disorders: advances, gaps, and needs in current research. Int J Methods Psych Res 23, 41–57 (2014).

13. Milkman, K. L., Chugh, D. & Bazerman, M. H. How Can Decision Making Be Improved? Perspect Psychol Sci 4, 379–383 (2009).

14. Lăzăroiu, G., Pera, A., Ștefănescu-Mihăilă, R. O., Mircică, N. & Negurită, O. Can Neuroscience Assist Us in Constructing Better Patterns of Economic Decision-Making? Front. Behav. Neurosci. 11, 188 (2017).

15. Rahman, S., Sahakian, B. J., Cardinal, R. N., Rogers, R. D. & Robbins, T. W. Decision making and neuropsychiatry. Trends in Cognitive Sciences 5, 271–277 (2001).

16. Zhu, Y. et al. A Newly Designed Mobile-Based Computerized Cognitive Addiction Therapy App for the Improvement of Cognition Impairments and Risk Decision Making in Methamphetamine Use Disorder: Randomized Controlled Trial. JMIR Mhealth Uhealth 6, e10292 (2018).

17. Gao, P. & Ganguli, S. On simplicity and complexity in the brave new world of large-scale neuroscience. Current Opinion in Neurobiology 32, 148–155 (2015).

18. Safaie, M. et al. Preserved neural dynamics across animals performing similar behaviour. Nature 623, 765–771 (2023).

19. Hamood, A. W. & Marder, E. Animal-to-Animal Variability in Neuromodulation and Circuit Function. Cold Spring Harb Symp Quant Biol 7G, 21–28 (2014).

20. Sridhar, G., et al. Uncovering multiscale structure in the variability of larval zebrafish navigation. *ArXiv* arXiv:2405.17143v1 (2024).

21. Scholz, M., Dinner, A. R., Levine, E. & Biron, D. Stochastic feeding dynamics arise from the need for information and energy. Proceedings of the National Academy of Sciences 114, 9261–9266 (2017).

22. Weinreb, C. et al. Keypoint-MoSeq: parsing behavior by linking point tracking to pose dynamics. Nat Methods 21, 1329–1339 (2024).

23. Golowasch, J., Goldman, M. S., Abbott, L. F. & Marder, E. Failure of averaging in the construction of a conductance-based neuron model. J Neurophysiol 87, 1129–1131 (2002).

24. Charpentier, C. J. et al. Heterogeneity in strategy use during arbitration between experiential and observational learning. Nat Commun 15, 4436 (2024).

25. Lynall, M.-E. & McIntosh, A. M. The Heterogeneity of Depression. AJP 180, 703–704 (2023).

26. Lombardo, M. V., Lai, M.-C. & Baron-Cohen, S. Big data approaches to decomposing heterogeneity across the autism spectrum. Mol Psychiatry 24, 1435–1450 (2019).

27. Ironside, M. et al. Approach-avoidance conflict in major depression: Congruent neural findings in human and non-human primates. Biological psychiatry 87, 399 (2019).

28. Morris, L.-A. et al. Decision cost hypersensitivity underlies Huntington’s disease apathy. Brain awae296 (2024) doi:10.1093/brain/awae296.

29. Morris, L. S. et al. Dissociating self-generated volition from externally-generated motivation. PLoS One 15, e0232949 (2020).

30. Hoeks, B. & Levelt, W. J. M. Pupillary dilation as a measure of attention: a quantitative system analysis. Behavior Research Methods, Instruments, & Computers 25, 16–26 (1993).

31. Van Den Brink, R. L., Murphy, P. R.& Nieuwenhuis, S. Pupil Diameter Tracks Lapses of Attention. PLoS ONE 11, e0165274 (2016).

32. Joshi, S. & Gold, J. I. Pupil Size as a Window on Neural Substrates of Cognition. Trends in Cognitive Sciences 24, 466–480 (2020).

33. Sforza, E., Jouny, C. & Ibanez, V. Cardiac activation during arousal in humans: further evidence for hierarchy in the arousal response. Clinical Neurophysiology 111, 1611– 1619 (2000).

34. Wascher, C. A. F. Heart rate as a measure of emotional arousal in evolutionary biology. Phil. Trans. R. Soc. B 376, 20200479 (2021).

35. Danziger, S., Levav, J. & Avnaim-Pesso, L. Extraneous factors in judicial decisions. Proceedings of the National Academy of Sciences 108, 6889–6892 (2011).

36. Cedernaes, J. et al. Increased impulsivity in response to food cues after sleep loss in healthy young men. Obesity (Silver Spring) 22, 1786–1791 (2014).

37. Anderberg, R. H. et al. The Stomach-Derived Hormone Ghrelin Increases Impulsive Behavior. Neuropsychopharmacol 41, 1199–1209 (2016).

38. Hill, K. & Duc, H. D. T. I Took a ‘Decision Holiday’ and Put A.I. in Charge of My Life. The New York Times (2024).

39. Beck, D. W. et al. Model of a striatal circuit exploring biological mechanisms underlying decision-making during normal and disordered states. Preprint at 10.1101/2024.07.29.605535 (2024).

40. Yun, M. et al. Distinct roles of the orbitofrontal cortex, ventral striatum, and dopamine neurons in counterfactual thinking of decision outcomes. Sci. Adv. 9, eadh2831 (2023).

41. Rushworth, M. F. S., Noonan, M. P., Boorman, E. D., Walton, M. E. & Behrens, T. E. Frontal cortex and reward-guided learning and decision-making. Neuron 70, 1054– 1069 (2011).

42. Krawczyk, D. C. Contributions of the prefrontal cortex to the neural basis of human decision making. Neuroscience & Biobehavioral Reviews 26, 631–664 (2002).

43. Walton, M. E., Croxson, P. L., Behrens, T. E. J., Kennerley, S. W. & Rushworth, M. F. S. Adaptive decision making and value in the anterior cingulate cortex. NeuroImage 36, T142–T154 (2007).

44. Graybiel, A. M. & Matsushima, A. Striosomes and Matrisomes: Scaffolds for Dynamic Coupling of Volition and Action. Annu. Rev. Neurosci. 46, 359–380 (2023).

45. Stopper, C. M. & Floresco, S. B. What’s better for me? Fundamental role for lateral habenula in promoting subjective decision biases. Nat Neurosci 17, 33–35 (2014).

46. Bromberg-Martin, E. S. & Hikosaka, O. Lateral habenula neurons signal errors in the prediction of reward information. Nat Neurosci 14, 1209–1216 (2011).

47. Bromberg-Martin, E. S., Matsumoto, M. & Hikosaka, O. Dopamine in Motivational Control: Rewarding, Aversive, and Alerting. Neuron 68, 815–834 (2010).

48. Eshel, N., Tian, J., Bukwich, M. & Uchida, N. Dopamine neurons share common response function for reward prediction error. Nat Neurosci 19, 479–486 (2016).

49. Giszter, S. F. Motor primitives—new data and future questions. Current Opinion in Neurobiology 33, 156–165 (2015).

50. Latash, M. L. On Primitives in Motor Control. Motor Control 24, 318–346 (2020).

51. Penel, S. et al. Databases of homologous gene families for comparative genomics. BMC Bioinformatics 10, S3 (2009).

52. Milo, R. et al. Network Motifs: Simple Building Blocks of Complex Networks. Science 2G8, 824–827 (2002).

53. Caetano-Anollés, G. & Caetano-Anollés, D. Universal Sharing Patterns in Proteomes and Evolution of Protein Fold Architecture and Life. J Mol Evol 60, 484–498 (2005).

54. Kibler, M. R. From the Mendeleev periodic table to particle physics and back to the periodic table. Found Chem 9, 221–234 (2007).

55. Pereyra, P. Fundamentals of Quantum Physics: Textbook for Students of Science and Engineering. (Springer Berlin Heidelberg, Berlin, Heidelberg, 2012). doi:10.1007/978-3-642-29378-8.

56. Andalman, A. S. et al. Neuronal Dynamics Regulating Brain and Behavioral State Transitions. Cell 177, 970–985.e20 (2019).

57. Kahneman, D. & Tversky, A. Prospect Theory: An Analysis of Decision under Risk. Econometrica 263–291 (1979).

58. Glimcher, P. W., Dorris, M. C. & Bayer, H. M. Physiological utility theory and the neuroeconomics of choice. Games Econ Behav 52, 213–256 (2005).

59. Friborg, O., Hjemdal, O., Martinussen, M. & Rosenvinge, J. H. Empirical Support for Resilience as More than the Counterpart and Absence of Vulnerability and Symptoms of Mental Disorder. Journal of Individual Differences 30, 138–151 (2009).

60. Macrì, S., Zoratto, F. & Laviola, G. Early-stress regulates resilience, vulnerability and experimental validity in laboratory rodents through mother–offspring hormonal transfer. Neuroscience & Biobehavioral Reviews 35, 1534–1543 (2011).

61. Bonanno, G. A. & Mancini, A. D. The Human Capacity to Thrive in the Face of Potential Trauma. Pediatrics 121, 369–375 (2008).

62. Chun, C. A. The Expression of Posttraumatic Stress Symptoms in Daily Life: A Review of Experience Sampling Methodology and Daily Diary Studies. J Psychopathol Behav Assess 38, 406–420 (2016).

63. Eisenlohr-Moul, T. A., Peters, J. R., Chamberlain, K. D. & Rodriguez, M. A. Weekly Fluctuations in Nonjudging Predict Borderline Personality Disorder Feature Expression in Women. J Psychopathol Behav Assess 38, 149–157 (2016).

64. Wright, A. G. C. & Simms, L. J. Stability and fluctuation of personality disorder features in daily life. Journal of Abnormal Psychology 125, 641–656 (2016).

65. Tillmann, J. F., Hsu, A. I., Schwarz, M. K. & Yttri, E. A. A-SOiD, an active-learning platform for expert-guided, data-efficient discovery of behavior. Nat Methods 21, 703– 711 (2024).

66. Pereira, T. D. et al. SLEAP: A deep learning system for multi-animal pose tracking. Nat Methods 1G, 486–495 (2022).

67. Datta, S. R., Anderson, D. J., Branson, K., Perona, P. & Leifer, A. Computational Neuroethology: A Call to Action. Neuron 104, 11–24 (2019).

68. Trevathan, J. K. et al. Calcium imaging in freely moving mice during electrical stimulation of deep brain structures. J. Neural Eng. 18, 026008 (2021).

69. Clark, J. J. et al. Chronic microsensors for longitudinal, subsecond dopamine detection in behaving animals. Nat Methods 7, 126–129 (2010).

70. Kim, H., Sul, J. H., Huh, N., Lee, D. & Jung, M. W. Role of Striatum in Updating Values of Chosen Actions. J. Neurosci. 29, 14701–14712 (2009).

71. Holly, E. N., Díaz-Hernández, E. & Fuccillo, M. V. A blueprint for examining striatal control of cognition. Trends in Neurosciences 45, 649–650 (2022).

72. Hikosaka, O., Kim, H. F., Yasuda, M. & Yamamoto, S. Basal Ganglia Circuits for Reward Value–Guided Behavior. Annu. Rev. Neurosci. 37, 289–306 (2014).

73. Marblestone, A. H., Wayne, G. & Kording, K. P. Toward an Integration of Deep Learning and Neuroscience. Front. Comput. Neurosci. 10, (2016).

74. Festjens, A., Bruyneel, S., Diecidue, E. & Dewitte, S. Time-based versus money-based decision making under risk: An experimental investigation. Journal of Economic Psychology 50, 52–72 (2015).

75. Van Den Bos, R., Homberg, J. & De Visser, L. A critical review of sex differences in decision-making tasks: Focus on the Iowa Gambling Task. Behavioural Brain Research 238, 95–108 (2013).

76. Sierra-Mercado, D. et al. Decision making in avoidance–reward conflict: a paradigm for non-human primates and humans. Brain Struct Funct 220, 2509–2517 (2015).

77. Sebiskveradze, D. et al. Automation of an algorithm based on fuzzy clustering for analyzing tumoral heterogeneity in human skin carcinoma tissue sections. Laboratory Investigation G1, 799–811 (2011).

78. Wang, W. & Zhang, Y. On fuzzy cluster validity indices. Fuzzy Sets and Systems 158, 2095–2117 (2007).

79. Friedman, A., et al. A decision-space model explains context-specific decision-making. Preprint at 10.21203/rs.3.rs-5499511/v1 (2024).

80. Evans, R. C. et al. Functional Dissection of Basal Ganglia Inhibitory Inputs onto Substantia Nigra Dopaminergic Neurons. Cell Reports 32, 108156 (2020).

