## Supplementary Information for "Computational Primitives for Cost-Benefit Decision-Making"

#### Extended Fig. 1

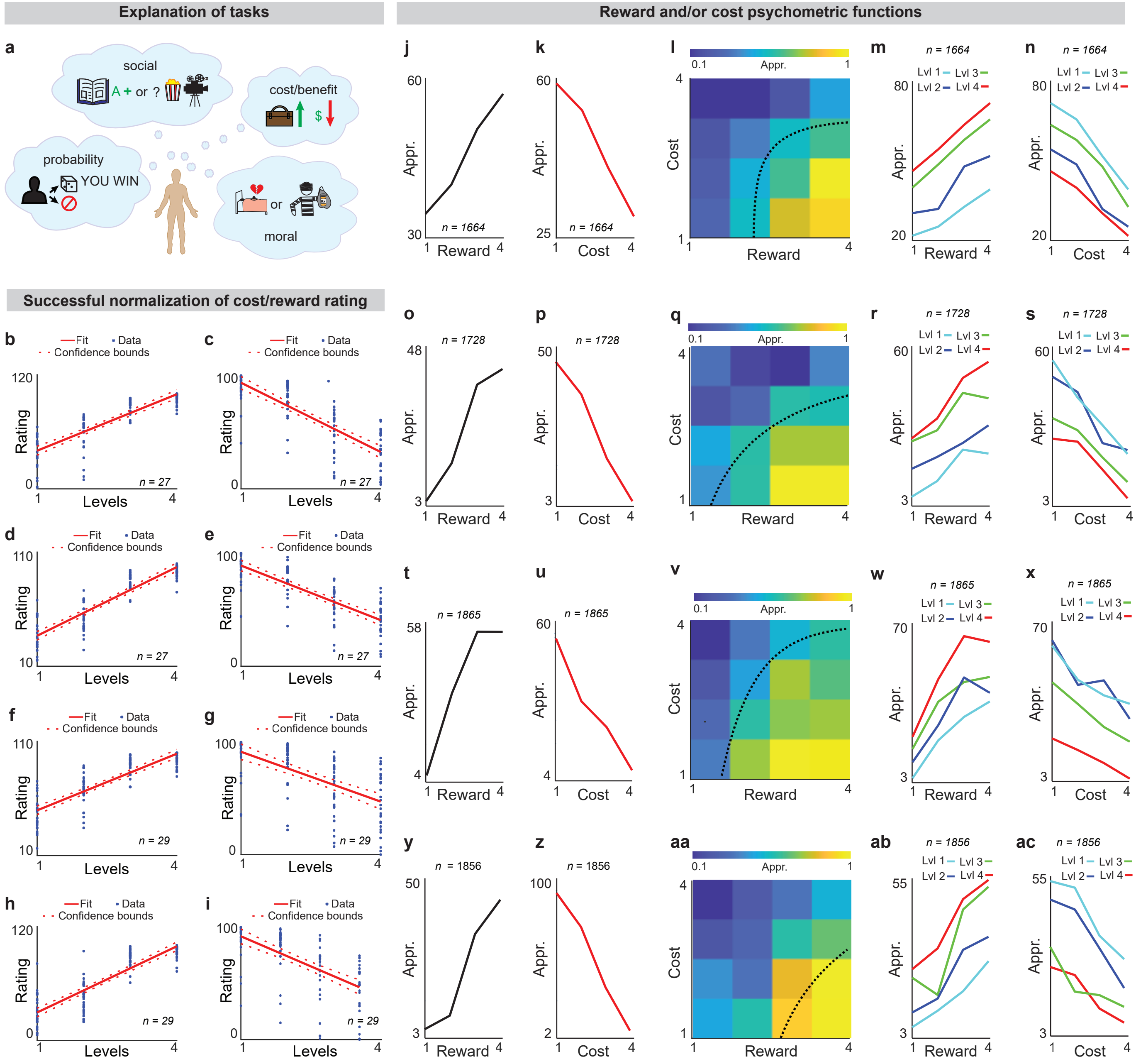

*Figure legend start on next page*

**Extended Data Fig. 1 | Task battery validation and average task performance**

**a.** Explanation of tasks (**Methods: Decision-making application). b-i**. Calibration of participant reward preferences and cost preferences. We perform a linear regression between the reward/cost levels and the respective point-value ratings given by the participants and report R-squared (**Methods: Normalization of preference ratings).** Each dot is an individual subject. The left column represents reward, and the right column represents cost. Approach-avoid (**b**) R-squared=0.7, (**c**) R-squared=0.63, social task (**d**) R-squared=0.79, (**e**) R-squared=.60, probability (**f**) R-squared=0.699, (**g**) R-squared0=.35, moral (**h**) R-squared=0.76, (**i**) R-squared=0.39. **j-n.** Psychometric analysis indicates that in the approach-avoid task, approach rate increases with reward and decreases with cost, and that participants consider both in their decision-making. Each participant contributes one data point and we average across participants (n=26 participants in approach-avoid task). Comparing approach rates at different reward and cost levels using ANOVA, we found significant differences for approach rates across cost levels ($F(3, 1660) = 205.24$, $p=3.24 \times 10^{-113}$) and across reward levels ($F(3, 1660) = 108.09$, $p=6.35 \times 10^{-64}$), as well as when holding reward and cost levels constant. **o-s.** Similar to j-n, but for the social task (cost: $F(3,1724) = 120.49$, $p=7.62 \times 10^{-71}$, reward: $F(3, 1724) = 67.14$, $p=4.61 \times 10^{-41}$, n=27 participants in social task). **t-x.** Similar to j-n, but for the probability task (cost: $F(3\ 1852) = 37.45$, $p=1.79 \times 10^{-23}$, reward: $F(3, 1852) = 76.48$, $p=1.24 \times 10^{-46}$, n=29 participants in probabilistic task). **y-ac.** Similar to j-n, but for the moral task (cost: $F(3, 1852) = 39.83$, $p=6.13 \times 10^{-25}$, reward: $F(3, 1852) = 70.21$, $p=5.26 \times 10^{-43}$, n=29 participants in moral task).

#### Extended Fig. 2

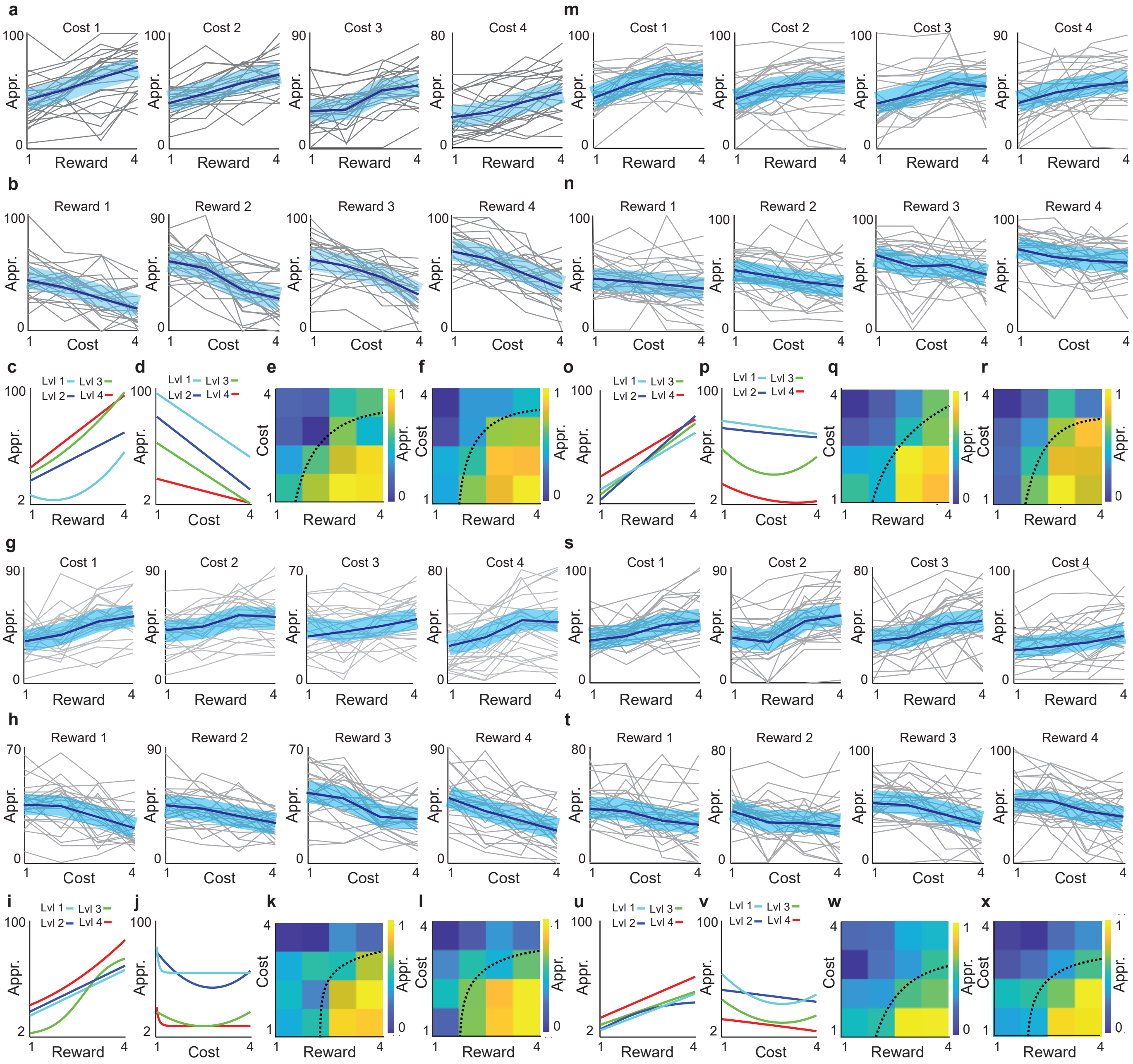

**Extended Data Fig. 2 | Approach rates increase with reward, decrease with cost**

**a,b.** As reward levels increase, the participant approaches more. As cost levels increase, the participant approaches less. The four figures represent different cost levels. Each gray line represents a participant while the bolded blue line represents the average of all subjects. **c,d.** Example of four psychometric functions across reward levels and cost levels for approach avoid task. **e,f.** Example of decision-making maps for the approach avoid task (n=26 participants in approach avoid task). **g-l.** Similar to a-f but for the social task (n=27 participants in social task) **m-r**. Similar to a-f but for the probability task (n=29 participants in probabilistic task) **s-x.** Similar to a-f but for the moral task (n=29 participants in moral task).

#### Extended Fig. 3

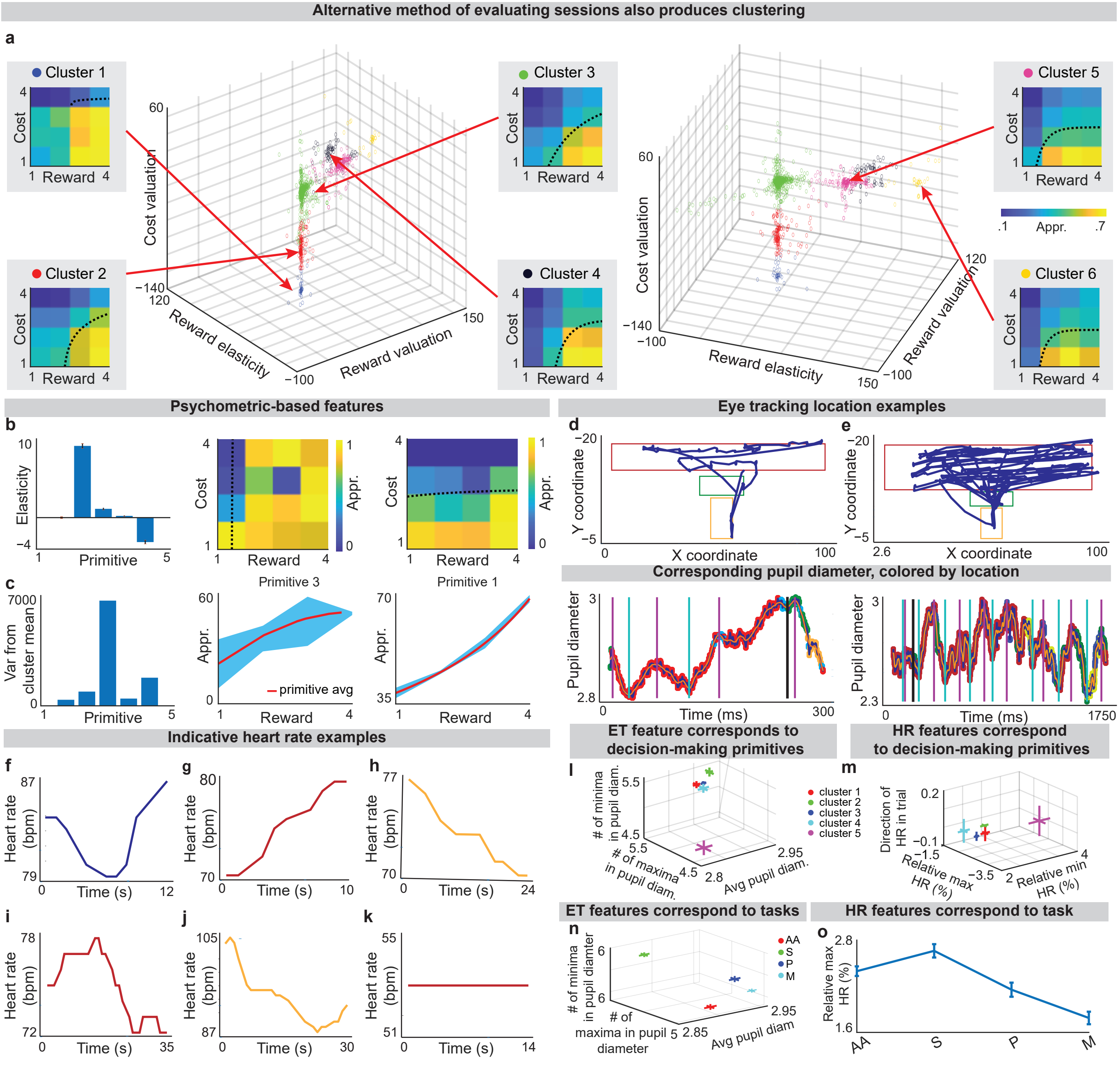

*Figure legend start on next page*

**Extended Data Fig. 3 | Physiological features and correlations to primitives and tasks**

**a.** Using a different method where we fit functions describing the boundary between approach and avoid preference on a decision-making map, we also achieve discrete clusters of functions. Log-scale was not used with this method **(Methods: Decision-Map Based Clustering). b,c.** Features derived from psychometric functions, elasticity and within-primitive variance, also highlight the differences between primitives. Elasticity is a parameter of a subject's psychometric function, while variation from cluster mean is calculated by finding the internal variance of psychometric functions within a cluster **(Methods: Features using Psychometric Functions)**. **d,e.** Eye-tracking data for a single trial shows how a participant's gaze moved over the course of the trial, as well as the diameter of their pupils throughout the task. The boxes on the positional data indicate the location of the question text (red), the approach-rate slider (green), and the submit button (yellow). These positions are also indicated on the pupil diameter data by the corresponding colors. Two distinct trials are shown with examples of the extracted features, including: average pupil diameter, local maxima in pupil diameter, local minima in pupil diameter, the reading period, and the decision-making period. We use the Tobii Pro Spark eye tracker to track eye movement **(Methods: Physiological Measurement, Physiological Features). f-k.** Heart-rate (HR) data for a single trial shows how a subject's beats per minute (bpm) changes over the course of the trial. Individual examples are selected to show extracted features including: biggest dip from the average HR, biggest increase from the average HR, and directionality (+1 if heart rate increased over the course of the trial or -1 if it decreased). We use the OpenANT pulse monitor to track heart rate **(Methods: Physiological Measurement, Physiological Features). l.** Primitives can be differentiated by the average pupil diameter using a one-way ANOVA ($F(4,5227) = 6.41$, $p=3.87\times10^{-5}$, primitive 1 n=960 trials, primitive 2 n=1152 trials, primitive 3 n=2480 trials, primitive 4 n=480 trials, primitive 5 n=160 trials). Here we plot average pupil diameter along with number of extrema (minima and maxima) in average pupil diameter to show the separation between primitives graphically. **m.** Primitives can be differentiated by the maximum heartrate relative to the average HR for each trial using a one-way ANOVA ($F(4,2870) = 5.07$, $p=0.0005$, primitive 1 n=446 trials, primitive 2 n=704 trials, primitive 3 n=1345 trials, primitive 4 n=240 trials, primitive 5 n=140 trials). Here we plot relative maximum HR, relative minimum HR, and direction of HR trend through trial. **n.** Tasks can be differentiated by the following eye-tracking features: number of local minima in the pupil diameter data, number of local maxima in the pupil diameter, and the average pupil diameter using a 2-way ANOVA correcting for interactions. We use the following abbreviations for tasks: AA = approach avoid task, S = social task, P = probability task, M = moral task. ($F(6, 15684) = 31.29$, $p=8.27\times10^{-73}$, AA: n=1520 trials, S: n=1858 trials, P: n=1296 trials, M: n=1232 trials). **o.** Tasks can be differentiated by the maximum HR relative to the average HR for each trial using a one-way ANOVA ($F(3,2871) = 15.83$, $p=3.27\times10^{-10}$, AA: n=952 trials, S: n=821 trials, P: n=608 trials, M: n=494 trials).

#### Extended Fig. 4

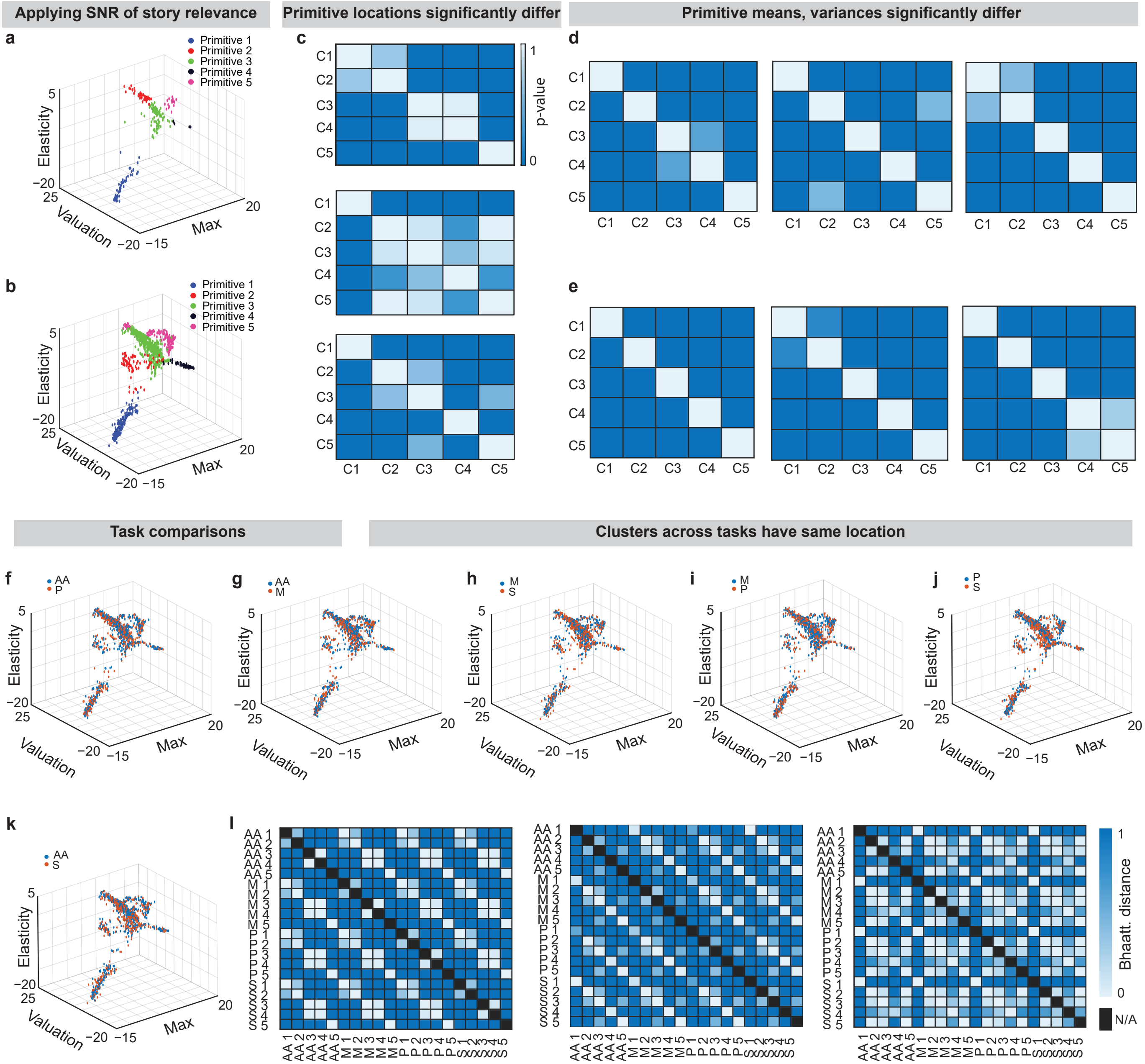

**Extended Data Fig. 4 | Comparisons of primitives across tasks**

**a.** Not all stories are relevant to each subject, therefore we ask participants to rate story relevance to understand the story's importance to that subject. Relevance is rated from 0-100 where 0 is not relevant at all and 100 is maximally relevant. The signal-to-noise (SNR) ratio of story relevance gives a tool to select particularly important stories for each subject. **b.** We apply the same SNR of story relevance with threshold SNR=70 to the data from Fig. 3c. This illustrates sessions where subjects were completing stories that were most relevant to them. We can use this to examine if strategies change depending on the interest level of the subject. **c.** Bhattacharya distances between the primitives for each feature show that the primitives differ significantly from each other in location. Blue indicates that locations differ significantly, while white indicates they do not (n=35 subjects, 2500 sessions, **Methods: Identifying Clusters**). **d.** Primitive means differ significantly for each feature (n=35 subjects, 2500 sessions). **e.** Primitive variances differ significantly for each feature (n=35 subjects, 2500 sessions)**. f-k.** Task comparison for approach avoid vs prob, moral vs social, prob vs social, appr avoid vs moral, moral vs prob, appr avoid vs social. We use the following abbreviations for tasks: AA = approach avoid task, S = social task, P = probability task, M = moral task. (AA: n=25 subjects, 683 sessions, S: n=27 subjects, 617 sessions, P: n=29 subjects, 623 sessions, M: n=29 subjects, 577 sessions). **l.** Bhaattacharyya distance is zero (denoted by the color white) for the same primitives across four tasks, meaning that these primitives overlap. The dimensions considered are max, valuation, and elasticity. The x-axes are task-primitive pairs and the y-axes are the same task-primitive pairs**.**

#### Extended Fig. 5

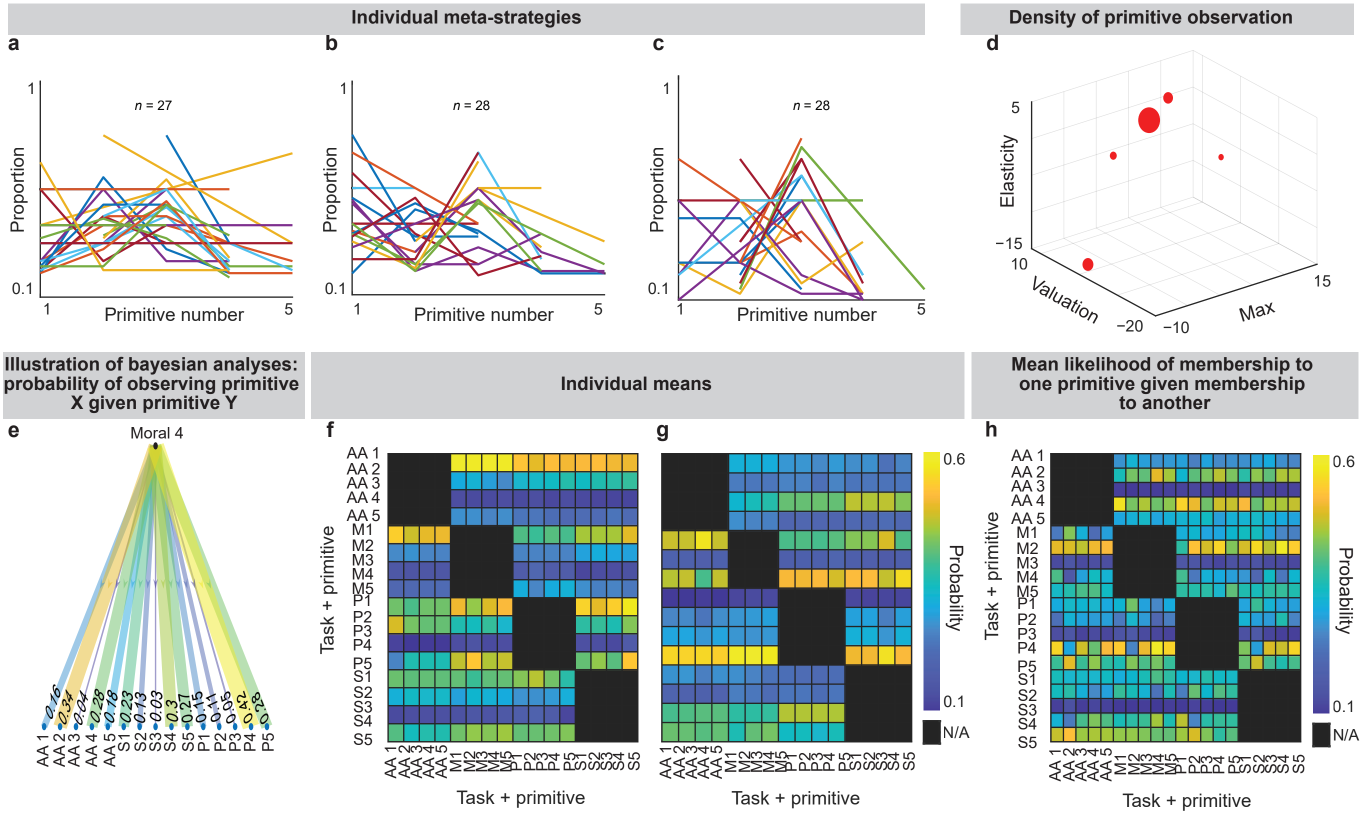

**Extended Data Fig. 5 | Use of decision primitives across individuals and relations between primitive use across tasks**

**a-c.** Same as **Fig. 4a** but for social, probability, and moral tasks respectively. **d.** The space of possible primitives is sparse and primitives are discrete. The location of each point indicates a primitive location, and the size of the point indicates how often the primitive was used relative to all primitives. **e.** Illustration of Bayesian analysis: probability of observing primitive X given primitive Y, in this case observing any primitive given moral primitive 4. Yellow corresponds to a higher likelihood of using a primitive in a task based on the primitive used in a separate task and blue corresponds to a lower likelihood of using a primitive in a task based on the primitive used in a separate task (**Methods: Conditional Primitive Membership). f-h.** We use Bayes rule to find the conditional probability of a participant using primitive X in a task, given that they've used primitive Y in another task. To examine statistical significance of these relationships, we use bootstrapping to take samples of the whole data and calculate the conditional probability for each sample. Then, we find the average conditional probability across all samples and pairs and compare the average probability for individual pairs to the average of all pairs. In **Fig. 4d-f,** we show the standard deviations of difference between the population mean and each individual mean, and in this figure we show the means themselves **(Methods: Conditional Primitive Membership). (f,g)** correspond to the same individuals as in **Fig. 4d,e** while **(h)** shows their average behavior across all individuals.

#### Extended Fig. 6

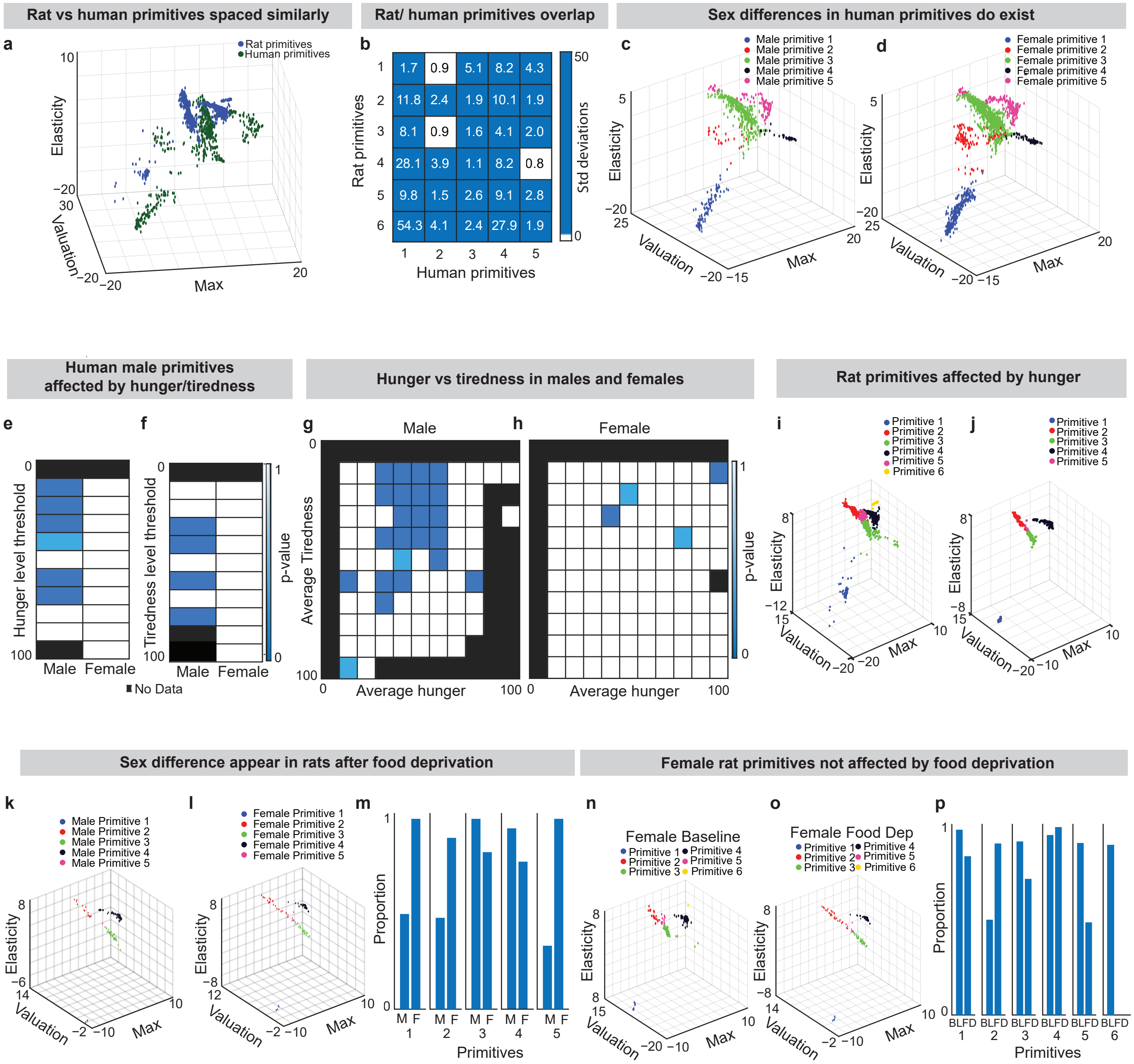

*Figure legend start on next page*

**Extended Data Fig. 6 | Sex differences in rat and human primitives**

**a.** Rat primitives and human primitives are both discrete. **b.** Mean Bhattacharya distances across primitives between each rat and human primitive. Rat primitives 1 and 3 overlap with human primitive 2, and rat primitive 4 overlaps with human primitive 5. **c,d**. Differences in primitives between male (n=11 subjects, 695 points) and female (n=24 subjects, 1805 points) humans do exist (p=0.001, chi-squared). In **Fig. 5d,e** we show this same comparison using a data set trimmed to have equal male and female sessions **(Methods: Statistical methods of data trimming).** In this figure, we use the full dataset without trimming to match. **e,f.** We compare the usage of primitives in participants with self-reported hunger **(e)** and tiredness **(f)** above and below various thresholds for male le and female participants. We find that across several thresholds, male primitives change across the threshold while female primitives. A blue tile means that the difference between groups was significant using chi-squared, while white means the difference was not significant (**Methods: Metadata collection). g,h.** We find that hunger and tiredness coincide more in male than in female humans. **i,j**. Food deprivation in rats causes a significant change in primitives (p=0.0069, chi-squared, food deprived rats = 22, 297 sessions). **k,l,m**. Differences in primitives between male (n=10, 137 points) and female (n=12, 137 points) food-deprived rats do exist indicated by a significant difference in primitive proportions (p=0.0256, chi-squared). **(k)** shows male rats, **(l)** shows female rat primitives, and **(m)** compares the primitive use of each. We use the following abbreviations: M=male and F=female. **n,o,p.** Female rat primitives are not significantly affected by food deprivation (n=14 subjects, 160 points). (**n)** shows ad libitum female primitives and **(o)** shows food-deprived female primitives, while (**p)** compares the primitive use of each. We use the following abbreviations: BL = baseline and FD = food-deprived.

#### Extended Fig. 7

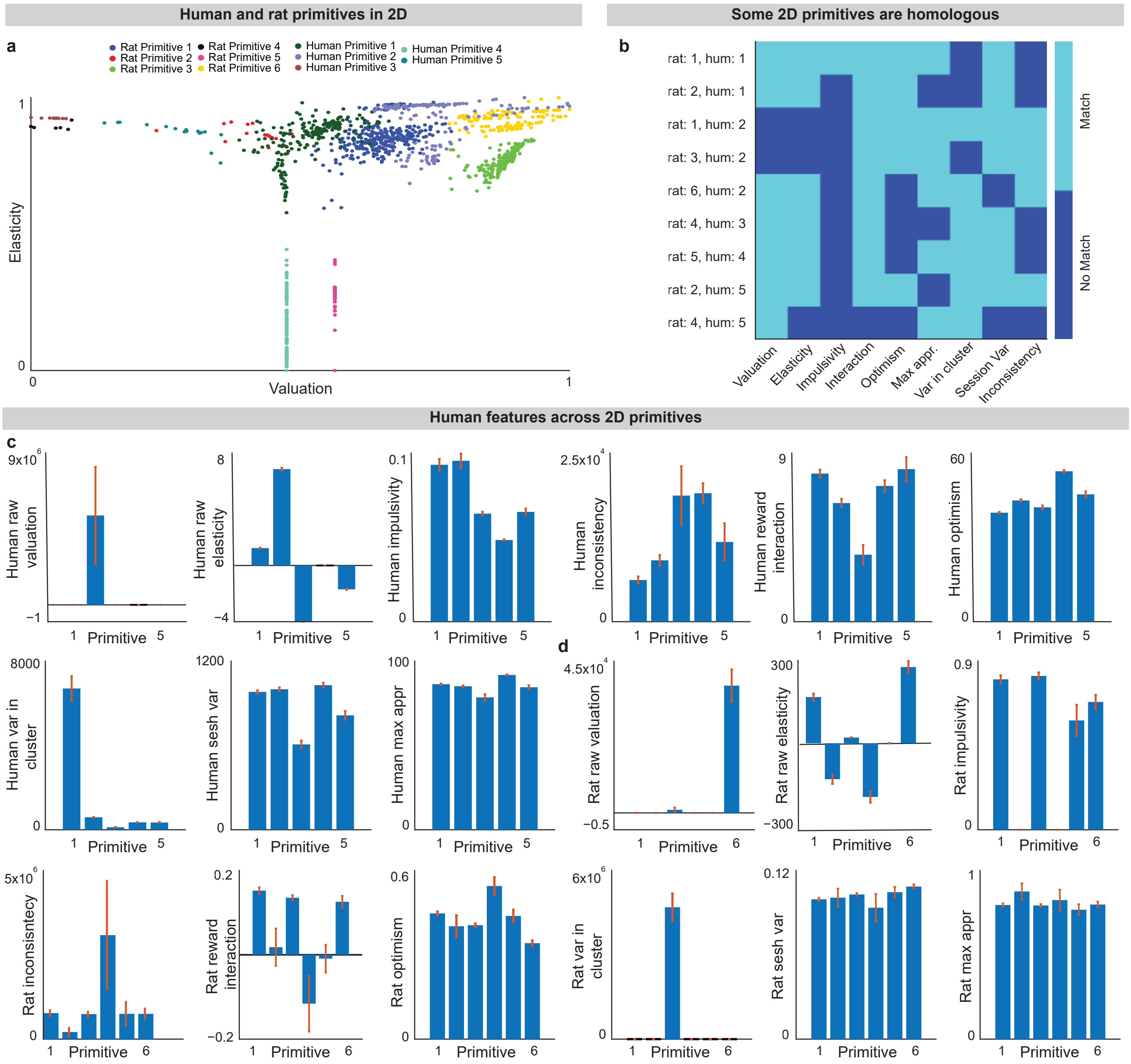

**Extended Data Fig. 7 | Similar primitives between human and rats exist**

**a.** Because rat performance was not calibrated to have a particular maximum approach rate (while humans were calibrated to have a maximum approach rate of 100), the max parameter is not directly comparable across rats and humans. Valuation and elasticity, however, are internal parameters of psychometric strategy. Therefore, we focus on these parameters in our comparison Clustering in those two dimensions, we find that human and rat primitives overlap in location. **b.** Of rat and human primitives with corresponding location, several pairs share other decision-making features as well (**Methods: Similar Primitives between Rats and Humans). c,d.** For each of the rat and human primitives, we calculate summary statistics for each feature including: “impulsivity”, "inconsistency”, “interaction”, “optimism”, “valuation”, “elasticity”, “variance of sessions within a cluster”, “variance of choices within a session”, and max approach rate **(Methods: Features using Psychometric Functions).** We use the 2-D clusters shown in (**a**) for these calculations, so that we can compare how primitives that overlap in parameter space may share other features as well.

### Extended Fig. 8

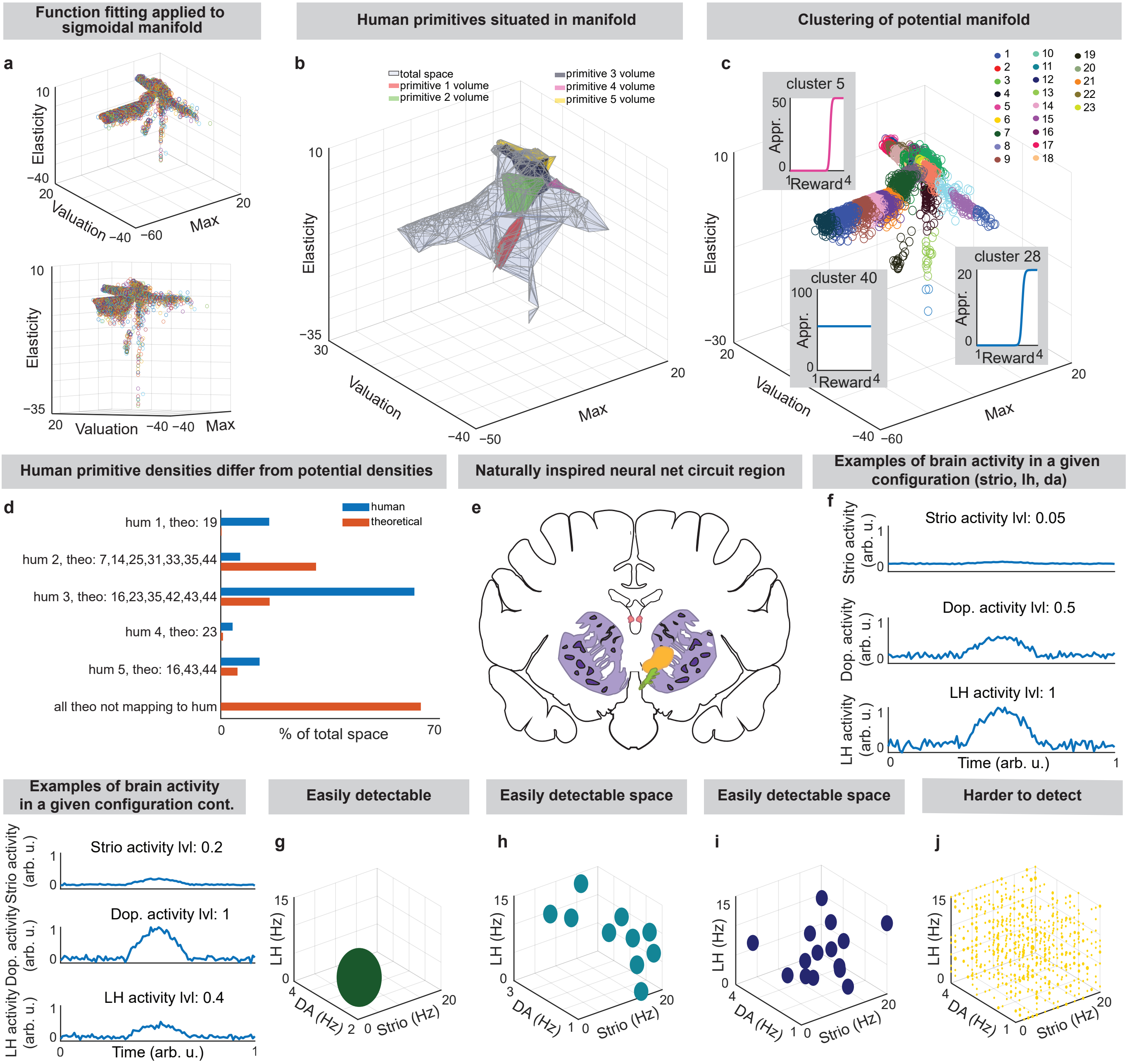

*Figure legend start on next page*

**Extended Data Fig. 8 | Sigmoidal space and naturally-inspired neural net predicts neuronal activity**

**a.** Manifold describing all sigmoidal decision-making functions, created by using the built-in Matlab fitting function to fit a sigmoid to each possible combination of approach rates across 4 levels of rewards **(Methods: Identifying Sigmoidal Space)**. **b.** Experimental clusters make up 9.7% of the total theoretical space. To find the total volume, we found the outer boundary of the manifold then calculated the volume using that boundary. We did the same for the clusters, finding the volume of each cluster individually. We then took the proportion of these volumes **(Methods: Identifying Sigmoidal Space). c.** The manifold of sigmoidal decision-making is not uniformly dense, there are certain coefficient combinations that are more probable than others. We use the built-in Matlab fitting function to fit sigmoidal curves to data **(Methods: Comparison of Primitives to Sigmoidal Space)**. **d.** The density of manifold clusters and experimental clusters located in the same place (via Bhattacharya distance) differs significantly, indicating that the mathematical processes incident to clustering does not account for the variations found in experimental data. **e.** Depiction of the brain regions  and connections included in the naturally-inspired neural network architecture, modeling the cortical-striosomal-GPi-lateral habenula-RMTg-SNc-matrix circuit (**Methods: Naturally-inspired neural network). f.** Examples of brain activity for configurations in **h,i. g-j.** Possible brain configurations for different clusters, same as in main Fig. 6e. **(h,i)** would be easily detectable by FMRI because there are many similar configurations with equal likelihood clustered in one area. **(g)** would be easily detectable because there is only one plausible configuration, with high confidence. **(j)** would be hard or impossible to detect because the signal to noise ratio is low.

### Supplemental Materials and Methods

### Statement of Open Science

Out of our commitment to fully transparent and open science, we provide all materials and data that were used in this project including the user guide to running this experiment, run-me scripts and instructions for reproducing each panel and analysis, database links to all of our data, and Github links to all of our code. Data collection was done automatically, as will be described in the Decision-Making App User Guide, and all analyses, statistics, and figures are reproducible using the steps, code, and data provided in this document.

### Research Ethics Statement

Human data collection for this project received approval for all protocols from the University of Texas at El Paso IRB on June 19, 2024 (FWA No. 00001224).

Animal data collection for this project received approval for all protocols from the University of Texas at El Paso Institutional Animal Care and Use Committee and followed the Guide for Care and Use of Laboratory Animals (IACUC reference number: A-202009-1).

### Decision-Making App User Guide

#### Purpose

We have developed the Human Model for Analysing Neuroeconomic Situations (HUMANS) app. The purpose of this app is to measure decision-making (DM) in human subjects by presenting them with a questionnaire specifically designed to elicit DM behaviours in the subject while also collecting biometric data as the subject goes through the different trials.

#### Versions of Application

There are two versions of this application. One was created for use at the University of Texas at El Paso, and another was created for use at Mount Sinai. There are three differences in these versions: where the data was stored, the collection of physiological data, and the stories used. In the version made for UTEP use, data was uploaded to the university server while in the Mount Sinai version, data was saved locally. This was done because we did not have access to the Mount Sinai server to set up a database to write to. In addition, we did not set up eye tracking and heart rate collection for the Mount Sinai version since we expected this version to be employed with fMRI and these additional setups would not be feasible within the machine. In the University of Texas set-up, we collected both eye-tracking and heart rate data.

In the Mount Sinai version, we selected stories that only had XXX qualities (decision-making boundaries) . Why? To make sure that the stories used were maximally effective and similarly used across the population, to make best use of the time under the scanner.

In this user guide, we cover the details of the University of Texas version since the data in this paper was collected using that version, and because that version was more technologically involved.

#### Data collected

Subjects are not directly identified by name in the data produced by this application. Rather, a randomised ID number between 10000 and 99999 is generated and assigned to the subject. We then collect demographic information including but not limited to sex, gender identity, age range, race and ethnicity, and education. We also collect interests such as hobbies and what kind of media they consume. We use this data to classify subjects into clusters to attempt to identify any meaningful trends.

During trials, we collect biometric data, including gaze (where the subject is looking at on the screen, pupil dilation (estimated, in millimetres), and heart rate (measured in beats per minute), to potentially measure attentiveness and stress response. All trials are time stamped from when the question is presented (trial start), to when the subject submits their answer (trial_end).

All data is packed into a PostgreSQL table and uploaded as soon as the subject submits their answer. Additionally, demographic information (identified by ID number) is stored in a local text file.

#### How it works

Each subject is presented with a selected number of scenarios, in which a scenario (story) context is given. The subject must then put themself in that situation and think about how they would react.

Next, a list of potentially rewarding options to solve the problem at hand is presented to the subject. The subject then chooses which options would be most preferable by ranking them on a continuous 0 - 100 scale. This is then repeated for a list of potentially costly or unpleasant options to solve the problem and the subject must then rank them from least unpleasant to most unpleasant on the same continuous 0 - 100 scale. We call these rewarding and costly options "preferences".

Finally, the subject is presented with a predetermined number of questions that pertain to the preferences that were previously chosen in the actual trials. The subject must then answer the question with yes, no, or maybe on a scale. In the case where two different options are presented in the trial, the subject must indicate which option they lean toward the most (this is not the same as ranking preferences). As subjects answer these questions, we collect biometric data (see "Data collected" section).

#### App operation

The app is written using mostly Python, and runs within a Flask webserver. All webpage templates are written in HTTP and styled with CSS. All data is stored in a local or remote PostgreSQL database (depending on how the app is set up), as well as locally for redundancy in case of database failures.

#### Visual of the App

Below are screenshots from one example session of the task, abbreviated to show only two trials instead of all 16. In this example, we do not show the initial calibration steps of the eye-tracker or heart-rate monitor. This is shown in a separate section below for installing and setting up the trackers. The first panel asks how many stories to view in total and some self-reported state information by the participant.

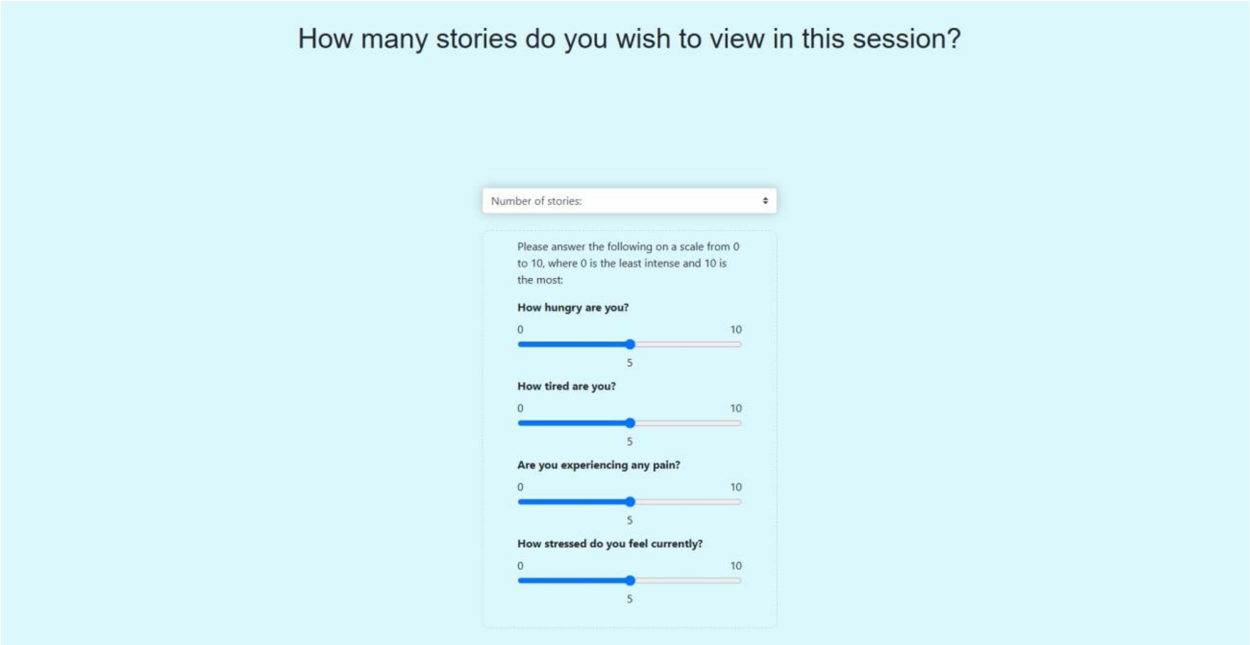

Once submitted, the app asks if the participant is new or returning. If returning, the participant goes straight through to the story panels. Otherwise, the participant is given an ID and asked to fill out demographic information.

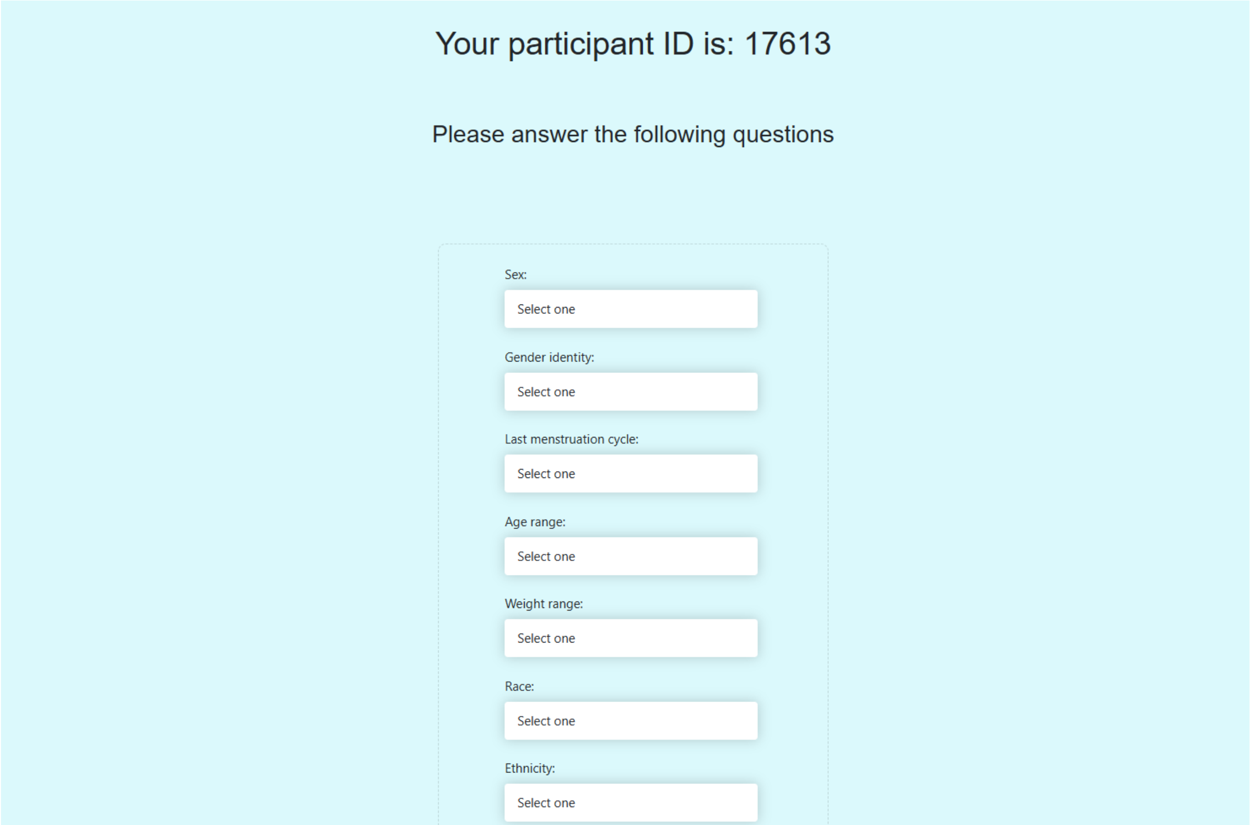

Subjects are asked about relevant story topics so that only stories of interest are presented to the subject. The story context is then introduced.

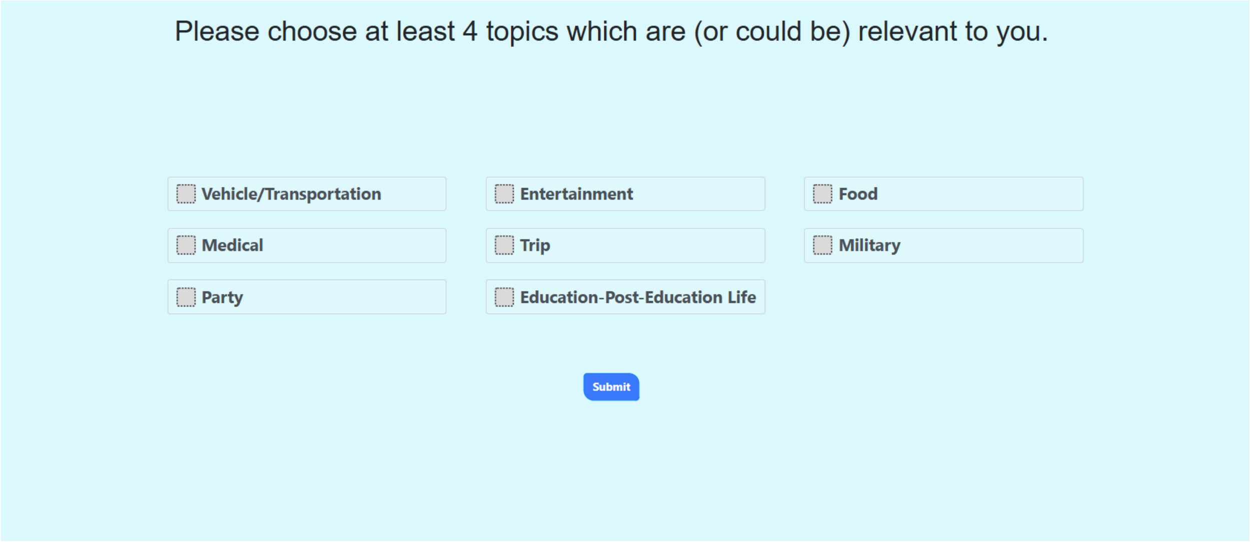

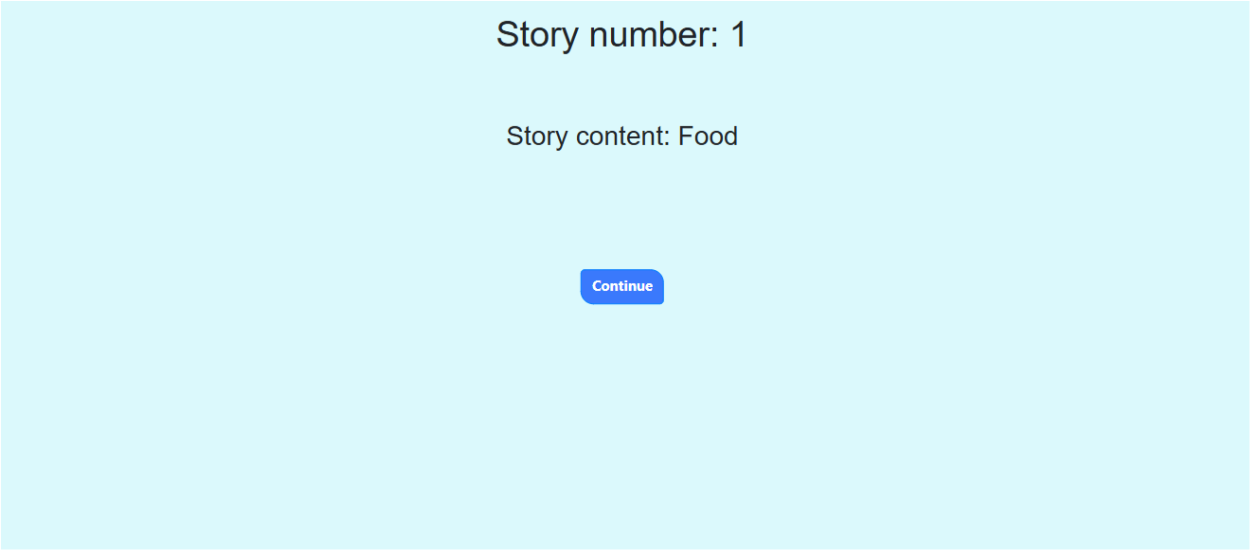

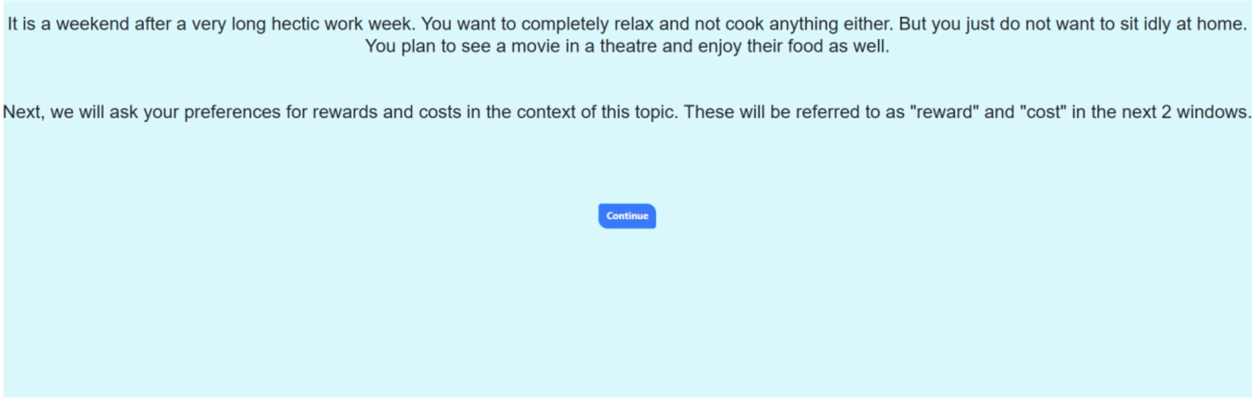

Once the story context has been presented, participants are asked about their reward and cost preferences within the story context.

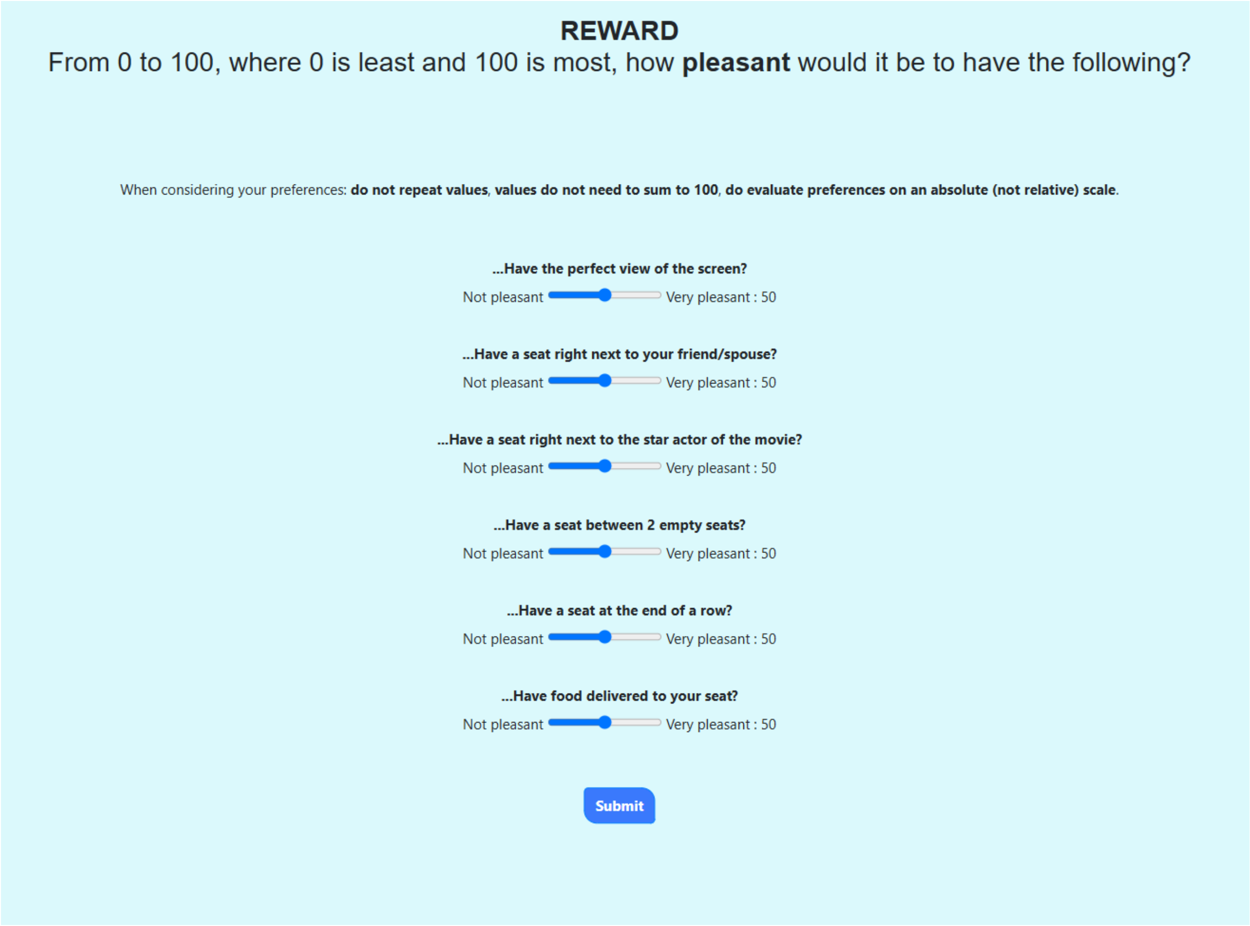
REWARD
From 0 to 100, where 0 is least and 100 is most, how pleasant would it be to have the following?
When considering your preferences: do not repeat values, values do not need to sum to 100, do evaluate preferences on an absolute (not relative) scale.
...Have the perfect view of the screen?
Not pleasant
Very pleasant : 50
...Have a seat right next to your friend/spouse?
Not pleasant
Very pleasant : 50
...Have a seat right next to the star actor of the movie?
Not pleasant
Very pleasant : 50
...Have a seat between 2 empty seats?
Not pleasant
Very pleasant : 50
...Have a seat at the end of a row?
Not pleasant
Very pleasant : 50
...Have food delivered to your seat?
Not pleasant
Very pleasant : 50
Submit

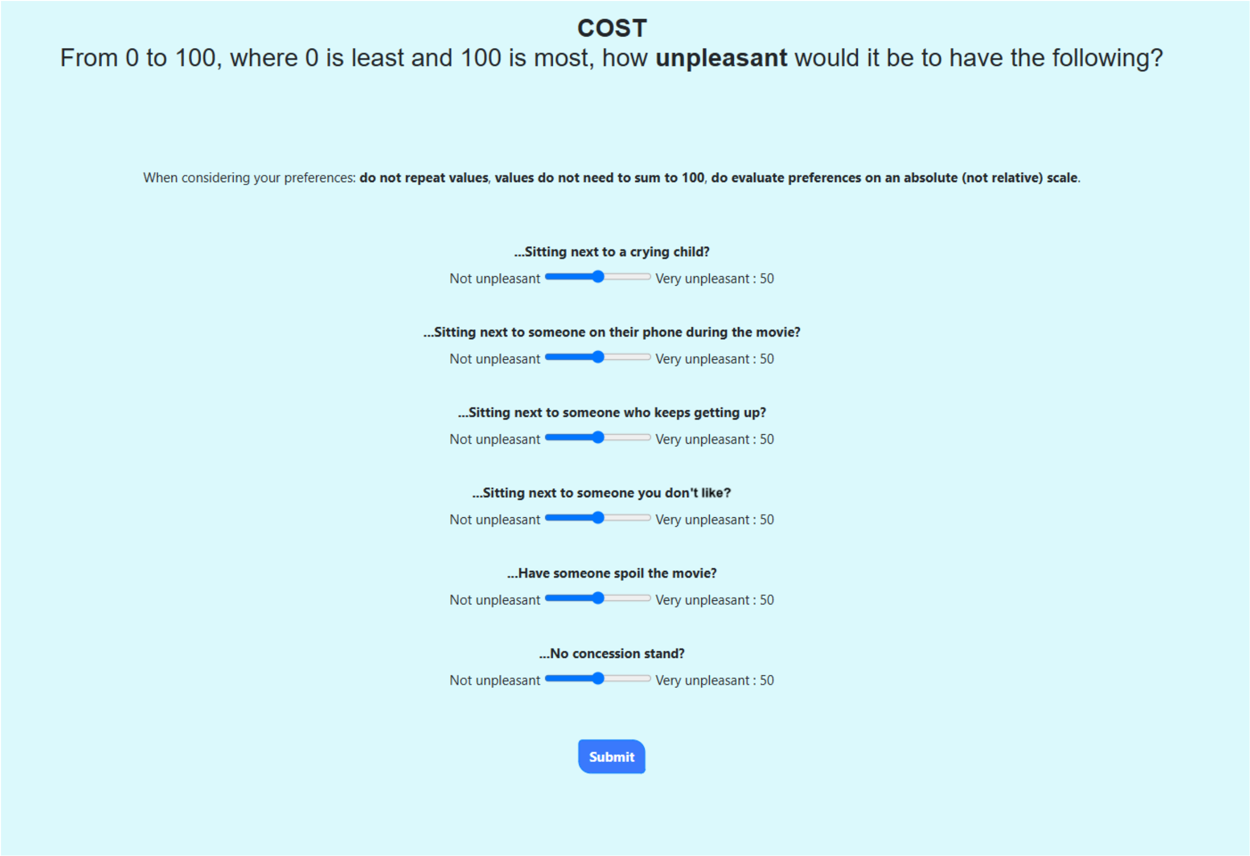

Once preferences have been submitted, the story context is again presented ahead of the trials.

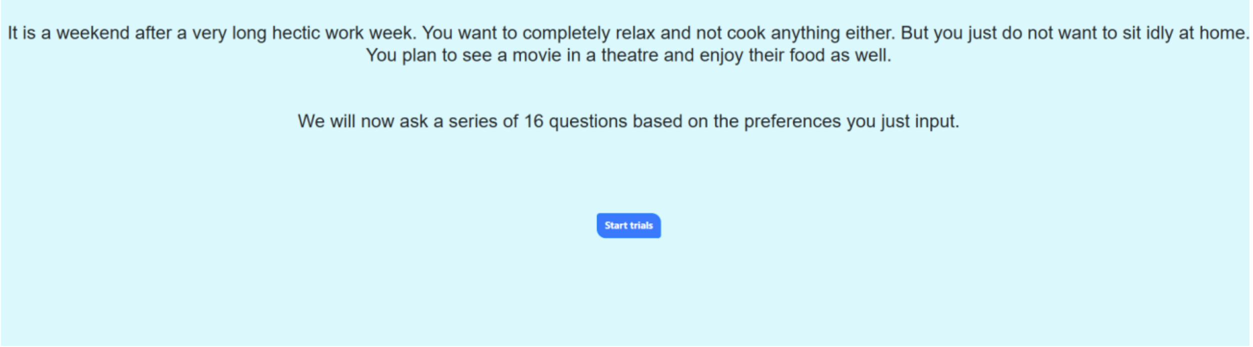

Of the six reward and six cost preferences presented, only four of each are chosen. The selection process is described in detail and the code is provided in the main methods text. For each combination of reward and cost, a trial is presented. We show two examples below.

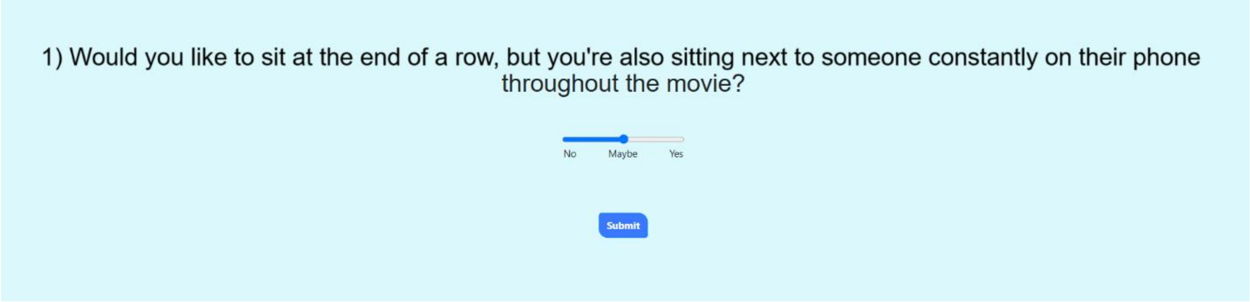

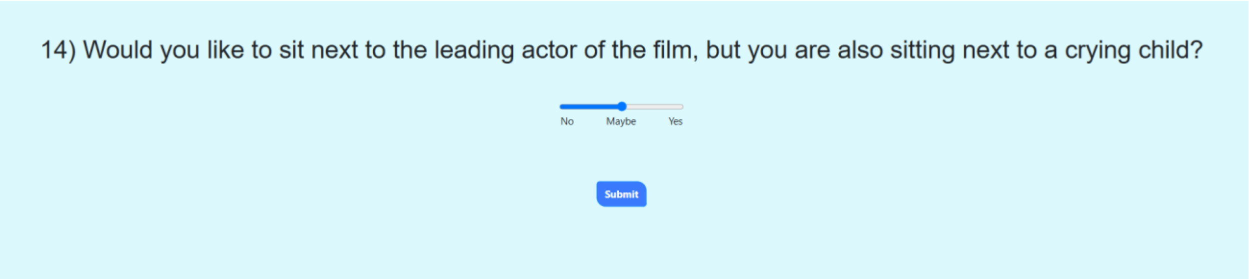

Once all 16 have been answered, subjects can decide if they'd like to reassess their preferences. If not, they can continue to finish their story session.

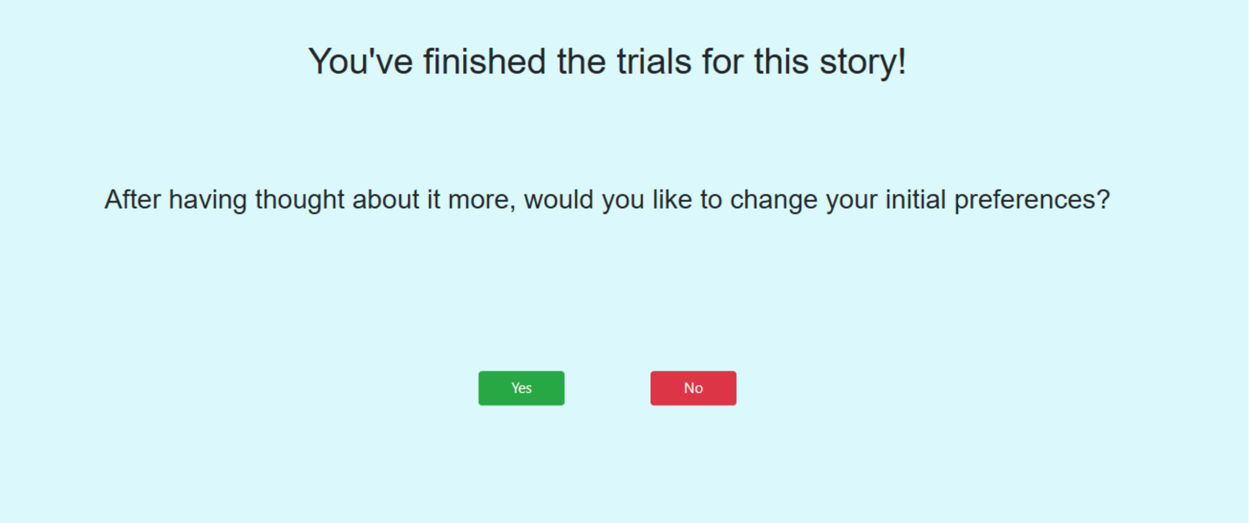

Subjects can mark how relevant the story was to them at the end of the story.

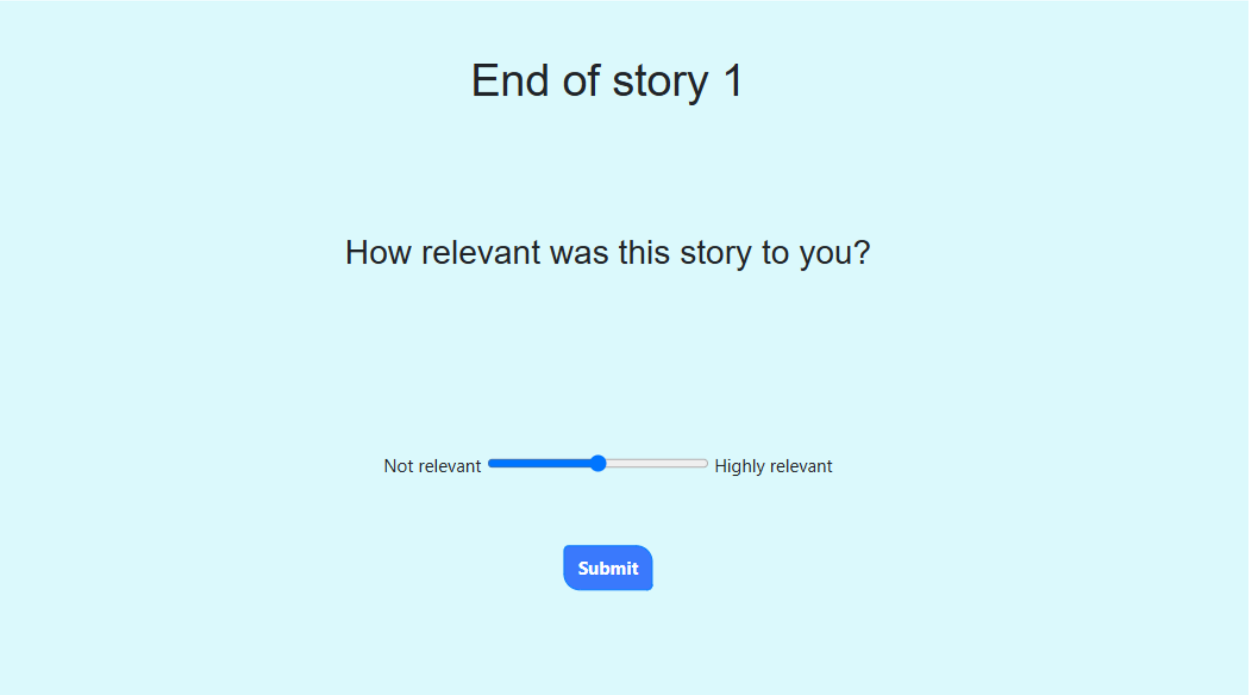

Once they have completed their sessions, any notes can be taken by the lab technician and will be uploaded to the database with the session.

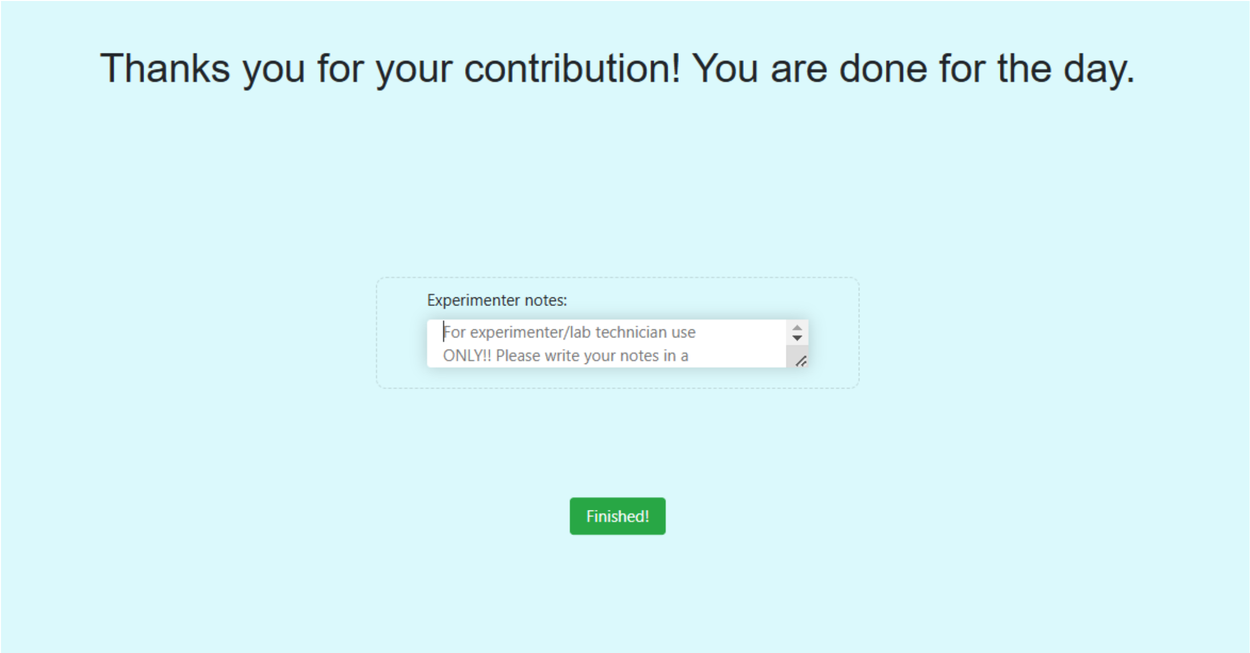

While the subject is completing their session and answering questions, their responses are automatically being captured and uploaded to our server database. At the same time, if the eye-tracker and heart-rate monitor have been set up, their physiological responses for each trial are being uploaded as well.

### What is included our repository

#### App elements

This includes files and scripts that critically pertain to the operation of the application. This includes Python scripts, HTML sheets, and CSS styling. This also critically includes the settings INI file.

#### Helper scripts

These are scripts that isolate certain functionalities that are built into the app but are not referenced from the app's code. Such scripts can be changed and run independently from the app without any impact. The following is a full list of these scripts:

- create_map.py
- distribute_stories.py
- grab_ids.py
- import_demodata.py
- randomise_relationship_levels.py
- write_to.py
- breakdown_stories.py

#### Resource links

1. HUMANS App Github Repository*
2. ANT+ Heart Rate (HR) monitor (example)
3. ANT+ USB Antenna (example)
4. Eye tracker (Tobii Pro Spark)
5. Tobii Eye Tracker Manager (software)
6. Tobii Pro SDK (software)
7. Tobii Pro Spark Runtime Driver (software)**
8. Tobii Pro Spark User Guide (documentation)**
9. Pulse Monitor (software)

* This guide pertains to only the code contained in the main branch of our Github repository. The code contained in any other branch is not documented in this guide. Please ensure you are downloading code from the main branch.

** Which runtime driver and documentation will depend on what eye tracking system you are using. In this guide, we assume the Tobii Pro Spark eye tracker, and all links in this guide were made with that in mind. If your eye tracking system differs from ours, consult the manufacturer's instructions for setup and installation.

### Setup

#### Installing the app

Our Human Decision-Making App (HDMA) is contained in our github repository. After zipping all files and downloading, extract the files onto any location on the hard drive.

#### Tweaking app settings

Navigate to the directory where you extracted the decision-making app files to. Navigate to the 'bin' folder and open 'settings.ini' with your text editor of choice (Notepad, Notepad++, Atom, etc...).

We first enter the PostgreSQL database credentials. This will look different depending on how the database was set up. However, refer to the following guidelines (or to the "App Settings" section) to help you know what each parameter is under the 'postgresql' section, or seek help from your database administrator:

- `host` : The host computer's IPv4 address or domain name. If the HMDA is running on the host computer, enter the word localhost.
- `database` : The name of your PostgreSQL database.
- `port` : The host computer's port-forwarded port for servicing database queries (try 5432 if you don't know)
- `user` : The PostgreSQL username. Keep in mind that the account that the HMDA uses must have database read-write privileges.
- `password` : The password tied to the account identified by the

‘user’ parameter.

Next, we need to change the settings that will affect how the HMDA behaves. This will look different depending on your own study’s needs. However, refer to the following guidelines to help you know what each parameter is under the ‘app_settings’ section:

- data_table : The database table to insert data into
- auto_create_table : If the table specified above does not exist in the database, the app may be able to create it using the participant's data. If auto_create_table is set to 1, the app will attempt to create the target data table once it has all the required data from the user. If set to 0, the app will simply crash when attempting to upload data if the table does not exist in the database.
- enable_consecutive_users : Enables back-to-back use of the app without the need to close and re-run startup.bat (which restarts the flask server. If set to 1, the app will reset all global app and user parameters to default when loading the landing portal (setup_session.html, routed as '/'). If this option is not enabled (set to 0), the flask server must be closed and re-run. Any changes to the source code or templates are not affected by this option.
- data_upload : Ensures that data is automatically uploaded to the database after each session.
- unique_ids_from : Controls where IDs are read from when trying to generate a unique ID for each user. Possible values are "database" and "local". "database" will look into the database and find all the unique IDs stored there. "local" will only look to the "/data/" directory to find all the taken IDs. The app will then generate a new random ID that does not exist in the retrieved list of existing IDs.
- next_story_from : Controls where the next story index is referenced from. Possible values are "database" and "local". "database" will look into the database to find the next story the user will see by counting all the unique entries of 'tasktypedone'. "local" will look into the user's local demographic info record in '/data/' and read the

'next_story_index' entry.

- timestamp_timezone : The time zone that timestamps are collected in. Recommend 'UTC', if this parameter is not set, the app will default to 'UTC'.
- minimum_topics : The minimum amount of topics that the subject must select. Replaces 'min_stories_to_choose' from version 30.
- questions_per_story : The number of questions selected to show the user PER STORY.
- ignore_legacy_story_data : How the app handles story data ('pref_stories' and 'story_order') from legacy app versions. If this is set to 0, the app will NOT ignore legacy story data from returning users and continue the remainder of the users' sessions with only 'approach_avoid' task types, keeping their preferred stories and previously calculated story order. If set to 1, the app will discard this information and ask the user to restart their sessions to add the new task types to their story data.
- randomise_relation_levels : FOR SOCIAL TASK ONLY: Whether to randomly choose a random relationship keyword and replace it into the text of the social task stories. If set to 1, a random relationship keyword will be picked from the list under the 'relation_levels' list parameter and replaced into every snippet of text throughout the entire story.
- relation_levels : FOR SOCIAL TASK ONLY: The words to look for and replace in story text. The app will randomly sample a word and replace it throughout the entire story text.
- relation_level_stories : FOR SOCIAL TASK ONLY: Stories where replacing the relationship level can be done
- validate_stories : Validates that the stories described in the 'Human DM Topics' relationship table are actually contained in '../stories/task_types/'. Stories that don't exist in that directory will be deleted from the pool of stories that can be selected for each subject. Setting this parameter to 1 can cause errors if the validated story pool ends up being smaller than the sample size for each task type (see below). If this is the case, check that there are enough stories for each task type in '../stories/task_types/'. Conversely, not validating the stories increases the odds of trying to access a story that does not exist, causing a different error.

- The next 7 parameters describe how many stories are to be selected (sample size) per task type at each user's first session:
    - approach_avoid
    - benefit_benefit
    - cost_cost
    - moral
    - multi_choice
    - probability
    - social

These settings may be tweaked further later, but keep in mind that they must be tweaked **before** running the HMDA.

Once you have made the changes needed, save the file and exit the text editor.

### Installing the eye tracker

#### Hardware and software setup

The eye tracker setup requires one hardware device, and three different software. Refer to the "Resource links" section of this guide. Download and install items 5, 6, and 7 first, then mount your tracking device following the manufacturer's guidelines (item 8). Connect the device to any available USB port, preferably one directly on the computer tower and not through a USB hub, as this could cause issues with data throughput.

Test the eye tracker by opening the eye tracker manager (item 5). Your eye tracking device should show up in the list of available devices at the top.

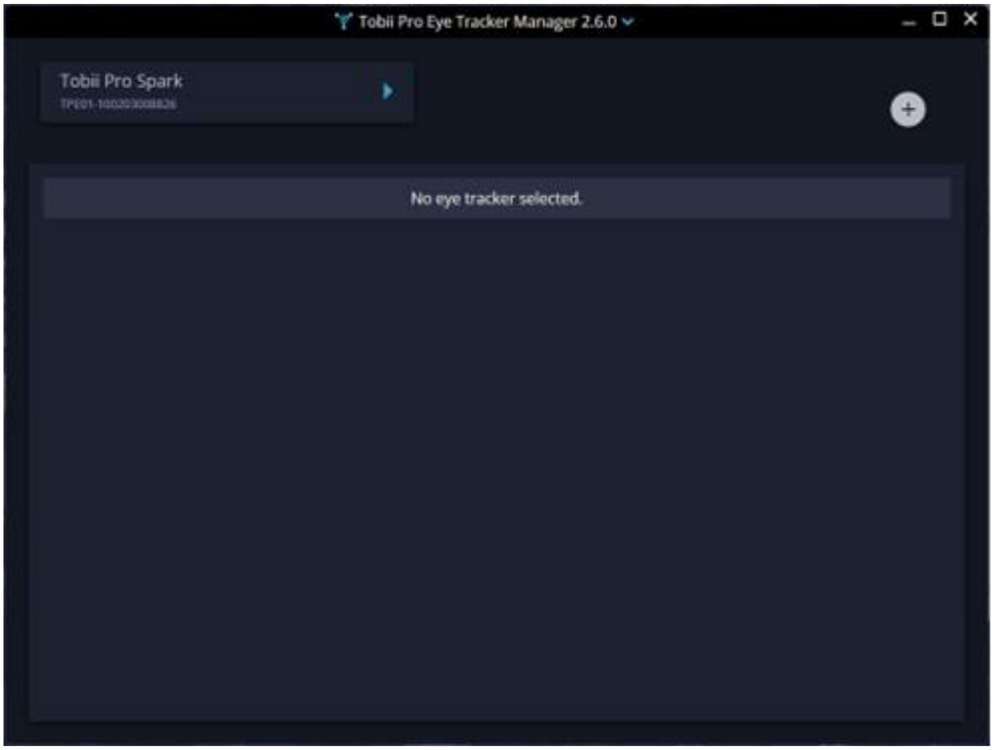

Click your device and the user interface (UI) will expand. Toggle both ‘gaze visualisation’ and ‘position guide’, then look around and see if the gaze visualiser (in pink in the example below) moves to where you are looking. If it doesn't, click the ‘Calibrate’ button and follow the calibration steps that show on screen.

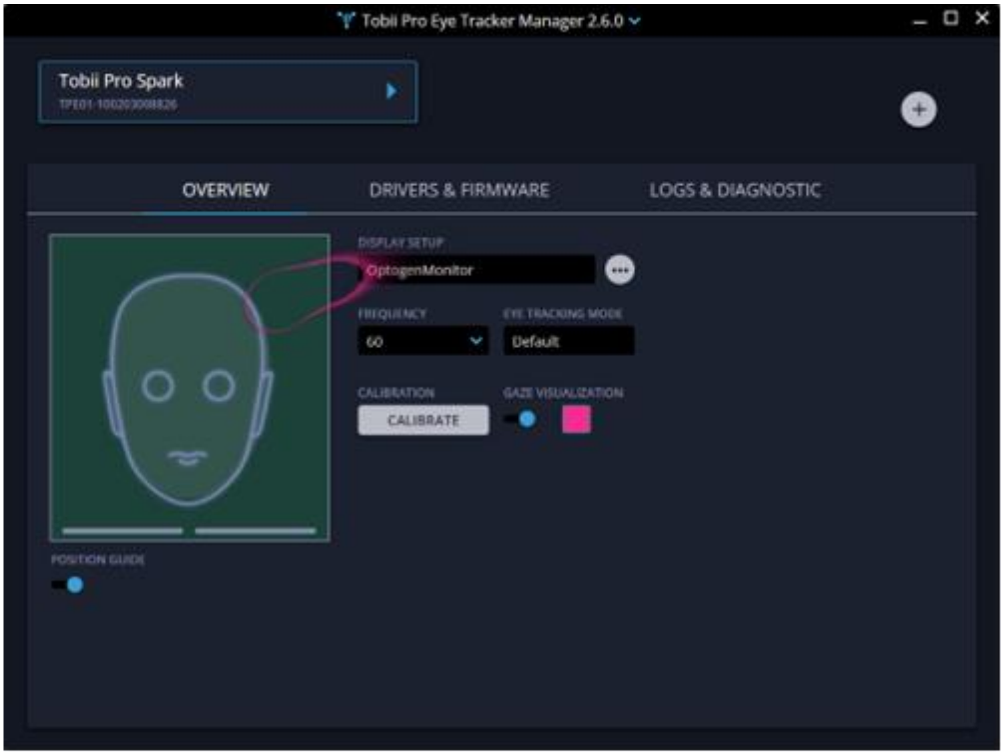

**Tweaking app settings**

Navigate to the directory where you extracted the decision-making app files to. Navigate to the 'bin' folder and open 'settings.ini' with your text editor of choice (Notepad, Notepad++, Atom, etc...). Make the following changes under the 'eye_tracker' section:

- `manager_install_path=C:\Users\{YOUR_WINDOWS_USERNAME}\AppData\Local\Programs\TobiiProEyeTrackerManager\TobiiProEyeTrackerManager.exe (this may differ depending on where you installed the Eye Tracker Manager program)`
- `subscriptions=['gaze','position'] ('openness' is also valid, however it may not be supported for your device)`
- `eyetracker_index=0 (if you only have one eye tracker connected, otherwise specify your device's index)`
- `use_eyetracker=1`

Save the file and exit the text editor.

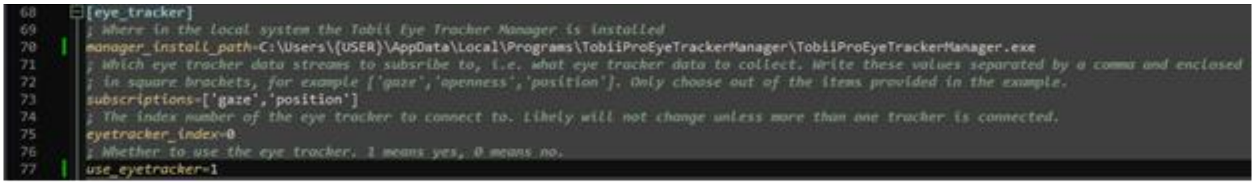
```
[eye_tracker]
; Where in the local system the Tobii Eye Tracker Manager is installed
manager_install_path=C:\Users\{USER}\AppData\Local\Programs\TobiiProEyeTrackerManager\TobiiProEyeTrackerManager.exe
; Which eye tracker data streams to subsribe to, i.e. what eye tracker data to collect. Write these values separated by a comma and enclosed
; in square brackets, for example ['gaze','openness','position']. Only choose out of the items provided in the example.
subscriptions=['gaze','position']
; The index number of the eye tracker to connect to. Likely will not change unless more than one tracker is connected.
eyetracker_index=0
; Whether to use the eye tracker. 1 means yes, 0 means no.
use_eyetracker=1
```

#### Installing a heart rate monitor

Throughout this section, refer to the "Resource links" section of this guide. If you are following our methods to collect heart rate data, you'll need to install the ANTUSB driver for Windows. You may follow this guide to do so.

##### If using an external program like Pulse Monitor

Make sure your program of choice is compatible with ANT+ devices. After installing your program, be sure to set up your device in whichever way the program needs. Here, we provide an example using Pulse Monitor (item 9).

Open Pulse Monitor and click the 'Devices' button.

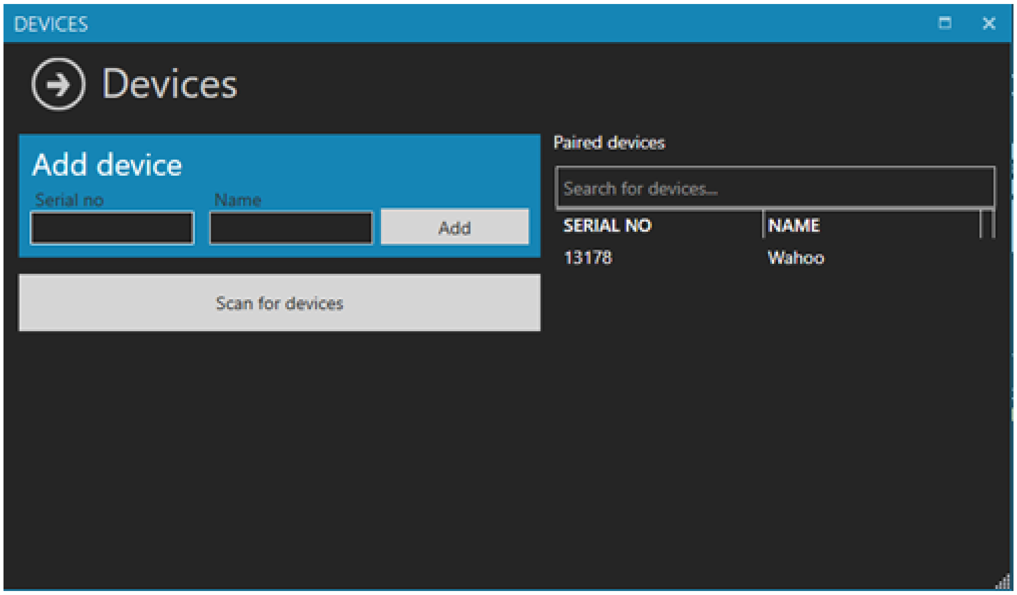

Connect an ANT+ USB antenna (item 3) and power up your heart rate monitor (item 2). Enter the heart rate monitor's serial number and name in the corresponding dialogue boxes, then click the 'Scan for devices' button and wait for your device to appear. Once your device appears, click the 'Add' button. Close this dialogue box when you're done.

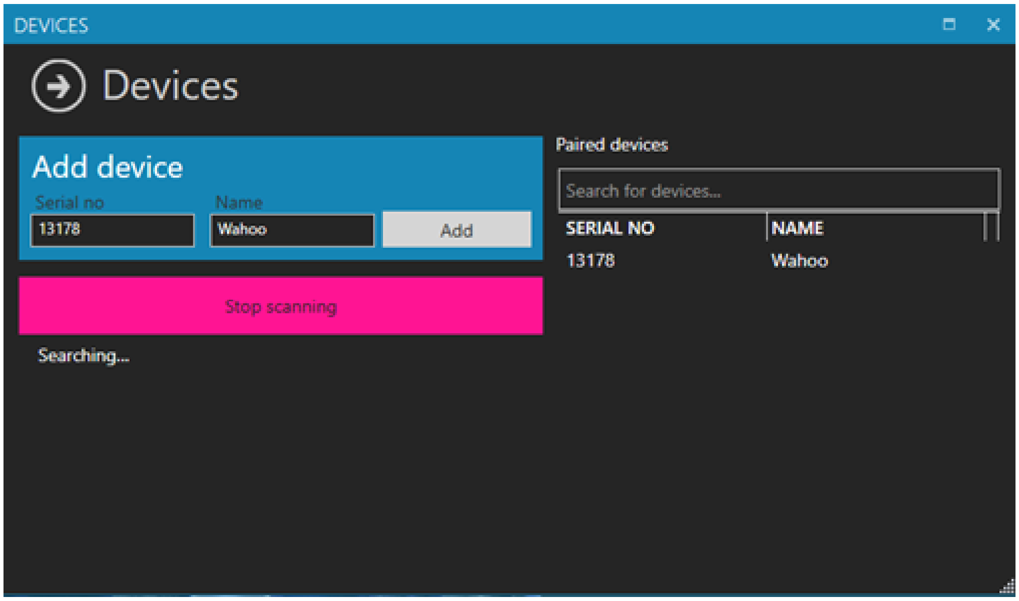

Next navigate to the directory where you extracted the decision-making app files to. Navigate to the 'bin' folder and open 'settings.ini' with your text editor of choice (Notepad, Notepad++, Atom, etc...). Change the 'use_external_app' parameter under the 'hr_tracker' section to 1 and copy your heart rate monitoring app's install directory path to the 'external_app_install_path' parameter (see example below for suggested format). Finally, set 'use_hrtracker' to 1.

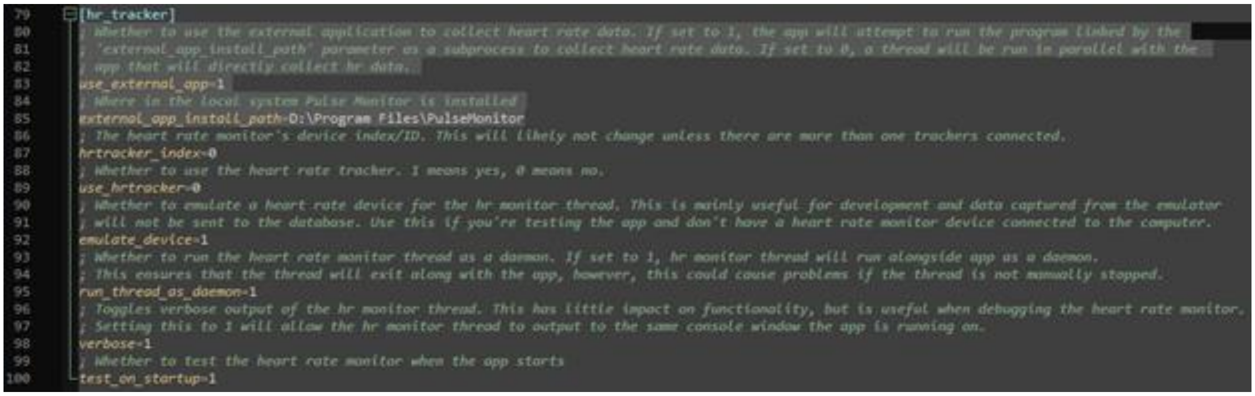

*Example is 'settings.ini' open in Notepad++ with Dark Mode enabled.*

This will signal the HDMA to open Pulse Monitor as a subprocess to collect heart rate data. Keep in mind that since this program runs asynchronously from the HDMA, the data produced by this program will not be directly uploaded to your database, rather, it must be exported, saved, proprocessed, and then uploaded.

#### If using 'built-in' heart rate monitoring

*Note: This feature is currently experimental and may not function as expected.*

Navigate to the directory where you extracted the decision-making app files to. Navigate to the 'bin' folder and open 'settings.ini' with your text editor of choice (Notepad, Notepad++, Atom, etc...). Make the following changes under the 'hr_tracker' section:

- `use_external_app=0`
- `use_hrtracker=1`
- `hrtracker_index=0 (if you only have one heart rate monitor connected, otherwise specify your device's index)`
- `emulate_device=0`
- `run_thread_as_daemon=1 (optional)`
- `verbose=0 (optional, can be set to 1 if you want the heart rate monitor output displayed on the command line)`
- `test_on_startup=1 (optional)`

Save the file and exit the text editor. These settings will ensure that the heart rate monitor data is directly collected by the HDMA and stored in the database. No additional steps are required.

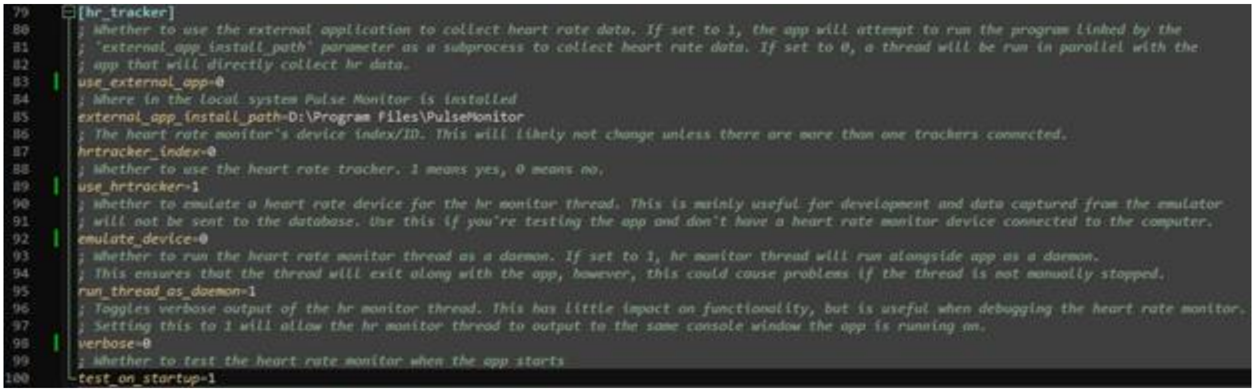

### App usage

#### Materials required

- HUMANS App (see GitHub repository under "Resource Links")
- ANT+ Heart Rate (HR) monitor (example)
- ANT+ USB Antenna (example)
- Alcohol wipes
- Keyboard and mouse
- *If subject requires:* Eye correction (for example eyeglasses, reading glasses, blue light filtering glasses, or contact lenses, if applicable)
- Eye tracker (Tobii Pro Spark)
- Tobii Eye Tracker Manager (software)
- Tobii Pro SDK (software)
- Tobii Pro Spark Runtime Driver (software)
- Pulse Monitor* (software)

* The use of Pulse Monitor is a temporary solution to collecting heart rate data. This software will be optional for later versions of the app. If you are running version 32.1 and later, no setup is needed for the heart rate monitor if the 'use_external_app' option is set to 0.

#### Running HUMANS

Double click on the batch script named **'startup.bat'** in the DM app folder.

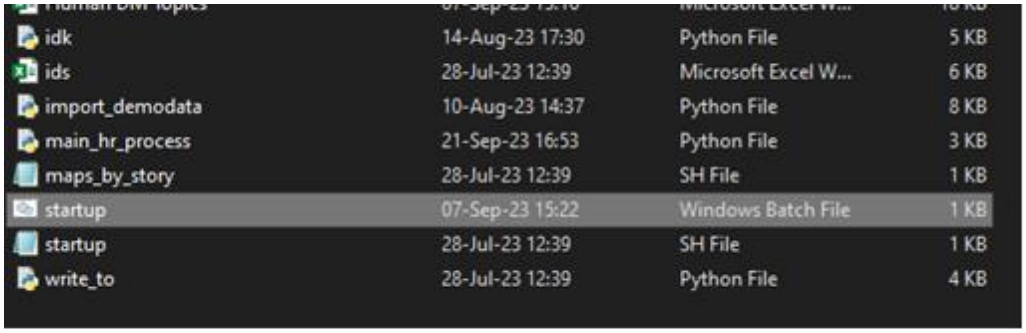

This will open a command line window, which will remain open as long as the app is running. **Do not close this window!!** This window is the program that powers the web app and will display information essential for troubleshooting.

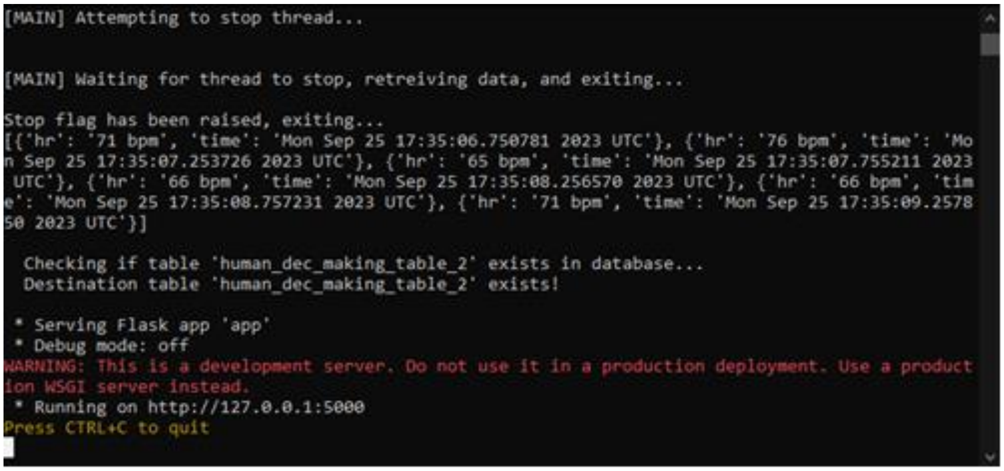

#### Subject Preparation

Before beginning a session, prepare the subject for the session by doing the following:

1. Allow the subject to choose whether they would like to wear the heart rate monitor. If they accept to wear the tracker, allow them to put on the tracker wherever they are comfortable doing so (for example, the restroom). If they do not accept to wear the tracker, do not force them, simply take note of this so that the absence of HR data is noted and move on.
2. Ask the participant for any relevant information you may need in private, make sure to store this information in a secure location. This may include:
    a. Consent forms
    b. Name to be associated with ID numbers for bookkeeping purposes
    c. Eyesight conditions that may require corrective lenses (see 'Eye Tracking Study Considerations')
    d. Use of pace-makers or health conditions that could complicate collection of heart rate data.
    e. Whether they choose to not wear a heart rate monitor.
3. If this is the participant's first session, give them a run-down of what they can expect from the recording session. Mention the eye tracker, what it does, and what it does not collect (the subject will notice the tracker turn on and off throughout the session, this could be intimidating to some participants!).
    a. The eye tracker is not programmed to capture any images during the trial. Only gaze (where the subject is looking on the screen), pupil dilation, and user position in front of the tracker are collected and recorded.
    b. Make sure that they know that they may choose not to disclose certain demographic information that the app asks for, and that their data will only ever be identified by a randomly generated ID number and not by their name.

4. **Ensure that the participant is wearing eye correction if they require it!** Eye correction such as contacts and eyeglasses must be worn during the eye tracker's calibration and all the way into the end of the session.
5. Allow the participant to get comfortable in their seat. Any adjustments to chair height or positioning can be done during eye tracker calibration.
6. Allow the participant to answer the questions in the first screen of the app. Ask them to let you know before continuing to the next screen.

#### Eye Tracker Calibration

Every single subject should go through the calibration process at least once before starting the trials. Remember that eye corrections such as eye glasses or contact lenses must be worn during this process for calibration to be accurate!

7. Upon continuing to the second screen, one or two* windows will be opened. Bring only the **Tobii Eye Tracker Manager** window to focus. Click on the button that says 'Pro Spark' on the top of this window.
    a. **If Tobii Eye Tracker Manager does not open, stop and check that the eye tracker is connected to the computer and that the 'use_eyetracker' option is set to 1 in '\bin\settings.ini'.**
    b. The app will ask you (through the command window) if you wish to continue without the eye tracker. Type 'Y' and press Enter on your keyboard to continue with the eye tracker disabled; or type 'N' to exit the app and troubleshoot.
    c. The use of Pulse Monitor is a temporary solution to collecting heart rate data. This software will not be needed for later versions of the app.
8. Ask the participant to get comfortable in their seat and click the 'Calibrate' button. Instructions will show up on-screen and calibration will begin.
9. After calibration is done, **close** ('X' out of) the **Tobii Eye Tracker Manager** window and the app will continue.
10. Allow the participant to click through the next screen ('Are you a new participant?').
11. If the subject is wearing the HR monitor and/or 'use_eyetracker' in '\bin\settings.ini' is set to 1, continue to the Heart Rate Monitor Setup section, otherwise skip ahead.

#### Heart Rate Monitor Setup (optional)

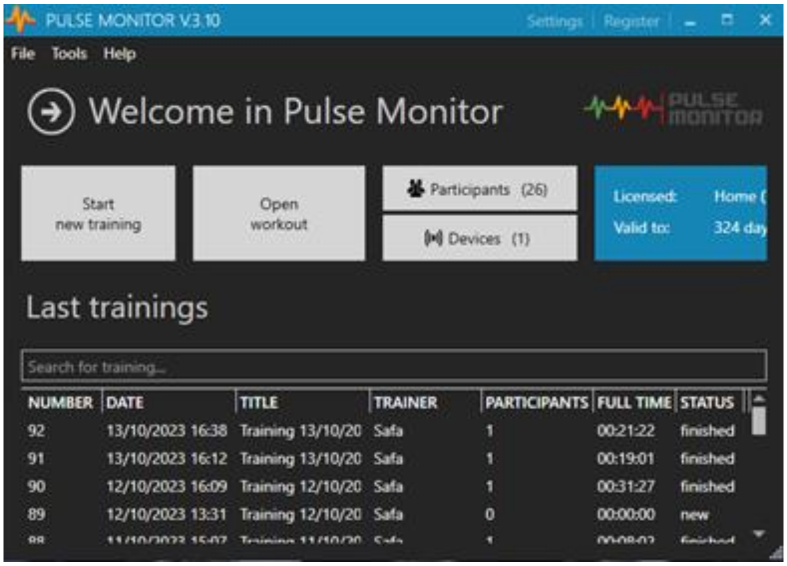

12. If the subject is a new participant, register them into the **Pulse Monitor** software by clicking the ‘Participants’ button. Click on ‘Add Participant’ and allow the subject to input their information; ask them to enter their generated ID number (from the web browser) under ‘name’ and ‘pseudo’. Click ‘Save’ when done.
    a. None of this information is used for our study, we only input this information because otherwise the Pulse Monitor software will not allow us to collect HR data.
13. Click on ‘Start New Training’. Add the participant’s ID number to the list of participants in the ‘workout’ and choose ‘Wahoo’ from the ‘Device’ dropdown menu. Finally, click ‘Add’ and ‘Go to practice’.

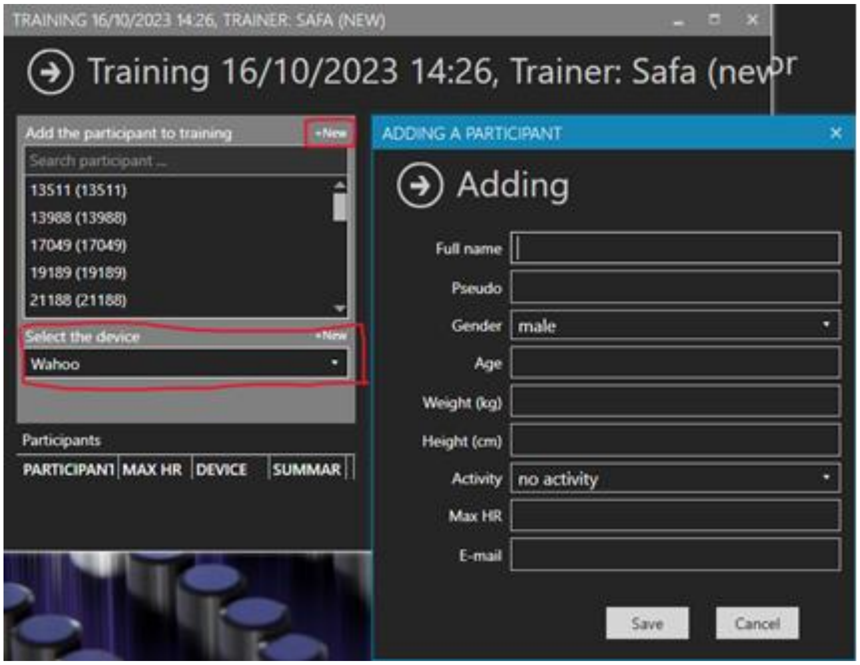

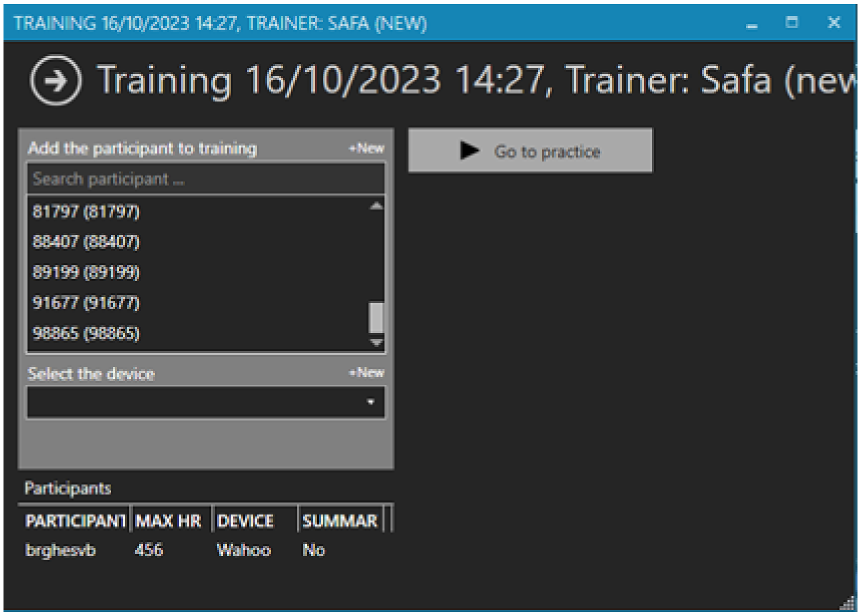

14. Start the ‘workout’, click the small grey button on the bottom right of the screen, and set this window aside.

#### Final setup

15. Take note of the participant’s ID number and ask them to do the same. Allow them to continue filling out the questionnaire.
16. The session will officially start as soon as the participant submits this questionnaire.

#### Ending a session

17. The ‘You are done for today!’ screen indicates the end of the session. Ask the subject to step away from the computer at this time.
18. Write down any notes about the session you wish to let the data analysts know. Such notes may include:
    a. Interruptions due to environmental factors
    b. Any comments that the participant may have had
    c. Catastrophic app failures
    d. Having to split the session into two
19. Click on the green button that reads ‘Finished!’.
20. Ask the subject to remove the heart rate monitor. If you are using external software (such as Pulse Monitor) to collect heart rate data, do the following:
    a. Stop the ‘workout’ on the Pulse Monitor software.

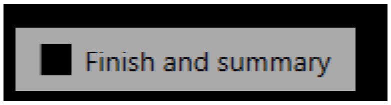

    b. Export the subject’s data by right clicking on their session summary and clicking

‘Save full HR data’.

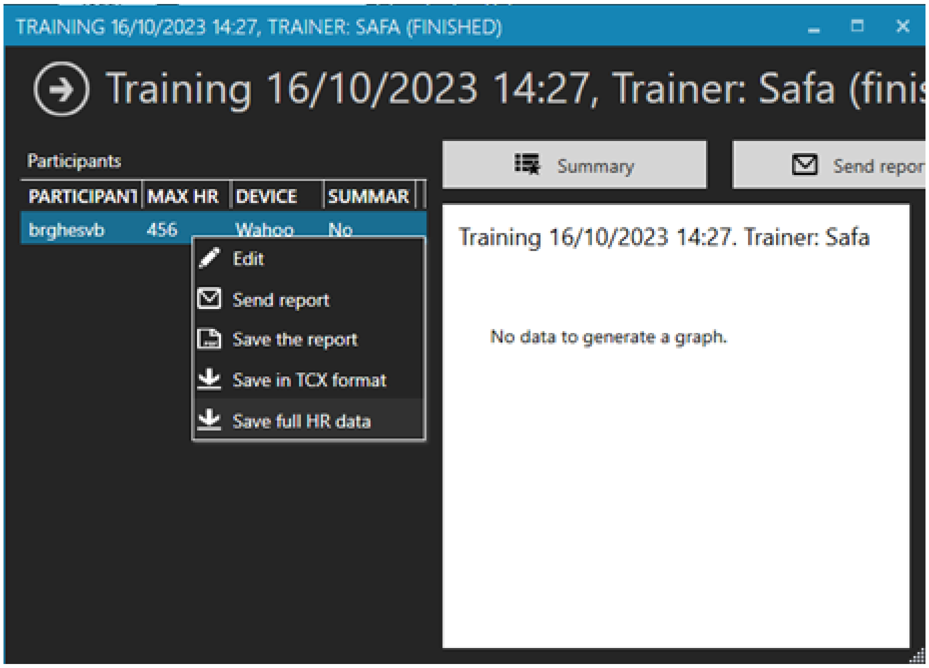

   c. Save this file under the directory named after the participant’s ID number in ‘\data\’ with the following format:

      *‘{participant ID}_{mm}-{dd}-{yyyy}_{hh}-{mm}-{ss}.csv’*

21. Wipe down the heart rate monitor with alcohol wipes for the next participant to wear.
22. If ‘enable_consecutive_users’ is set to 1 in ‘\bin\settings.ini’, the app will be ready to take the next subject. Otherwise, close the command window, run **‘startup.bat’**, and refresh the app in the browser window.

### Troubleshooting

As mentioned before, a command window will open whenever you run the app. This command window will be your best friend during troubleshooting! The program that powers the app will display information such as...

1. The user’s retrieved or generated ID number.
2. The way in which the app samples stories to add to the user’s **story order**.
3. The user’s story order.
4. The current **task type**, story number, and story index in the **story order** list.
5. The current story **topic**.

This information is displayed to help you identify where issues come from whenever they arise. The command window is also where the program will dump a description of the issue it ran into. These messages can be cryptic and might seem difficult to understand, however we'll cover some of the most common problems you may encounter, where they're likely to come from, and how to fix them.

#### Story data format

In this section, we refer to '**story data**' as any content that was written for each specific scenario that the app displays. This content is written by hand, then run through a script that breaks each scenario down into four different files; *context.txt, pref_cost.txt, pref_reward.txt, and questions.txt*.

Each of these files has to be in the correct format for the app to recognise what is in the file. Some of the most common problems are caused by the formatting of these files. The following is the expected format for each of the files listed above:

`context.txt`

Each story context should be written in one paragraph, meaning no line breaks such as new paragraphs. Punctuation is allowed, but no unicode characters are allowed.

`pref_cost.txt and pref_reward.txt`

1) The content of these files must be in list form
2) Each item must be preceded by a number, an unpaired right parenthesis, and a space
3) Do not add an indent before the number
4) No new lines are allowed within one list item
5) Punctuation is allowed but unicode characters are not
6) Do not add empty line in-between items, before, or after the list

`questions.txt`

Question items must be in list form? (R1, C1)
Each element in this list is un-numbered, un-bulleted, and starts in a new line, do not add spaces at the beginning? (R1, C2)
Questions must begin with a capitalised word and end in a question mark, period, or exclamation point. (R1, C3)
Each item must also end with a code which represents a reward and cost level before the new line, enclosed in left and right parentheses?

```
(R1, C4)
The cost and reward level codes do not need to be separated, but
should contain either a capital R or C, followed by a number! (R1, C5)
Parenthesis are allowed in the middle of a question (the app will
simply display them as they are), but be careful to also include the
code at the end? (R1, C6)
The code follows a certain pattern, but list items do not have to be
ordered by code? (R2, C1)
Breaking sentences up is allowed. Each sentence in the questions will
be stitched together and shown in a single line? (R2, C2)
```

#### Problems with story data

Problems with **story data** might be the most common problem you'll see. They will manifest themselves as one of many types of errors ranging from indexing errors to type casting errors. Here are few examples and how to fix them:

##### `ValueError: invalid literal for int() with base 10: '1.'`

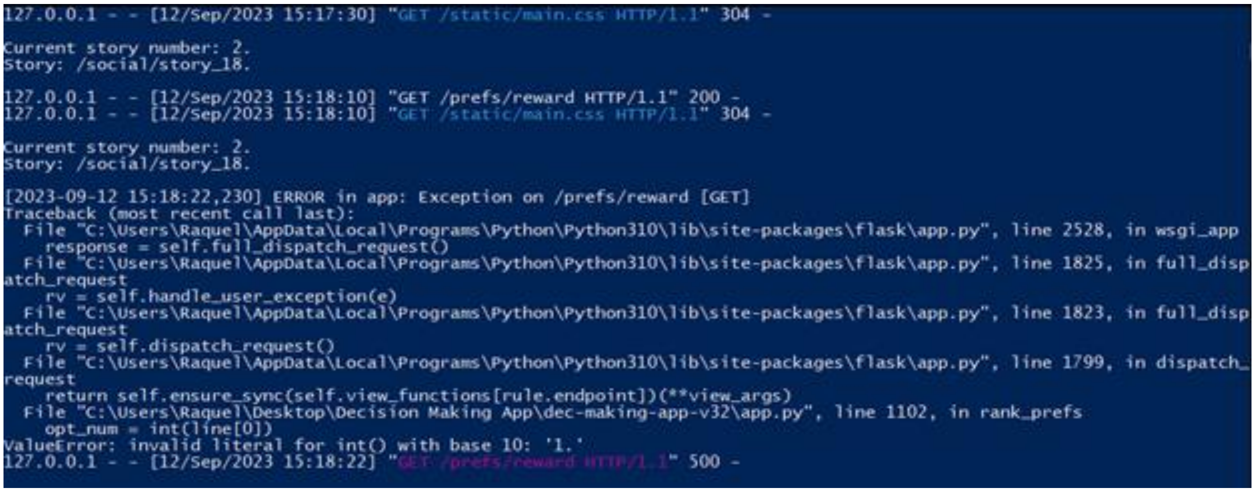

```
127.0.0.1 - - [12/Sep/2023 15:17:30] "GET /static/main.css HTTP/1.1" 304 -

Current story number: 2.
Story: /social/story_18.

127.0.0.1 - - [12/Sep/2023 15:18:10] "GET /prefs/reward HTTP/1.1" 200 -
127.0.0.1 - - [12/Sep/2023 15:18:10] "GET /static/main.css HTTP/1.1" 304 -

Current story number: 2.
Story: /social/story_18.

[2023-09-12 15:18:22,230] ERROR in app: Exception on /prefs/reward [GET]
Traceback (most recent call last):
  File "C:\Users\Raquel\AppData\Local\Programs\Python\Python310\lib\site-packages\flask\app.py", line 2528, in wsgi_app
    response = self.full_dispatch_request()
  File "C:\Users\Raquel\AppData\Local\Programs\Python\Python310\lib\site-packages\flask\app.py", line 1825, in full_disp
atch_request
    rv = self.handle_user_exception(e)
  File "C:\Users\Raquel\AppData\Local\Programs\Python\Python310\lib\site-packages\flask\app.py", line 1823, in full_disp
atch_request
    rv = self.dispatch_request()
  File "C:\Users\Raquel\AppData\Local\Programs\Python\Python310\lib\site-packages\flask\app.py", line 1799, in dispatch_
request
    return self.ensure_sync(self.view_functions[rule.endpoint])(**view_args)
  File "C:\Users\Raquel\Desktop\Decision Making App\dec-making-app-v32\app.py", line 1102, in rank_prefs
    opt_num = int(line[0])
ValueError: invalid literal for int() with base 10: '1.'
127.0.0.1 - - [12/Sep/2023 15:18:22] "GET /prefs/reward HTTP/1.1" 500 -
```

This error tends to happen because 'pref_reward.txt' or 'pref_cost.txt' was formatted incorrectly. In this case, the 'value error' line gives us a clue on what might be wrong. The code is trying to convert the string '1.' into a number, but it can't because there is a period in the way.

To solve this, notice the story that the user encountered the error in; this is indicated in the command line window, where it reads 'Story:'. In the example above, the culprit is '/social/story_18'. For this example, we would need to navigate to '{path_to_DM_app}\stories\task_types\social\story_18\pref_reward.txt', where we encounter…

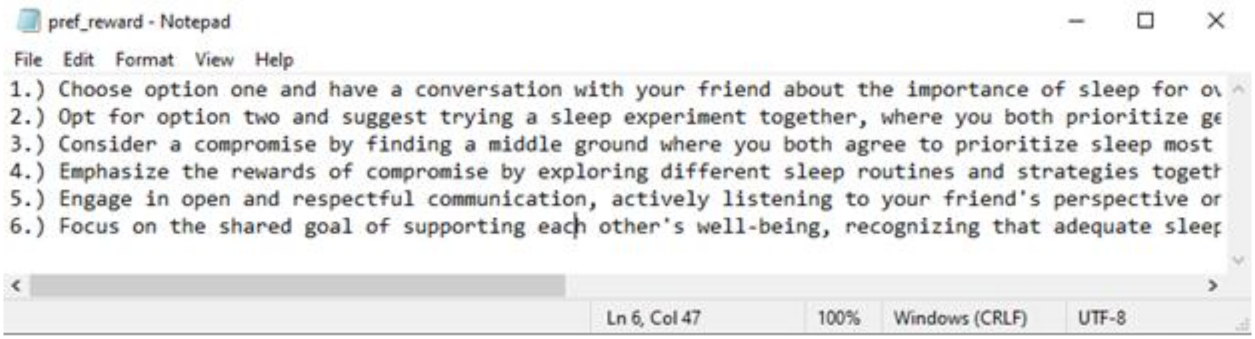

Sure enough the problem was caused by the periods after the numbers. Remove them and save...

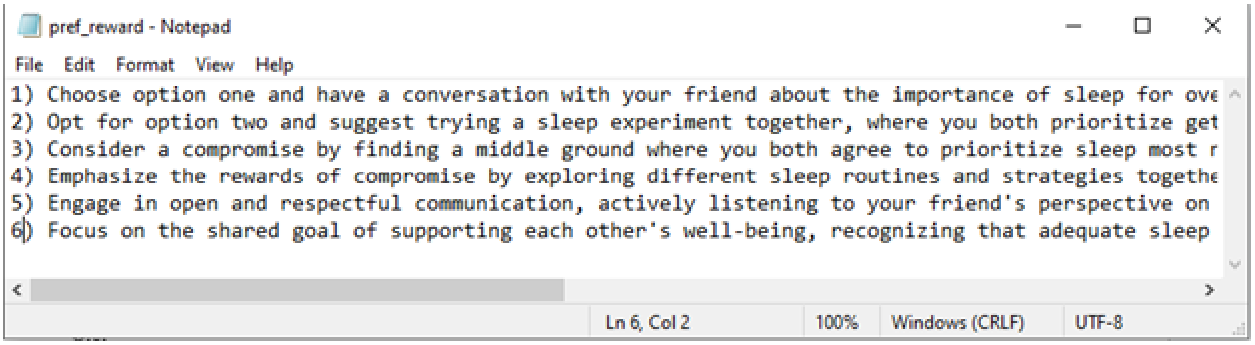

...then refresh the page the user left off on.

#### Empty preference options

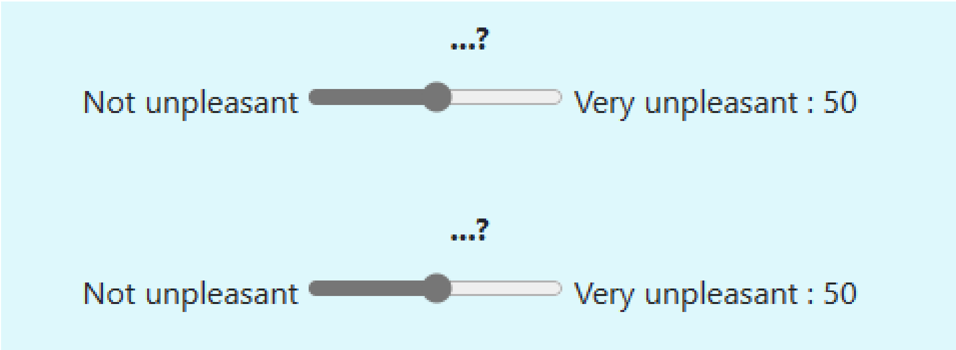

Although this error may not cause the app to halt, it may be very confusing for the participant to encounter these glitches and can negatively affect the data.

Check the command line window to figure out what story this happened in. In our example, the culprit is ‘/social/story_18’. For this example, we would need to navigate to ‘{path_to_DM_app}\stories\task_types\social\story_18\pref_reward.txt’, where we encounter...

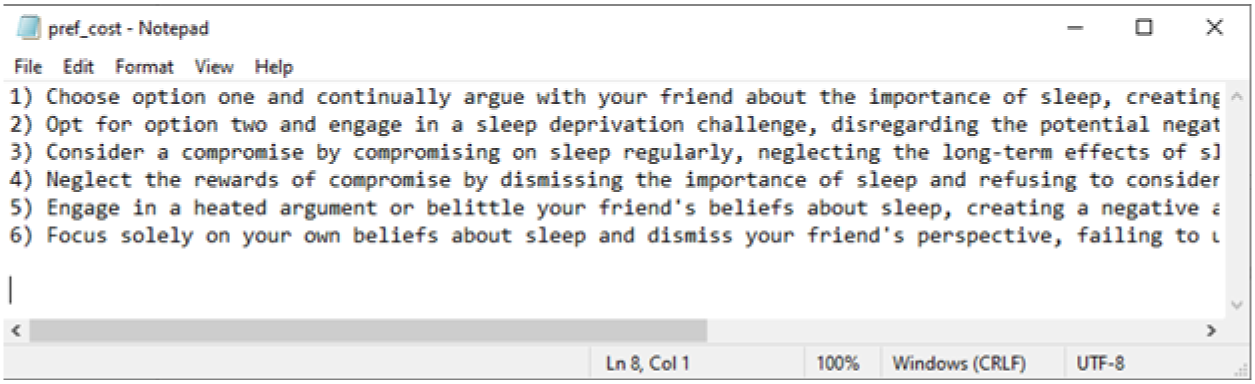

```
pref_cost - Notepad
File  Edit  Format  View  Help
1) Choose option one and continually argue with your friend about the importance of sleep, creating
2) Opt for option two and engage in a sleep deprivation challenge, disregarding the potential negat
3) Consider a compromise by compromising on sleep regularly, neglecting the long-term effects of sl
4) Neglect the rewards of compromise by dismissing the importance of sleep and refusing to consider
5) Engage in a heated argument or belittle your friend's beliefs about sleep, creating a negative a
6) Focus solely on your own beliefs about sleep and dismiss your friend's perspective, failing to u

Ln 8, Col 1    100%    Windows (CRLF)    UTF-8
```

The problem is the two empty lines at the end of the list. To solve this problem, delete these two lines so that there are no empty lines at the end of the list, save the file, then refresh.

#### IndexError: list index out of range

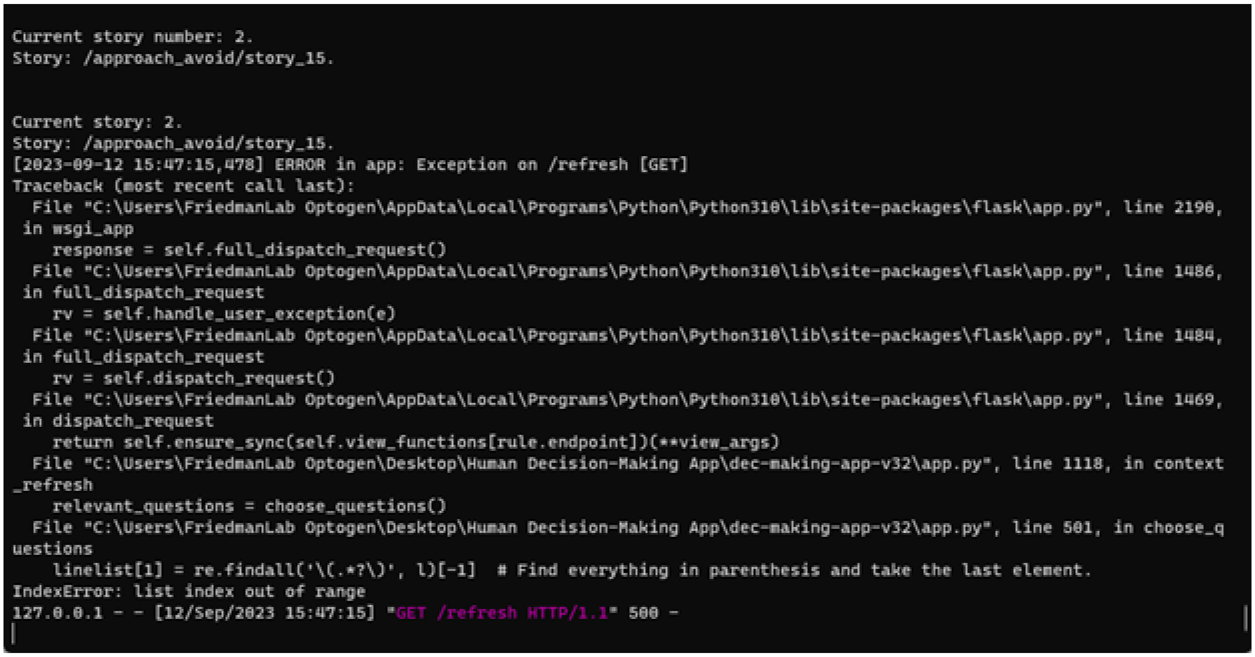

```
Current story number: 2.
Story: /approach_avoid/story_15.

Current story: 2.
Story: /approach_avoid/story_15.
[2023-09-12 15:47:15,478] ERROR in app: Exception on /refresh [GET]
Traceback (most recent call last):
  File "C:\Users\FriedmanLab Optogen\AppData\Local\Programs\Python\Python310\lib\site-packages\flask\app.py", line 2190,
 in wsgi_app
    response = self.full_dispatch_request()
  File "C:\Users\FriedmanLab Optogen\AppData\Local\Programs\Python\Python310\lib\site-packages\flask\app.py", line 1486,
 in full_dispatch_request
    rv = self.handle_user_exception(e)
  File "C:\Users\FriedmanLab Optogen\AppData\Local\Programs\Python\Python310\lib\site-packages\flask\app.py", line 1484,
 in full_dispatch_request
    rv = self.dispatch_request()
  File "C:\Users\FriedmanLab Optogen\AppData\Local\Programs\Python\Python310\lib\site-packages\flask\app.py", line 1469,
 in dispatch_request
    return self.ensure_sync(self.view_functions[rule.endpoint])(**view_args)
  File "C:\Users\FriedmanLab Optogen\Desktop\Human Decision-Making App\dec-making-app-v32\app.py", line 1118, in context
_refresh
    relevant_questions = choose_questions()
  File "C:\Users\FriedmanLab Optogen\Desktop\Human Decision-Making App\dec-making-app-v32\app.py", line 501, in choose_q
uestions
    linelist[1] = re.findall('\(.*?\)', l)[-1]  # Find everything in parenthesis and take the last element.
IndexError: list index out of range
127.0.0.1 - - [12/Sep/2023 15:47:15] "GET /refresh HTTP/1.1" 500 -
```

This error can be caused by many things. It essentially means that the code tries to access an item that does not exist within a list of variables. This could mean that the list is incomplete or empty. In the example above, the line above the 'IndexError' gives us a clue of what might have happened.

When choosing the questions that will be shown to the user for each trial, the numbers in the parentheses are used to select only the most relevant questions for the user. The code looks for anything that is enclosed in parentheses on each line of the text and separates it from the rest. If the code is not enclosed in parenthesis, the program will not find it and thus cannot continue.

Make sure all parentheses are closed, save the file, and refresh the page.

##### Missing question during trials

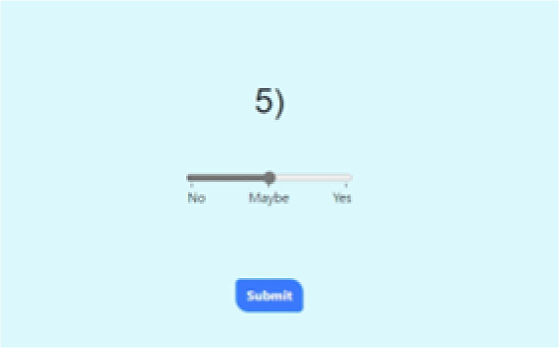

This error does not cause an internal server error, however, it may be confusing for the user and can confound the data. The error is caused because the question was not found in the ‘questions.txt’ file.

To find the question, the program loads the ‘questions.txt’ file and looks through each line of the file searching for sentences. It looks for a capitalised word and continues scanning the sentence until it finds some kind of punctuation (a period, a question mark, or an exclamation mark). If one or both of these conditions isn’t met, then the program will not be able to recognise a sentence as a question and won’t have anything to display to the user.

To solve this, make note of which story caused the issue, find this by looking at the command line and find the most recent line that starts with ‘Story:’. Navigate to ‘{path_to_DM_app}\stories\task_types\{target_story}\’, open ‘questions.txt’, and check that all sentences start with a capitalised word and end with punctuation. Save the file, **but don’t refresh the page as this may cause data loss.**

After saving, you may proceed in one of two ways; please refer to the section ‘Recovering from catastrophic failures’.

#### Problems with story sampling

##### *Raised exception: Sample larger than population or is negative*

```
Task type: Probability
Stories in pool: ['14', '1', '18']
Length of story pool: 3
Attempting to sample: 0

Generated story pool for Cost-Cost including topic Vehicle/Transportation: ['1', '11']
Generated story pool for Cost-Cost including topic Medical: ['1', '11', '5', '12']
Generated story pool for Cost-Cost including topic Party: ['1', '11', '5', '12', '6', '10']
Generated story pool for Cost-Cost including topic Entertainment: ['1', '11', '5', '12', '6', '10', '2']

Task type: Cost-Cost
Stories in pool: ['1', '11', '5', '12', '6', '10', '2']
Length of story pool: 7
Attempting to sample: 12

Could not sample stories for the selected task type 'Cost-Cost', this is usually because the amount of stories to
sample for this exceeds the number of stories available for the task type. Please make sure there are enough
stories to sample from for the user's selected topics: ['Vehicle/Transportation', 'Medical', 'Party',
'Entertainment']
Raised exception: Sample larger than population or is negative
```

This error can present itself when a particular task type  does not have the amount of stories the ‘settings.ini’ file is asking the app to sample.

The app goes through each topic that the user selected, then for each task type, creates a story pool by adding the stories that fall into both those categories. Once the story pool is created, the app randomly samples a certain amount of stories for each task type. The amount of stories that get sampled is the number indicated by the approach_avoid, benefit_benefit, cost_cost, moral, multi_choice, probability, and social (lines 51 - 57) settings in ‘settings.ini’.

To solve this issue, first identify which task type is causing the problem by looking at what was printed to the command window before the error occurred. The command window will break down the process of sampling stories. The last ‘Task type:’ line will tell you which task type caused the problem, and the different topics where stories were sampled from will be indicated in the preceding lines. If the length of the resulting story pool is lesser than the amount of stories that the app attempts to sample, this error will be raised.

For the example above, we can see that the task type that caused the issue was ‘Cost-Cost’. If we look at the breakdown of how the story pool was generated, we can see that the generated pool ended up being smaller than the amount to be sampled (7 vs. 12).

Solving this problem can be tricky, until more content will be added to the app, here is a temporary solution…

1. Stop the app (close the command window). The user’s data will likely need to be discarded.
2. The amount of stories that get sampled is the number indicated by the approach_avoid, benefit_benefit, cost_cost, moral, multi_choice, probability, and social (lines 51 - 57) settings in ‘settings.ini’. Decrease the appropriate values to however large the story pool

ended up being.

3. [Restart the app](#).
4. Repeat the user setup process.

#### Problems with database

Problems with the database can arise when the host, database, port, user, or password parameters are not set correctly in ‘settings.bin’. Here are a few problems that you may encounter:

##### *psycopg2.OperationalError: could not translate host name "local_host" to address: Unknown host*

```
  Checking if table 'human_dec_making_table_2' exists in database...
Traceback (most recent call last):
  File "C:\Users\Raquel\AppData\Local\Programs\Python\Python310\lib\runpy.py", line 196, in _run_module_as_main
    return _run_code(code, main_globals, None,
  File "C:\Users\Raquel\AppData\Local\Programs\Python\Python310\lib\runpy.py", line 86, in _run_code
    exec(code, run_globals)
  File "C:\Users\Raquel\AppData\Local\Programs\Python\Python310\lib\site-packages\flask\__main__.py", line 3, in <module
>
    main()
  File "C:\Users\Raquel\AppData\Local\Programs\Python\Python310\lib\site-packages\flask\cli.py", line 1050, in main
    cli.main()
  File "C:\Users\Raquel\AppData\Local\Programs\Python\Python310\lib\site-packages\click\core.py", line 1055, in main
    rv = self.invoke(ctx)
  File "C:\Users\Raquel\AppData\Local\Programs\Python\Python310\lib\site-packages\click\core.py", line 1657, in invoke
    return _process_result(sub_ctx.command.invoke(sub_ctx))
  File "C:\Users\Raquel\AppData\Local\Programs\Python\Python310\lib\site-packages\click\core.py", line 1404, in invoke
    return ctx.invoke(self.callback, **ctx.params)
  File "C:\Users\Raquel\AppData\Local\Programs\Python\Python310\lib\site-packages\click\core.py", line 760, in invoke
    return __callback(*args, **kwargs)
  File "C:\Users\Raquel\AppData\Local\Programs\Python\Python310\lib\site-packages\click\decorators.py", line 84, in new_
func
    return ctx.invoke(f, obj, *args, **kwargs)
  File "C:\Users\Raquel\AppData\Local\Programs\Python\Python310\lib\site-packages\click\core.py", line 760, in invoke
    return __callback(*args, **kwargs)
  File "C:\Users\Raquel\AppData\Local\Programs\Python\Python310\lib\site-packages\flask\cli.py", line 911, in run_comman
d
    raise e from None
  File "C:\Users\Raquel\AppData\Local\Programs\Python\Python310\lib\site-packages\flask\cli.py", line 897, in run_comman
d
    app = info.load_app()
  File "C:\Users\Raquel\AppData\Local\Programs\Python\Python310\lib\site-packages\flask\cli.py", line 312, in load_app
    app = locate_app(import_name, None, raise_if_not_found=False)
  File "C:\Users\Raquel\AppData\Local\Programs\Python\Python310\lib\site-packages\flask\cli.py", line 218, in locate_app

    __import__(module_name)
  File "C:\Users\Raquel\Desktop\Decision Making App\dec-making-app-v32.1\app.py", line 944, in <module>
    if (not exists(app_settings['data_table'])) and app_settings['auto_create_table']:
  File "C:\Users\Raquel\Desktop\Decision Making App\dec-making-app-v32.1\app.py", line 578, in exists
    conn = psycopg2.connect(**server)
  File "C:\Users\Raquel\AppData\Local\Programs\Python\Python310\lib\site-packages\psycopg2\__init__.py", line 122, in co
nnect
    conn = _connect(dsn, connection_factory=connection_factory, **kwasync)
psycopg2.OperationalError: could not translate host name "local_host" to address: Unknown host

PS C:\Users\Raquel\Desktop\Decision Making App\dec-making-app-v32.1> _
```

This error arises when the ‘host’ parameter is set to something that cannot be recognised by Python. In this case, ‘local_host’ was entered when the correct term would be ‘localhost’.

#### Problems with missing peripherals

Before starting the app, please ensure that both the eye tracker and the heart rate monitor antenna are properly connected to the host computer. If problems with hardware persist, please consult the device company’s support resources.

#### Problems with missing or misnamed files/directories

Missing file problems tend to arise when the code tries to access some external file to read or write, but does not find anything from the file path that was specified. Thus, the file structure of the app is extremely important; all files and folders must have the correct names, otherwise these errors may arise.

*FileNotFoundError: [Errno 2] No such file or directory*

```
Starting story /moral/story_24
Current topic: Trip (5)

127.0.0.1 - - [19/Oct/2023 15:57:54] "GET /story_num_overall HTTP/1.1" 200 -
127.0.0.1 - - [19/Oct/2023 15:57:54] "GET /static/main.css HTTP/1.1" 304 -

Current story number: 11.
Story: /moral/story_24.

[2023-10-19 15:58:09,710] ERROR in app: Exception on /context [GET]
Traceback (most recent call last):
  File "C:\Users\Raquel\AppData\Local\Programs\Python\Python310\lib\site-packages\flask\app.py", line 2528, in w
sgi_app
    response = self.full_dispatch_request()
  File "C:\Users\Raquel\AppData\Local\Programs\Python\Python310\lib\site-packages\flask\app.py", line 1825, in f
ull_dispatch_request
    rv = self.handle_user_exception(e)
  File "C:\Users\Raquel\AppData\Local\Programs\Python\Python310\lib\site-packages\flask\app.py", line 1823, in f
ull_dispatch_request
    rv = self.dispatch_request()
  File "C:\Users\Raquel\AppData\Local\Programs\Python\Python310\lib\site-packages\flask\app.py", line 1799, in d
ispatch_request
    return self.ensure_sync(self.view_functions[rule.endpoint])(**view_args)
  File "C:\Users\Raquel\Desktop\Decision Making App\dec-making-app-v32.2\app.py", line 1261, in context
    txt = open(path).read().replace("’", "'")
FileNotFoundError: [Errno 2] No such file or directory: 'stories/task_types/moral/story_24/context.txt'
127.0.0.1 - - [19/Oct/2023 15:58:09] "GET /context HTTP/1.1" 500 -
```

This error has to do with a particular file not being accessible because it either does not exist, or the name of the file is not what is expected. In the example above, the code tried to find a context file using the path ‘stories/task_types/moral/story_24/context.txt’. If we navigate to this directory, we will find that the context file is misnamed.

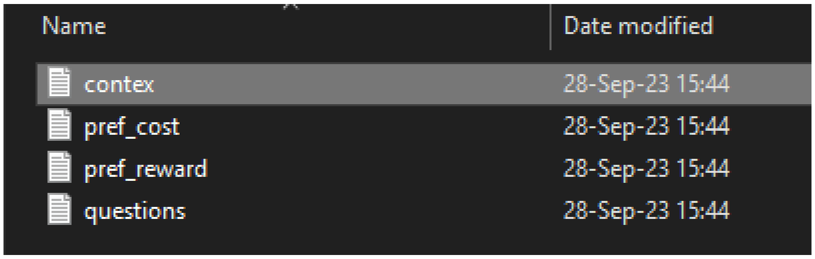

To fix this, simply rename the file to what the code expects. Most of the time, one can recover from this error by refreshing the page, but in some cases, a full restart may be necessary.

*KeyError: ‘topic’*

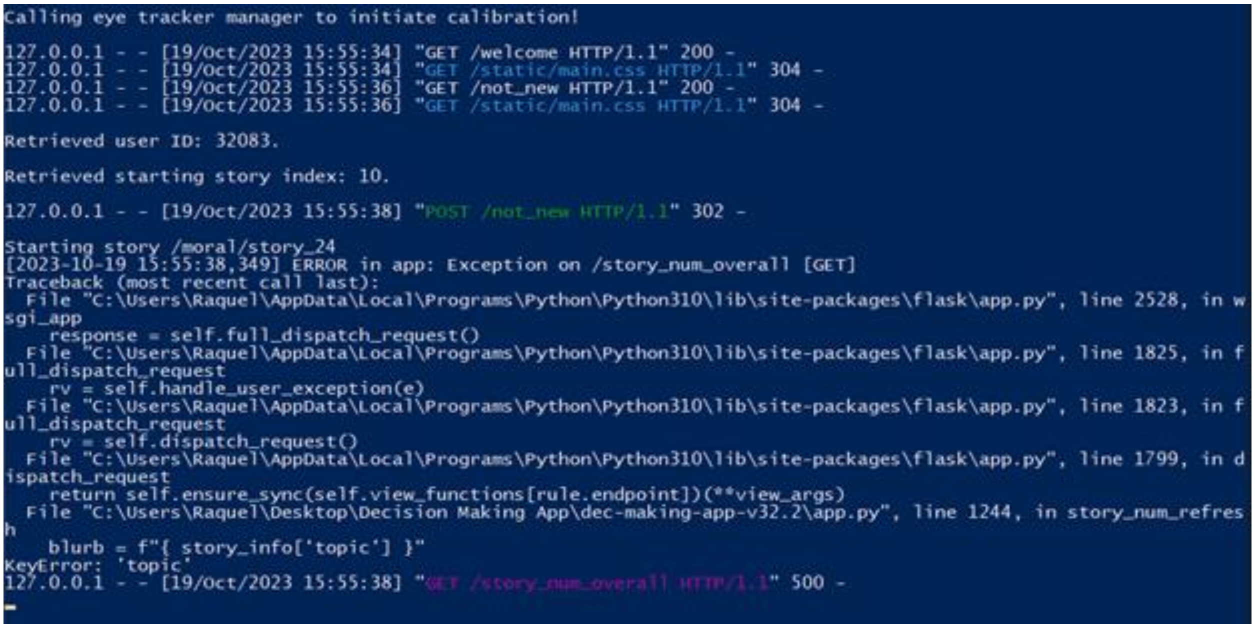

This error is much more obscure and may not have anything to do with file or directory names. However, if you find that this error occurs at the start of a story, when the story context is supposed to appear, the cause of the error could be a misnamed folder/directory.

To identify the problem, refer to the command line printout that identifies the story that was last attempted to be loaded. In the example above, we see '/moral/story_24'. If we navigate to the directory that contains all the 'moral' tasks, we find the following:

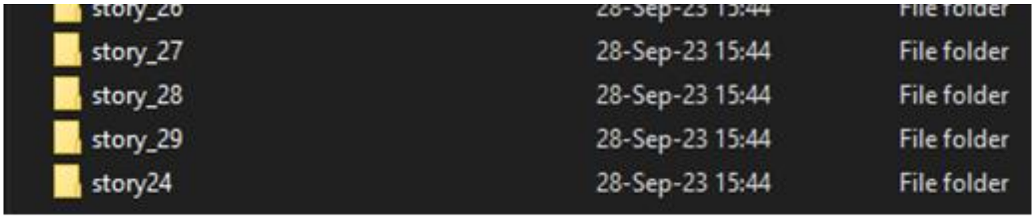

In this case, the 24th story directory is misnamed. Story directories must be named 'story_{story number}' in order for the code to recognise them as such. To fix this issue, we rename this directory to 'story_24'.

Unfortunately, this error occurred at a point where refreshing the page will not recover the code from the failure. An app restart is needed in this case. Refer to the '[Recovering from catastrophic failures](#)' section for steps to recover from this.

#### Recovering from catastrophic failures

Catastrophic failures occur when the code raises an exception that cannot be recovered from by refreshing the page. This may be because some critical data was pre-loaded into memory and cannot be re-created using new information, or because external hardware prevents the code from re-running. A catastrophic failure may also occur when the code soft fails (does not stop running), but shows incorrect information to the user, which can affect data.

What measures to take will depend on your unique situation and whether data may be potentially lost or 'corrupted'. If there is potential data loss during a trial, follow either **solution 1** or **solution 2**. If there is no expected data loss (for example, the error did not occur using a trial question), proceed to **solution 2**.

**Solution 1**
You may want to simply ask the subject to skip a question altogether and continue through the rest of the trials, however, the subject may encounter the same error again if there was more than one question that had the same issue in the 'questions.txt' file. If you choose to proceed with the trials, make note of which questions had a problem and write them down in the session notes at the end of the session.

The app will record the subject's data and immediately upload it to the database whenever the subject clicks the 'Submit' button during a trial, so there will be some data that can be preserved and analysed.

**Solution 2**
You might decide that because there were errors in the session, the data will be bad and unusable in its entirety, or you might not be able to recover from the error by simply refreshing the page. If this is the case, you may interrupt the session and ask the subject to re-do the problematic story after you've troubleshooted it. To do this, follow the following steps:

1. Bring the command line window up and enter the keyboard combination 'Ctrl + C'. If you're prompted to confirm, enter 'Y' and press the Enter key.
2. Make sure you make any necessary corrections to the 'questions.txt' file and navigate to '\bin\settings.ini'. Open this file with your text editor of choice (f.e. Notepad, or Notepad++). Make the following changes:
    a. (line 34) next_story_from=local
3. Save the settings file and minimise it. Restart the app as you did before. Ask the subject to select however many stories they had left in the previous session and run them through the same setup as before. When asked 'Are you a new participant?', ask the subject to click 'No' and enter their ID number.
4. The app should continue on the story where it left off before the error happened.
5. Once the subject is done with their session, make sure to add a note of what happened to the session notes.
6. Stop the app through the command line as in step 1, undo the changes you made to settings, and restart the app.

Following these steps should minimise data loss.

#### Adding or replacing stories in participant’s story order

Sometimes it may be necessary to add or replace stories to an existing participant’s story order. This may be because some new task types were included into the study, in which case a new story order including the new task types needs to be added. It could also be that a particular story needs to be removed or replaced from a participant’s story order. In both of these cases, one can directly manipulate the stories that the subject will see.

##### *Adding stories*

To generate a new story order, refer to the distribute_stories.py script. This script runs directly from the DM app’s root directory and references the ‘settings.ini’ file contained in the ‘bin’ directory.

Distribute_stories.py is the same algorithm that the dm app uses to create a user's story order upon beginning their first-ever session. It is a command line script that will take a list of at least X topics, where X is defined as the minimum amount of stories that users are required to select from the dm app; this is defined by the 'minimum_topics' parameter in 'bin\settings.ini'. If no parameters are entered, the script will use a hard-coded list of topics as an execution example. The script will fail if less than X topics are passed. You may pass more than X topics at a time, as long as they exist in the 'Human DM Topics.xlsx' file.

Assuming no errors are found in the code execution, this script will spit out a freshly-made story order following the same instructions the 'settings.ini' file gives. This means that you must modify and save the appropriate parameters in 'settings.ini', this is a list of parameters that will affect the outcome of this script:

1. approach_avoid,
2. benefit_benefit,
3. cost_cost,
4. moral,
5. multi_choice,
6. probability,
7. social,

All of which indicate how many stories to pull from each task type, and...

8. minimum_topics

To specify which topics to pick stories from, add them as command line arguments by writing them one-by-one after 'python distribute_stories.py ', each inside single quotes and separated by a space (see example below).

**USAGE EXAMPLE**

> in 'settings.ini': (`approach_avoid=12, benefit_benefit=0, cost_cost=0, moral=0, multi_choice=0, probability=0 social=12, and minimum_topics=4`)

Write in command line:

```
python distribute_stories.py 'Education-Post-Education Life' 'Food'
'Vehicle/Transportation' 'Entertainment'
```

Output:

```
['/social/story_20', '/approach_avoid/story_16', '/social/story_23',
'/approach_avoid/story_14', '/social/story_24',
'/social/story_7', '/social/story_8', '/approach_avoid/story_9',
'/social/story_11', '/approach_avoid/story_19',
'/social/story_12', '/approach_avoid/story_24',
'/approach_avoid/story_15', '/approach_avoid/story_20',
'/social/story_6',
'/approach_avoid/story_2', '/approach_avoid/story_10',
'/social/story_22', '/social/story_3', '/approach_avoid/story_6',
'/approach_avoid/story_21', '/social/story_10', '/social/story_21',
'/approach_avoid/story_5']
```

Keep in mind that the topics must be written exactly as they are in the 'Human DM Topics.xlsx' file!

If you are generating an additional story order for a particular user, access their demographic data by navigating to '\data\{user_id}\demographic_info.txt', and reference the 'pref_stories' line.

Once the new story order is generated, copy it from the command line window and **append** it to the **end** of the user's 'story_order' line. Do not delete the existing story order!!

When pasting, make sure to delete the original closing square bracket (']'), add a comma and a space, and paste the new story order without the opening square bracket ('['). Also make sure you have not accidentally added an empty line to the end of the file.

Finally, save the demographics info file and close it. The app will continue from the last story it left off on. This is indicated by the user's 'next_story_index' (if 'next_story_from=local'), or by the number of different stories gone through registered in the database (if '` next_story_from=database` ').

##### *Replacing stories*

To replace a story in the user's story order, navigate to the 'data' directory from the HMDA root directory, find the user's ID number, and open their 'demografic_info.txt' file. Under the 'story_order' line, replace whichever story you mean to replace, make sure the new story does exist in the 'stories/task_types' folder. Include the task type name and the story number, for example, '/social/story_5'.

If you need the user to go back a few stories to do the new story, modify the 'next_story_index' line by subtracting however many stories you need so that the user sees the story. Remember that 'next_story_index' is the index value of the next story in the story order list, where the first element in the list is element number 0.

Make sure you also set the 'next_story_from' parameter to 'local' so that the app references this number in the demographic data and does not look into the database to retrieve the next story index.

##### *Removing stories*

To remove a story from the user's story order, navigate to the 'data' directory from the HMDA root directory, find the user's ID number, and open their 'demografic_info.txt' file. Under the 'story_order' line, remove whichever story you need to remove.

You may need to modify the 'next_story_index' line if the removal impacts which story the user will see next. For example...
Say you need to remove '/social/story_7' from the following story order.

```
['/probability/story_3', '/multi_choice/story_11', '/moral/story_23',
'/multi_choice/story_1', '/social/story_7', '/cost_cost/story_2',
'/benefit_benefit/story_10', '/moral/story_15',
'/approach_avoid/story_7', '/probability/story_14', '/social/story_4',
'/social/story_17']
```

If we count the elements in the list starting from 0, we can figure out the index number (shown in superscript) for each story in the list…

['/probability/story_3'$^{0}$, '/multi_choice/story_11'$^{1}$,
'/moral/story_23'$^{2}$, '/multi_choice/story_1'$^{3}$, '/social/story_7'$^{4}$,
'/cost_cost/story_2'$^{5}$, '/benefit_benefit/story_10'$^{6}$,
'/moral/story_15'$^{7}$, '/approach_avoid/story_7'$^{8}$,
'/probability/story_14'$^{9}$, '/social/story_4'$^{10}$, '/social/story_17'$^{11}$]

Now, imagine that next_story_index: 6, which indicates that the user's next story will be

'/benefit_benefit/story_10', the seventh element in the list.

If you simply remove ‘/social/story_7’ from the list, index 6 would point to '/moral/story_15' instead, which would mean that the user would never see ‘/benefit_benefit/story_10’.

['/probability/story_3'[0], '/multi_choice/story_11'[1], '/moral/story_23'[2], '/multi_choice/story_1'[3], '/cost_cost/story_2'[4], '/benefit_benefit/story_10'[5], '/moral/story_15'[6], '/approach_avoid/story_7'[7], '/probability/story_14'[8], '/social/story_4'[9], '/social/story_17'[10]]

In this case, in order for the user to see '/benefit_benefit/story_10', we would need to change next_story_index to 5.

Make sure you also set the 'next_story_from’ parameter to ‘local’ so that the app references next_story_index in the demographic data and does not look into the database to retrieve the next story index.

### Eye Tracking Study Considerations

A full version of these guidelines can be found on the official Tobii Connect website. The information contained in this section was paraphrased or taken directly from the documentation provided there. See the sources section for links to the original documentation.

#### Participant Screening

Because corrective lenses (glasses or contacts) might cause issues with the eye tracker during our trials, it might be good to screen for the following people:

- Wearers of corrective lenses (glasses) for cataracts, as well as bifocals
- Individuals who have had eye surgeries: corneal, cataracts, intraocular implants
- Individuals with eye movement or alignment abnormalities: lazy eye, nystagmus, others
- If participants can wear contact lenses instead of glasses, please advise them to do so

#### Eye Tracker limitations

The eye tracker will not usually have a problem with corrective lenses; however issues may arise if:

- there are internal reflections being caused by lighting in the room,
- the glasses themselves keep moving on the participant,
- the lens correction is very strong (+/- 6 or more),
- or the frames somehow occlude the eye-image in the camera (a simple adjustment of the angle of the tracker can usually remedy this).

The eye tracker will not cope with bi- or varifocal lenses and this is because the lens distorts the shape of the pupil which causes detection errors in the eye tracking software. So, when recruiting, it's important that you always include questions about glasses in the screening and *ensure participants are wearing the appropriate pair for the task BEFORE calibrating*. Oh, and yes, offer them a lens cloth before you start calibration if the glasses look a bit grubby!

Contact lenses also come in bi-focal/varifocal form these days. So, whilst contact lenses are usually OK with all kinds of eye-trackers, the multi-prescription lenses CAN cause problems, so it's always worth checking!

#### Tips for facilitating eye tracking studies with people who wear corrective lenses – and even those who don't...

***Corrective glasses should be worn during calibration!!***

*If you see your participant squinting, it's almost certainly going to cause issues for the tracker* and it suggests their vision is not corrected, so **keep an eye on your participant during data collection.**

If you have access to one, use a Snellen or LogMAR chart at the distance of your stimuli to test your participant's vision before you start. You're not an optician and you're not going to diagnose their condition, but this is a great way to check if their vision is as good as they claim it to be and so they should be wearing their lenses whilst doing this!

**Show them some example images/text on screen and get them to describe them to you or read out loud and you'll soon know if you're going to have a problem – because they'll squint, lean in to the screen or simply not be able to do it.** If you can, have the eye tracker on for this to test how well the eyes are being detected.
If you're using a desktop tracker, have a range of cheap reading glasses, covering different corrections – ideally from -6 to +6 diopters. When the inevitable happens and someone forgets their glasses you can offer them as a temporary solution and avoid having to discard the participant. These are typically available from large supermarkets or chemists.

#### Important Reminders

Remind your participants for the day of the eye tracking study:

- Avoid wearing heavy make-up, mascara, false eye lashes or colored contact lenses.
- If doing 'intercept' recruitment, keep an eye out for the above in addition to physical features (such as unusually droopy eyelids) that may prevent the eye tracking sensors from 'seeing' the eyes.
- Check and confirm that all participants have read and agreed to the consent form!

Don't forget to inspect your equipment:

- Your eye tracker is cleaned and ready with any accessories.
- Your computer or tablet is up to date, with all required software functioning as expected.
- You have additional accessories such as power cables, chargers, extensions, cleaning cloth and sanitary wipes, external storage drives, and any required dongles or adapters.
- Perform a test run of the data collection, exactly like you would for the participants, to make sure your setup and environment are up to expectations.

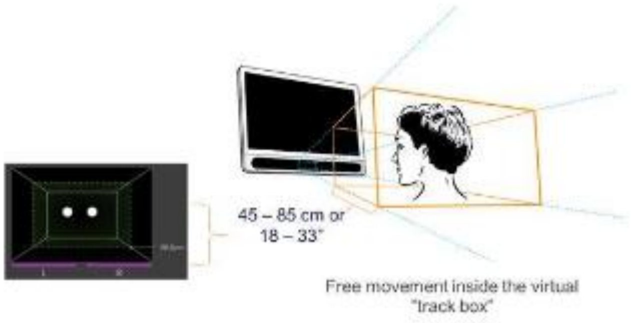

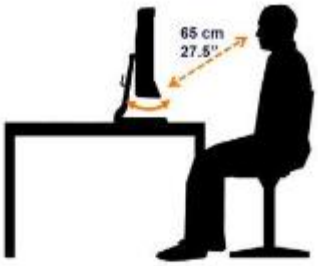

#### Positioning Participants

- Make sure your participant is comfortable, has a good position relative to the screen, and can reach the keyboard/mouse if used.
- You can check the participant's head's range of movement using the track box guide. It is available on the top left before starting a recording in Tobii Pro Lab.
- Use of a chair without wheels and swivelling is recommended as it can limit how far the participant can move from the track box.

#### Running the Test

1. After positioning your participant in front of the screen, start a recording in Tobii Pro Lab.
2. Since all eyes are different, you will first run a calibration.
3. During the calibration, ensure the participant remains focused; avoid conversation or laughter.

#### Interpret the results

- Measurement points should be within the cross' boundaries for best accuracy
- The bigger the distance of the points to the cross' centre, the larger the potential measurement error
- The further the elements on your stimulus are separated, the larger error can be accepted
- For target groups with difficult calibration, consider modifying the stimulus

#### Recording

After successful calibration, you may start the recording.

- Explain the task slowly and thoroughly to the participant and make sure they know what they are supposed to do.
- During the task, you as a data collector should only observe. Let the participant complete the task in a natural way and only interrupt if necessary. Take notes of relevant behaviour and observations during the test - this is not only be useful for post-interviews but also analysing and interpreting your data.
- After the presented tasks are complete, stop and save the recording.
- Perform the post-interview (optionally with retrospective think aloud)
- Don't forget the consent form!

##### Section Sources

This entire section mirrors the content of the following websites:

1. https://connect.tobii.com/s/field-guide-screen-based?language=en_US&t=1693606963655
2. https://www.tobii.com/blog/eye-tracking-study-recruitment-managing-participants-with-vision-irregularities

#### App settings

The HUMANS app accesses a list of parameters located in the '/bin/settings.ini' configuration file (INI). Below is a detailed explanation of what each parameter does and where it is accessed in the source code.

| Parameter | Section | Type | Location | Description |
|---|---|---|---|---|
| host | postgresql | str | | The host computer's IPv4 address or domain name. If HUMANS is running on the host computer, enter the word localhost. |
| database | postgresql | str | | The name of your PostgreSQL database. |
| port | postgresql | int | | The host computer's port-forwarded port for servicing database queries (PostgreSQL defaults to port 5432). |
| user | postgresql | str | | The PostgreSQL username. Keep in mind that the account that the HMDA uses must have database read-write privileges. |
| password | postgresql | str | | The password tied to the account identified by the 'user' parameter. |
| data_table | app_settings | str | | The name of the data table structure defined in the PostgreSQL database. Data |

| | | | | |
|---|---|---|---|---|
| | | | | generated by the HUMANS app will be stored here. If a data table has not already been created, refer to the ‘auto_create_table’ parameter below. |
| auto_create_table | app_settings | bool | | If the table specified above does not exist in the database, the app may be able to create it using the participant's data. If this parameter is set to 1, the app will attempt to create the target data table once it has all the required data from the user If set to 0, the app will not upload data to the database. |
| enable_consecutive_users | app_settings | bool | | Enables back-to-back use of the app without the need to close and re-run startup.bat (which restarts the flask server). If set to 1, the app will reset all global app and user parameters to default when loading the landing portal (setup_session.html, routed as '/'). If this option is not enabled (set to 0), the flask server must be closed and re-run. Any changes to the source code or templates are not affected by this option. |
| data_upload | app_settings | bool | | Ensures that data is automatically uploaded to the database after each session if set to 1. When set to 0, the write_trial_to_database routine is silently skipped. |
| unique_ids_from | app_settings | str | | Controls where IDs are read from when trying to generate a unique ID for each user. Possible values are "database" and "local".<br>∉ "database" will look into the database and find all the unique IDs stored there.<br>∉ "local" will only look to the "/data/" directory to find all the taken IDs. The app will then generate a new random ID that does not exist in the retrieved list of existing IDs. |
| next_story_from | app_settings | str | | Controls where the next story index is referenced from. Possible values are "database" and "local". |

| | | | | |
|---|---|---|---|---|
| | | | | ∉ "database" will look into the database to find the next story the user will see by counting all the unique entries of 'tasktypedone'.<br>∉ "local" will look into the user's local demographic info record in '/data/' and read the 'next_story_index' entry. |
| timestamp_timezone | app_settings | str | | The time zone that timestamps are collected in. 'UTC' is recommended. If this parameter is not set, the app will default to 'UTC'. |
| minimum_topics | app_settings | int | | The minimum amount of topics that the subject must select. Replaces 'min_stories_to_choose' from version 30. |
| questions_per_story | app_settings | int | | The number of questions selected to show the user **per story**. |
| ignore_legacy_story_data | app_settings | bool | | How the app handles story data ('pref_stories' and 'story_order') from legacy app versions. If this is set to 0, the app will NOT ignore legacy story data from returning users and continue the remainder of the users' sessions with only 'approach_avoid' task types, keeping their preferred stories and previously calculated story order. If set to 1, the app will discard this information and ask the user to restart their sessions to add the new task types to their story data. |
| randomise_relation_levels | app_settings | bool | | FOR SOCIAL TASK ONLY: Whether to randomly choose a random relationship keyword and replace it into the text of the social task stories. If set to 1, a random relationship keyword will be picked from the list under the 'relation_levels' list parameter and replaced into every snippet of text throughout the entire story. |
| relation_levels | app_settings | list | | FOR SOCIAL TASK ONLY: The words to look for and replace in story text. The app will randomly sample a word and replace it throughout the entire story text. |
| relation_level_sto | app_settings | list | | FOR SOCIAL TASK ONLY: Stories where |

| | | | | |
|---|---|---|---|---|
| ries | | | | replacing the relationship level can be done |
| validate_stories | app_settings | bool | | Validates that the stories described in the 'Human DM Topics' relationship table are actually contained in '../stories/task_types/'. Stories that don't exist in that directory will be deleted from the pool of stories that can be selected for each subject. Setting this parameter to 1 can cause errors if the validated story pool ends up being smaller than the sample size for each task type (see below). If this is the case, check that there are enough stories for each task type in '../stories/task_types/'. Conversely, not validating the stories increases the odds of trying to access a story that does not exist, causing a different error. |
| approach_avoid | app_settings | int | | The amount of stories of the corresponding task type to sample when creating a user’s story order. |
| benefit_benefit | app_settings | int | | The amount of stories of the corresponding task type to sample when creating a user’s story order. |
| cost_cost | app_settings | int | | The amount of stories of the corresponding task type to sample when creating a user’s story order. |
| moral | app_settings | int | | The amount of stories of the corresponding task type to sample when creating a user’s story order. |
| multi_choice | app_settings | int | | The amount of stories of the corresponding task type to sample when creating a user’s story order. |
| probability | app_settings | int | | The amount of stories of the corresponding task type to sample when creating a user’s story order. |
| social | app_settings | int | | The amount of stories of the corresponding task type to sample when creating a user’s story order. |
| manager_install_ | eye_tracker | str | | The install path to the eye tracker’s |

| | | | | |
|---|---|---|---|---|
| path | | | | management program. Tobii eye trackers will install the manager in C:\Users\{HOST USERNAME}\AppData\Local\Programs\TobiiProEyeTrackerManager\TobiiProEyeTrackerManager.exe |
| subscriptions | eye_tracker | list | | A list of data streams to subscribe to. Different eye tracking devices may offer different data streams. The app’s EyeTracker class considers only 'gaze', 'openness', and 'position' as valid options. |
| eyetracker_index | eye_tracker | int | | If multiple devices are connected to the system, this number will indicate which device to use. If only one device is set up, use 0. |
| use_eyetracker | eye_tracker | bool | | Activates or deactivates the subroutines involving the eye tracker.<br>∉ If 1, the app will attempt to make use of any eye trackers connected to the host.<br>∉ If 0, all subroutines will be skipped. |
| use_external_app | hr_tracker | bool | | Determines whether an external program will be called as a subprocess for collection of heart rate data.<br>∉ 1, an external program is called for data collection (see below).<br>∉ 0, a thread will be spawned by the main application process (see last four params in this section). |
| external_app_install_path | hr_tracker | str | | The external heart rate-capturing program’s location in disk. Any program may be used as long as it is callable from this path. |
| hrtracker_index | hr_tracker | int | | If multiple devices are connected to the system, this number will indicate which device to use. If only one device is set up, use 0. |
| use_heartracker | hr_tracker | bool | | Activates or deactivates the subroutines involving the heart rate tracker.<br>∉ If 1, the app will attempt to make use of any heart rate monitor devices connected to the host. |

| | | | | |
|---|---|---|---|---|
| | | | | ∉ If 0, all subroutines will be skipped. |
| emulate_device | hr_tracker | bool | | Whether to emulate a heart rate device for the heart rate monitor thread. This is mainly useful for development. Use this if you're testing the app and don't have a heart rate monitor device connected to the computer. Emulated data is also packed and sent to the database for troubleshooting purposes. |
| run_thread_as_daemon | hr_tracker | bool | | Whether to run the heart rate monitor thread as a daemon. If set to 1, the heart monitor thread will run alongside the app as a daemon. This ensures that the thread will exit along with the app, however, this could cause problems if the thread is not manually stopped |
| verbose | hr_tracker | bool | | Toggles verbose output of the hr monitor thread. This has little impact on functionality, but is useful when debugging the heart rate monitor. Setting this to 1 will allow the hr monitor thread to display messages on the same console window the app is running on. |
| test_on_startup | hr_tracker | bool | | Whether to test the heart rate monitor when the app starts.<br>∉ 1, the device will be tested by initialising and collecting data for 3 seconds.<br>∉ 0, device will not be tested on startup. |

### Database Guide

#### How to Create a Local Copy of Database

#### Notes on Environment

1. The PostgreSQL version used is PostgreSQL 14.1.
2. All code is written in MATLAB, and additional toolboxes may be necessary such as the Matlab Database Explorer and Matlab Statistics and Machine Learning Toolbox.
3. The database should only be initialized on a computer with at least 16GB of RAM and 256GB of storage.
4. Windows machines were used for all development.

#### Important Links

1. PostgreSQL: https://www.enterprisedb.com/downloads/postgres-postgresql-downloads
2. MATLAB: https://www.mathworks.com/products/matlab.html
3. MATLAB Database Explorer: https://www.mathworks.com/products/database.html
4. MATLAB Statistics and Machine Learning Toolbox: https://www.mathworks.com/products/statistics.html
5. Database Backup: https://doi.org/10.7910/DVN/OZARPL

#### Steps for Setup

1. Ensure that you have PostgreSQL and MATLAB properly installed and have all the variable paths set on your windows machine.
2. Download a copy of the database backup, located at the following link: https://doi.org/10.7910/DVN/OZARPL
3. Unzip the file into a file on your local machine. It should be a .tar file.
4. Open the windows command line (ensure that you open it as an administrator).
5. Connect to the PostgreSQL server, using the following command:

```
psql -U postgres
```

6. Create a new empty database, using the following command:

```
create database live_database
```

7. Exist psql, using the command “exit”.
8. Restore the database from the backup .tar file, modified to suit local file paths, using the following command:

```
pg_restore -U postgres -d live_database file_path_of_tar_file
```

9. Connect to the live_database using:

```
psql -U postgres -d live_database
```

10. Tables can be listed using “/dt”. If successful, the following list of tables should appear:

```
                                 List of relations
 Schema |                           Name                            | Type  |  Owner
--------+-----------------------------------------------------------+-------+----------
 public | alcoholreactiontimepsychometricfunctions                  | table | postgres
 public | alcoholrewardchoicepsychometricfunctions                  | table | postgres
 public | basepsychometricfunctions                                 | table | postgres
 public | bonsai_table                                              | table | postgres
 public | dummyrattable                                             | table | postgres
 public | featuretable                                              | table | postgres
 public | featuretable2                                             | table | postgres
 public | featuretable3                                             | table | postgres
 public | fooddeprivationreactiontimepsychometricfunctions          | table | postgres
 public | fooddeprivationrewardchoicepsychometricfunctions          | table | postgres
 public | fooddeprivationrotationpointspsychometricfunctions        | table | postgres
 public | fooddeprivationstoppingptspsychometricfunctions           | table | postgres
 public | fooddeprivationtravelpixelpsychometricfunctions           | table | postgres
 public | ghrelin_featuretable                                      | table | postgres
 public | ghrelinreactiontimepsychometricfunctions                  | table | postgres
 public | ghrelinrewardchoicepsychometricfunctions                  | table | postgres
 public | ghrelinrotationpointspsychometricfunctions                | table | postgres
 public | ghrelinstoppingptspsychometricfunctions                   | table | postgres
 public | ghrelintravelpixelpsychometricfunctions                   | table | postgres
 public | human_dec_making_table                                    | table | postgres
 public | human_dec_making_table_utep                               | table | postgres
 public | inscopix_table                                            | table | postgres
 public | isoflurenereactiontimepsychometricfunctions               | table | postgres
 public | isoflurenerewardchoicepsychometricfunctions               | table | postgres
 public | isoflurenerotationpointspsychometricfunctions             | table | postgres
 public | isoflurenestoppingptspsychometricfunctions                | table | postgres
 public | isoflurenetravelpixelpsychometricfunctions                | table | postgres
 public | jan13                                                     | table | postgres
 public | lg_boost_rewardchoicepsychometricfunctions                | table | postgres
 public | lg_boost_rotationpointspsychometricfunctions              | table | postgres
 public | lg_boost_stoppingpointspsychometricfunctions              | table | postgres
 public | lg_boost_travelpixelpsychometricfunctions                 | table | postgres
 public | lg_etoh_rewardchoicepsychometricfunctions                 | table | postgres
 public | lg_etoh_rotationpointspsychometricfunctions               | table | postgres
 public | lg_etoh_stoppingpointspsychometricfunctions               | table | postgres
 public | lg_etoh_travelpixelpsychometricfunctions                  | table | postgres
 public | live_table                                                | table | postgres
-- More  --
```

### Database Table Descriptions

#### Rat Table

##### *live_table*

1. Each row of the live_table represents a trial.
2. All raw data is stored in the live_table.
3. Sessions are composed of multiple trials and can be identified by unique date and id.

| Column name (varible) | data_type | Description |
|---|---|---|
| **id** | integer | An int number which represents a trial id, each trial has a unique id. |
| **gender** | character varying | A string which represents the gender of the rat |
| **birthdate** | character varying | A string which is the birthdate of the rat |
| **genotype** | character varying | A string representing the genotype of the rat. |
| **cagenumber** | character | A string representing the cage number of |

| | | |
|---|---|---|
| | varying | the rat. |
| **health** | character varying | A string representing the current experiment being performed on the rat. |
| **cagemates** | character varying | A string representing what rat is the cage mate of the current rat. |
| **experimenter** | character varying | A string representing the name of the person who ran the experiment. |
| **tasktypedone** | character varying | A string representing which task type the trial is being run on. |
| **notes** | character varying | A string which represents any notes which may be notes the uploader wishes to include. |
| **intensityofcost1** | character varying | A string which represents the light level in lux for feeder 1. |
| **intensityofcost2** | character varying | A string which represents the light level in lux for feeder 2. |
| **intensityofcost3** | character varying | A string which represents the light level in lux for feeder 3. |
| **intensityofcost4** | character varying | A string which represents the light level in lux for feeder 4. |
| **costprobability1** | character varying | Not currently used. |
| **costprobability2** | character varying | Not currently used. |
| **costprobability3** | character varying | Not currently used. |
| **costprobability4** | character varying | Not currently used. |
| **rewardconcentration1** | character varying | A string which represents the percentage of sucrose in feeder 1. |
| **rewardconcentration2** | character varying | A string which represents the percentage of sucrose in feeder 2. |
| **rewardconcentration3** | character varying | A string which represents the percentage of sucrose in feeder 3. |
| **rewardconcentration4** | character varying | A string which represents the percentage of sucrose in feeder 4. |
| **rewardvolume1** | character varying | A string which represents the volume of sucrose solution in feeder 1, represented in ul. |
| **rewardvolume2** | character varying | A string which represents the volume of sucrose solution in feeder 2, represented in ul. |
| **rewardvolume3** | character varying | A string which represents the volume of sucrose solution in feeder 3, represented in ul. |
| **rewardvolume4** | character varying | A string which represents the volume of sucrose solution in feeder 4, represented |

| | | |
|---|---|---|
| | | in ul. |
| **rewardprobability1** | character varying | Not currently used. |
| **rewardprobability2** | character varying | Not currently used. |
| **rewardprobability3** | character varying | Not currently used. |
| **rewardprobability4** | character varying | Not currently used. |
| **mazenumber** | character varying | A string which represents the maze which the rat was in during trial running. |
| **approachavoidtimestamp** | character varying | A string representing the time at which the approach avoid variable was set relative to the trial. Will only take on a value other than 0 when approach avoid is true. |
| **approachavoid** | character varying | A string which represents whether the rat approached the feeder. |
| **playstarttrialtone** | character varying | The time at which the start trial tone is recorded (in seconds from the start of the trial. |
| **presentcost** | character varying | The time at which the cost is presented during the trial (in seconds from the start of the trial) |
| **lightlevel** | character varying | A string representing the light level of the current trial. |
| **referencetime** | character varying | A string which represents the date and start time of the current trial. |
| **videostartime** | character varying | A string which represents the date and start time of the video. |
| **feeder** | character varying | A string representing what feeder was active during the trial. |
| **stoptrack** | character varying | A string which represents the trial event for when the Ethovision stops keeping track of where the rat is on the arena. A trial may be running but the rat's position won't be tracked until the track starts, and will stop being tracked at "stoptrack". |
| **trialname** | character varying | A string representing which trial of the current session the current row is. |
| **detectionsettings** | character varying | A string which represents the detection settings of the current trial. |
| **trialcontrolsettings** | character varying | A string which represents the current trial’s control settings. |
| **referenceduration** | character varying | A string which represents the |

| | | |
|---|---|---|
| **animalid** | character varying | A string which represents each animal's unique ids. |
| **mazeofferdelivery** | character varying | A string which represents |
| **mazenoofferdelivery** | character varying | A string which represents |
| **starttime** | character varying | The string of the date and time of the trial start. |
| **recordingafter** | character varying | A string which represents how long the recording kept going after the trial. |
| **recordingduration** | character varying | A string representing how long the recording is in seconds. |
| **trialduration** | character varying | A string representing how long the trial is in seconds. |
| **mazecostoff** | character varying | A string representing the time when the cost light is turned off, this can be either after sucrose is dispensed, or if the rat did not approach the offer. |
| **coordinatetimes** | ARRAY | No longer used. |
| **xcoordinates** | ARRAY | No longer used. |
| **ycoordinates** | ARRAY | No longer used. |
| **presentcostend** | character varying | A string representing the time from the beginning of the trial at which point the cost ends, in seconds. |
| **costpresenetationfinish** | character varying | A string representing the time in seconds from the beginning of the trial at which the cost stops |
| **stopincopixrecording** | character varying | A string representing the time in seconds when a TTL signal is sent to the calcium imagining system (inscopix) to signal it to stop recording. |
| **decisionmakingtime** | character varying | A string representing the time in seconds given to the rat to make a decision. |
| **startincopixrecording** | character varying | The time in seconds when inscopix starts recording. Will always be 0. |
| **date** | character varying | The date which the session was run. |
| **activezonetimestamp** | character varying | The time in seconds from the beginning of the trial which the active zone variable changes. |
| **activezonevalue** | character varying | ?? |
| **presentcostontimestamp** | character varying | A string representing the time, in seconds, at which the cost (light) is turned on. |
| **costacknowledgementtimestamp** | character varying | Time of the acknowledgement signal for the cost from the microcontroller. |

| | | |
|---|---|---|
| **deliveryacknowledgementtimestamp** | character varying | Time of acknowledgement signal for the delivery from the microcontroller. |
| **coordinatetimes2** | ARRAY | A string array of coordinate times for the current trial. |
| **xcoordinates2** | ARRAY | A string array of the x coordinates of the rat's center. |
| **ycoordinates2** | ARRAY | A string array of the y coordinates of the rat's center. |
| **subjectid** | character varying | A string which represents the name of the rat running the current trial. |
| **ytail** | ARRAY | No longer used. |
| **xnose** | ARRAY | No longer used. |
| **ynose** | ARRAY | No longer used. |
| **direction** | ARRAY | No longer used. |
| **truextail** | ARRAY | An array of strings which represents the x coordinates of the rat's tail |
| **trueytail** | ARRAY | An array of strings which represents the y coordinates of the rat's tail |
| **truexnose** | ARRAY | An array of strings which represents the x coordinates of the rat's nose. |
| **trueynose** | ARRAY | An array of strings which represents the y coordinates of the rat's nose. |
| **truedirection** | ARRAY | An array which represents the current direction of the rat. |
| **xtail** | ARRAY | No longer used. |

##### *Helpful Queries for live_table*

1. The following query returns an entire column of data:

```
SELECT column_name from live_table;
```

2. The following query returns data relating to a specific session:

```
SELECT * from live_table WHERE date = 'DATE YOU DESIRE'AND subjectid
='NAME OF SPECIFIC RAT';
```

3. The following query returns the number of rat sessions in the database:

```
SELECT count(distinct(subjectid,date)) FROM live_table where notes is
null and (health='N/A';
```

4. The following query returns all of sessions completed by rats in control or baseline condition:

```
SELECT DISTINCT date,subjectid from live_table where notes is null and
(health='N/A' OR health = 'NA' OR health = 'No injection' OR health =
```

```
'(Control)' OR health = 'No Injection (Control)') ORDER BY DATE;
```

5. The following query returns all food deprived rat sessions in the database:

```
SELECT DISTINCT date,subjectid FROM live_table where LOWER(health)
LIKE '%food%dep%';
```

6. The following query returns the subjectid, referencetime, feeder, approachavoid, rewardconcentration1,rewardconcentration2, rewardconcentration3, rewardconcentration4, trialcontrolsettings for a specific session.

```
SELECT subjectid, referencetime, feeder,
approachavoid,rewardconcentration1,rewardconcentration2,rewardconcentr
ation3,rewardconcentration4,trialcontrolsettings FROM live_table WHERE
referencetime LIKE 'INSERT DESIRED SESSION DATE HERE%' AND
subjectid='INSERT NAME OF DESIRED RAT HERE'
```

#### Human Tables

The application assumes a PostegreSQL database has been created and linked (see **Decision-Making App User Guide** above**)** prior to running any session. If an appropriate data table has not been created to store session data, the application can create one automatically from the session variables if the "auto_create_table" option in the settings file (*bin/settings.ini*) is set to 1, *and* no table is detected with the name specified by the "data_table" parameter in the database linked under the "postgresql" section of the settings file. If the "data_upload" option is enabled in settings, uploads to the database will happen at three different points during a session: upon submission of the preference rankings form (see section "**Preference ranking**"), after submission of each trial question (see section "T**rial questions**"), and at the end of a session (see section "**End of session**"). If preferences are re-ranked (see section "**Preference re-ranking"**), the updated preferences will be uploaded upon form submission as well.

##### *live_database*

1. Each row of the live_database represents a trial.
2. All raw data is stored in the live_database.
3. Sessions are composed of multiple trials and can be identified by unique date and id.

Each database row is populated with the following parameters:

| Column name (varible) | data_type | Description |
|---|---|---|
| **Session_notes** | character varying | Notes made by an experimenter at the end of each session |
| **Num_stories** | character varying | The number of stories displayed in a session, set by the user in the first screen (see section "**Session startup**"), as a number between 1 and 4. |

| | | |
|---|---|---|
| pref_stories | character varying | Preferred story topics selected by the user at the Session setup window (see section "**Session setup**"). |
| Story_order | character varying | The set of stories selected to be presented to the subject based on their story preferences (see section "**Session setup**"). |
| hunger | Int | Perceived level of hunger, on a scale from 0 - 100. |
| tired | Int | Perceived level of tiredness, on a scale from 0 - 100. |
| pain | int | Perceived level of pain, on a scale from 0 - 100. |
| stress | int | Perceived level of patresssin, on a scale from 0 - 100. |
| sex | string | The subject's sex. |
| genderid | string | The subject's gender identity. |
| menstruation | string | Vague timeframe since the end of the subject's menstrual cycle, if applicable. |
| age | string | The subject's age, as a rage. |
| weight | string | The subject's weight, as a range. |
| race | string | Subject demographics. |
| relationship_status | string | The subject's romantic relationship status. |
| sexual_orientation | string | The subject's sexual orientation |
| education | string | Academic classification, if applicable |
| major | string | The academic major the subject is currently studying, if applicable |
| college | string | The college department that the subject's major belongs to, if applicable. |
| exercise | int | Rough estimation of exercise frequency |
| exercise_time_min | Int | Estimation of how long the subject's workouts tend to be on the lower range. |
| exercise_time_max | int | Estimation of how long the subject's workouts tend to be on the upper range. |
| caffeine | int | Frequency of consumption of caffeine products. |
| nicotine | int | Frequency of consumption of nicotine products. |
| alcohol | int | Frequency of consumption of alcohol |
| Vis_media | array | Categories of visual media typically consumed by the subject |
| hobbies | array | Categories of hobbies that the subject practices. |
| Next_story_index | int | The queue position of the next story. |

| | | |
|---|---|---|
| Eye_tracker_data | Json array | Data produced by the eye tracker. |
| Gaze_data | Json array | Where the subject was looking on the screen |
| eye_openness_data | array | How open the subject's eyes are during tracking, in millimeters. |
| user_position_data | aray | The user's position in the eye tracker's tracking box. |
| heart_rate_data | array | An array of heart rate values for each trial, in beats per minute (bpm). |
| tasktypedone | string | The type of task conducted, either approach-avoid, benefit-benefit, cost-cost, moral, multi-choice, probability, or social. |
| reward_prefs | array | The recorded preferences for each reward item displayed in the preference ranking window (see section **"Preference ranking"**). Each item is referenced by index |
| cost_prefs | array | The recorded preferences for each cost item displayed in the preference ranking window (see section **"Preference ranking"**). Each item is referenced by index |
| Cost_level | int | The cost intensity of the trial question presented (see section **"Trial questions").** |
| Reward_level | int | The reward intensity of the trial question presented (see section **"Trial questions"**). |
| Decision_made | int | The subject's recorded answer to a trial question, on a scale from 0 to 100. |
| trial_index | int | The question number, from 0 to one minus the value set by the "questions_per_story" parameter in settings. |
| trial_start | Timestamp string | The timestamp recorded at the start of a trial question. |
| trial_end | Timestamp string | The timestamp recorded when the subject submits their answer. |
| trial_elapsed | String | The time between the start of a trial question and answer submission. |
| Story_prefs | Array | Story preference ranking. |
| subjectidnumber | String | Subject identifier. |

| | | |
|---|---|---|
| Relationship_level | String | (For social-type stories only) The word injected into the presented story. Relationship words are selected from the word bank defined by the "relation_levels" parameter in settings. |

##### *How to Connect to database via MATLAB's "Database Explorer" App*

1. Ensure you have already downloaded MATLAB and set up your database on some machine which you can connect to through the internet.
2. Once this is done you can open MATLAB, and navigate to the "Apps" tab.
3. If you do not already have the "Database Explorer" app downloaded you can click on the "Get More Apps" button and search for "Database Toolbox."
4. Download it and it will install database explorer.
5. Open the Database Explorer App.
6. In the toolbar there is a section called "Data Source", find it (should be on the left side.
7. Click on the dropdown for the "Configure Data Source" button, it will open a drop down menu.
8. On that dropdown menu click on "Configure JDBC datasource", this will open a new window.
9. The new window will be titled "JDBC Data Source Configuration".
10. This new window will have several boxes for you to fill in.
11. The first box is labeled "name", and you should fill it in with "live_database". CAPITILIZATION IS IMPORTANT.
12. The second box is labeled "Vendor", select "PostgreSQL".
13. The third box is labeled "Driver Location", download the file from the following link:
14. https://github.com/lddavila/UTEP-Brain-Computation-Lab-Remote-Databases-and-Serendipity-App/blob/main/App%20Deployment%20Folder/postgresql-42.3.1.jar
15. Click on the "…" located next to the "Driver Location" box and select the file you just downloaded.
16. The next section is labeled "Connection Parameters", and has 3 boxes.
17. The first box is labeled "Database", inside of it put "live_database". CAPITILIZATION IS IMPORTANT.
18. The second box is labeled "Sever". In this box type the IP address of your machine. By default it contains the value localhost, but this will not work even if the computer you are working on is the one hosting the database, you must type in the IP address of the machine.
19. The third box is labeled "Port Number". The default port number is 5432, and it should still be this unless you changed it when you set up PostgreSQL on your local machine.
20. Now Look for the button labeled "test".
21. For username type in "postgres".
22. For password type in "1234".
23. Click on test.

24. If you get a success message then you have successfully connected, if not you made a mistake somewhere. Please review the steps.
25. Click on Save.

##### *Helpful MATLAB codes*

1. The following code will enable you to connect to the live_database, provided you have successfully completed the following section.

```
datasource = 'live_database'; %ENTER YOUR DATASOURCE NAME HERE,
default should be "live_database"

username = 'postgres'; %ENTER YOUR USERNAME HERE, default should be
"postgres"

password = '1234'; %ENTER YOUR PASSWORD HERE, default should be "1234"
```

2. The following code shows a general syntax for writing a query in MATLAB, which can be submitted to the database. (Reference the “Helpful Queries for live_table” section for ideas on queries you might want)

```
query = “YOUR QUERY HERE;”;
```

3. The following code will execute the query, and store the result of your query into a variable that MATLAB can work for. (remember that everything will be returned to MATLAB as a table, so you should handle it accordingly.

```
conn = database(datasource,username,password);

result = fetch(conn,query)
```

4. The following code will execute a query which doesn’t return anything.

```
execute(conn,query);
```

### Figure Recreation Guide

#### Code Repositories

*Figures 1G, 1J, 3C, 3D-G, 5C-P and corresponding supplements can be found in the following GitHub repository:* https://github.com/lddavila/human_dec_making

##### Setup Part 1

1. To access code relevant to the above figures, run the following command:

```
git clone https://github.com/lddavila/human_dec_making.git
```

2. Open the file "session_cost_clustering_figures_threshold_0.m", linked here: https://github.com/lddavila/human_dec_making/blob/main/session_cost_clustering_figures_threshold_0.m
3. Run the section titled "SETUP PART 1: add the Utility functions to my path", lines 1-9.
4. Run the section titled "SETUP PART 5: put rat and human data into a table to be used", lines 25-32.
5. Run the section titled "SETUP PART 6: get human and rat data information (i.e. # of subjects, # of data points)", lines 36-38.
6. This loads all the data required to recreate the figures listed above.

*Figures 1H, 1J, 2A-K, 3A, 3B, 4A-F, 5A, 5B, 6A-E and corresponding supplements can be found in the following GitHub repository:*https://github.com/lrakocev/human_dm/tree/permutations .

##### Set up Part 2

1. To access code relevant to the above figures, run the following command:

```
git clone https://github.com/lrakocev/human_dm.git
```

2. Open the file "hum_new_tasks_runme.m", linked here: https://github.com/lrakocev/human_dm/blob/permutations/hum_new_tasks_runme.m
3. Change the variables datasource (line 3), username (line 4), password (line 5) to the variables which you assigned when setting up your database connection in the MATLAB "Database Explorer" app.
4. Run the section titled "Human New Tasks Run Me", located in lines 1-36.
5. If instead you would like to use the pre-loaded data (which was attained by following the process above), download the Matlab files from: https://github.com/lrakocev/human_dm/blob/permutations/ingest%20helpers/human%20data.mat

#### Data Repositories

https://doi.org/10.7910/DVN/OZARPL

We ultimately collected data from 35 subjects across all four tasks, resulting in 982 sessions in total. Our primary fitting process found 71% of this data to be sigmoidal, yielding 693 sessions used in total for analyses that are dependent on the usage of psychometric functions. Differences in this number may arise depending on when the analysis was performed, as we continued to collect more data throughout the writing of this manuscript. For some analyses, we make use of an additional level of granularity by splitting sessions by cost level as well. For these analyses, we start with 3928 initial points, to which we apply the same fitting process. We again use only sigmoidal session-cost level curves, yielding 2500 points in total. All these analyses are based on the data in the database linked above.

#### Figure 1G

1. Open "session_cost_clustering_figures_threshold_0.m", linked here: https://github.com/lddavila/human_dec_making/blob/main/session_cost_clustering_figures_threshold_0.m
2. Run the section titled "SETUP PART 1: add the Utility functions to my path", lines 1-9.
3. Run the section titled "SET UP PART 5: put rat and human data into a table to be used", located from lines 32-25.
4. Run the section titled "SET UP PART 6: get human and rat data information (i.e. # of subjects, # of data points)", located from lines 36-38.
5. Run the section titled "Fig 1G: average decision making maps across tasks", located in lines 39-40

#### Statistics for Fig. 1G

To test the significance of splitting by reward and cost, we ran a 3-way ANOVA test (anova, MATLAB, https://www.mathworks.com/help/stats/anova.html#mw_54c33315-7e4d-48cf-8174-8db71d907536 ). This testing returned a significance level of p=6.4718e-54 for reward and p=5.5797e-57 for cost.

#### Figure 1H

To re-create the psychometric function fitting process from the initial behavioral data:

1. Open "cluster_runme.m", linked here: https://github.com/lrakocev/human_dm/blob/permutations/clustering/fit%20psychometric%20functions/cluster_runme.m
2. In line 3, set "new_dir" to the name of the directory to save fit functions to.
3. Run the section labeled "find session sigmoids", lines 29-40.
4. Once finished, run the section titled "get % sigmoidal vs non sigmoidal", lines 42-49.

To start with the already performed psychometric function fitting:

1. Open "cluster_runme.m", linked here:

https://github.com/lrakocev/human_dm/blob/permutations/clustering/fit%20psychometric%20functions/cluster_runme.m

2. Change the location in line 44 to point to the following table: https://github.com/lrakocev/human_dm/tree/permutations/test_run/session_clustering
3. Change the “save_to” variable, located on line 46, to point towards where you would like the results saved: https://github.com/lrakocev/human_dm/tree/permutations/test_run/psych_cost_session_counts
4. Run the section titled “get % sigmoidal vs non sigmoidal”, located in lines 42-49.

#### Figure 1J

For the main plot showing clusters:

1. Open “session_clustering_figures.m”, linked here: https://github.com/lddavila/human_dec_making/blob/main/session_clustering_figures.m
2. Run the section titled “SETUP PART 1: add the Utility functions to my path”, located from lines 1-9.
3. Run the section labeled “Figure 1J: Used just to get plot, don't refer to this”.
4. Run the section titled “SET UP PART 5: put rat and human data into a table to be used”, located in lines 32-35.

For the cluster-specific subplots:

5. Open “per_cluster_runme.m” linked here: https://github.com/lrakocev/human_dm/blob/permutations/clustering/per%20cluster%20analysis/per_cluster_runme.m
6. Change the location in line 3 to point to the following table: https://github.com/lrakocev/human_dm/blob/permutations/all_session_updated.xlsx
7. Run the runme.

#### Statistics for Fig. 1J

We fit 982 sessions with sigmoidal models (fit, MATLAB, https://www.mathworks.com/help/curvefit/fit.html ) and 693 of these sessions returned a sigmoid with an r-squared of at least 0.4. The fitting parameters were scaled, (log, MATLAB), (abs, MATLAB). These scaled parameters clustered into 5 groups, identified using Fuzzy C-Means (fcm, MATLAB, https://www.mathworks.com/help/fuzzy/fcm.html ). To measure the validity of this clustering we calculated the Modified Partition Coefficient (Friedman, https://github.com/lddavila/human_dec_making/blob/main/Utility%20Functions/calculate_mpc.m ), returning a score of 0.63697.

#### Extended Data Fig. 1

1. Open the file “hum_new_tasks_runme.m”, linked here: https://github.com/lrakocev/human_dm/blob/permutations/hum_new_tasks_runme.m
2. Load the following data: https://github.com/lrakocev/human_dm/blob/permutations/ingest%20helpers/human%20data.mat
    a. This was created at an earlier date by running lines 1-36 of the “hum_new_tasks_runme.m” file.
3. Run the section titled “get counts,” located in lines 44-66.
4. Update the “path_to_save” variable on line 106 and 135 to point to a directory on your local computer.
5. Run the section titled “normalization bar plots” in lines 71-81 for Extended Data Figs. 1B-I.
6. Run the section titled “plotting summary stats”, located in lines 125-158.
    a. Line 150 yields: Extended Data Figs. 1J, 1K, 1O, 1P, 1T, 1U, 1Y, 1Z
    b. Line 153 yields: Extended Data Figs. 1M, 1N, 1R, 1P, 1W, 1X, 1AB, 1AC
7. Run the section titled “dec making maps”, located in lines 96-108 for example maps such as in Extended Data Figs. 1L, 1Q, 1V, 1AA.

#### Extended Data Fig. 2

1. Open the file “hum_new_tasks_runme.m”, linked here: https://github.com/lrakocev/human_dm/blob/permutations/hum_new_tasks_runme.m
2. Load the following data: https://github.com/lrakocev/human_dm/blob/permutations/ingest%20helpers/human%20data.mat
    a. This was created at an earlier date by running lines 1-36 of the “hum_new_tasks_runme.m” file.
3. Run the section titled “get counts,” located in lines 44-66.
4. Update the “path_to_save” variable on line 106 and 135 to point to a directory on your local computer.
5. Run the section titled “plotting summary stats”, located in lines 125-158.
    b. Line 146 yields: Extended Data Figs. 2A, 2B, 2G, 2H, 2M, 2N, 2S, 2T
    c. Line 156 yields: Extended Data Figs. 2C, 2D, 2I, 2J, 2O, 2P, 2U, 2V.
8. Run the section titled “dec making maps”, located in lines 96-108 for example maps such as in Extended Data Figs. 2E, 2F, 2K, 2L, 2Q, 2R, 2W, 2X.

#### Extended Data Figure 3A

1. Open “alt_clustering_runme.m”, linked here: https://github.com/lrakocev/human_dm/blob/permutations/clustering/alternative%20clustering/alt_clustering_runme.m
2. Change the location in line 5 to point to the following table:

    https://github.com/lrakocev/human_dm/blob/permutations/ingest%20helpers/human%20

data.mat

3. Change the “file_name” variable in line 26 to what you would like to name your new file.
4. Change the “type” and “save_to” variables in line 42-43 to point to the following table: https://github.com/lrakocev/human_dm/blob/permutations/all_session_updated.xlsx
5. Change the “table_name” variable in line 54 to point to the “file_name” set in step 3.
6. Run the rest of the runme. The section titled “clustering 2d sigs” will create the plots of clustered points, and the section titled “dec making plot per “cluster”” will

#### Statistics for Extended Fig. 3A

We fit 982 sessions to a 2D sigmoidal model (fit, MATLAB, https://www.mathworks.com/help/curvefit/fit.html ) and took the parameters of the functions as coordinates. The parameters clustered into 6 groups, identified using Fuzzy C-Means (fcm, MATLAB, https://www.mathworks.com/help/fuzzy/fcm.html ). To measure the validity of this clustering we calculated the Modified Partition Coefficient (Friedman, https://github.com/lddavila/human_dec_making/blob/main/Utility%20Functions/calculate_mpc.m ), returning a score of 0.73084.

### Figure 2A

To re-create all of the features from scratch that will be used, follow these directions first. This will take several hours to run:

1. Run the following git clone command:

   git clone https://github.com/lrakocev/dec-making-app.git

2. Run “eye_tracker_feats.sh", located here: https://github.com/lrakocev/dec-making-app/blob/lr_analysis/eye_tracking_feats.sh
   a. This will also yield examples like in Extended Data Figs. 3C, 3D.
3. Run “eye_tracker_helper.py”, located here: https://github.com/lrakocev/dec-making-app/blob/lr_analysis/eye_tracker_helper.py
4. Run “hr_feats.sh”, located here: https://github.com/lrakocev/dec-making-app/blob/lr_analysis/hr_feats.sh
   a. This will also yield examples like in Extended Data Figs. 3E-J.
5. Run “hr_tracker_helper.py”, located here: https://github.com/lrakocev/dec-making-app/blob/lr_analysis/hr_tracker_helper.py

To use the already-created features, continue with:

6. Open “run_corr_to_clusters.m”, located here: https://github.com/lrakocev/human_dm/blob/permutations/physio/run_corr_to_clusters.m
7. Download the eye-tracking and heart-rate feature data from the following link: https://doi.org/10.7910/DVN/OZARPL.

8. Change locations in lines 2-6, 11-14 to point to data downloaded in step 7.
9. Change the location in line 19 to point to the following table: https://github.com/lrakocev/human_dm/blob/permutations/all_session_updated.xlsx
10. Change the location in line 24 to point to the following table: https://github.com/lrakocev/human_dm/blob/permutations/ingest%20helpers/human%20data.mat
11. Run sections “get psych data”, “merging tables”, and “3d plots for clusters”. Line 69 produces Fig. 2A.

#### Statistics for Fig. 2A

We used 6938 trials in total for these tests both in this figure and in Extended Data Figs. Data was collected on a trial-by-trial basis, so for each session there are theoretically 16 recordings. However, not every recording was successful or could be measured. We applied no additional thresholding beyond what the equipment was able to record. To test the significance of differences in physiological features across clusters, we ran a 3-way ANOVA test (anova, MATLAB, https://www.mathworks.com/help/stats/anova.html#mw_54c33315-7e4d-48cf-8174-8db71d907536 ). This testing returned a significance level of p=0.012 for differences in features across clusters.

### Figure 2B, 2C

1. Open “prims_run_me.m”, linked here: https://github.com/lrakocev/human_dm/blob/permutations/primitive%20building/prims_run_me.m
2. Change “hum_table_name” in line 4 to point to the following table: https://github.com/lrakocev/human_dm/blob/permutations/clustering/create%20clusters/og_cluster_dir_10-28-2024/all%20human%20data.xlsx
3. Change the location in line 9 to point to the following table:
4. https://github.com/lrakocev/human_dm/blob/permutations/ingest%20helpers/human%20data.mat
5. Run the sections titled “concatenating the session tables into one big table per task”. This section produces Fig. 2B, 2C.

### Figure 2D, 2E

1. Run the steps 1-4 for Fig. 2B,2C.
2. Open “example_prims.m”, linked here; https://github.com/lrakocev/human_dm/blob/permutations/primitive%20building/example_prims.m
3. Run the section “individual exs”.
    a. For 2C, change the following variables: “id” in line 18 to 88407, “task” in line 19 to “moral”, and “story” in line 20 to XX.

b. For 2D, change the following variables: “id” in line 18 to 21188, “task” in line 19 to “probability”, and “story” in line 20 to XX.

#### Figure 2F, G

1. Once steps 1-4 of the process for creating Fig 2B,2C have been run, continue within the same file: https://github.com/lrakocev/human_dm/blob/permutations/primitive%20building/prims_run_me.m
2. Run the section titled “subject variance plots”

#### Figure 2H, I

1. Run the steps 1-4 for Fig. 2B, 2C.
2. Open “example_prims.m”, found here; https://github.com/lrakocev/human_dm/blob/permutations/primitive%20building/example_prims.m
3. Run the section “individual exs”.
   c. For 2I, change the following variables: “id” in line 18 to 80925, “task” in line 19 to “probability”, and “story” in line 20 to XX.
   d. For 2J, change the following variables: “id” in line 18 to 98865, “task” in line 19 to “social”, and “story” in line 20 to XX.

#### Figure 2J, Extended Data Figs. 3B, 3C

1. Open “prims_run_me.m”, linked here: https://github.com/lrakocev/human_dm/blob/permutations/primitive%20building/prims_run_me.m
2. Change “hum_table_name” in line 4 to point to the following table: https://github.com/lrakocev/human_dm/blob/permutations/clustering/create%20clusters/og_cluster_dir_10-28-2024/all%20human%20data.xlsx
3. Change the location in line 9 to point to the following table: https://github.com/lrakocev/human_dm/blob/permutations/ingest%20helpers/human%20data.mat
4. Run the sections titled “concatenating the session tables into one big table per task”
5. Run the section titled “prim histogram”. In this figure, we show:
   a. Variance within subject – line 37
   b. Reward interaction – line 40
   c. Reward impulsivity – line 42
   d. Mean approach rate – line 44
6. In the Extended Data Figs. 3A, 3B, we also show:
   a. Raw elasticity – line 48.
   b. Variance from cluster mean – line 39.

#### Statistics for Fig. 2J

To test the significance of differences in mean features across clusters, we ran a one-way ANOVA test (anova, MATLAB, https://www.mathworks.com/help/stats/anova.html#mw_54c33315-7e4d-48cf-8174-8db71d907536 ). This testing returned a significance level of $p=9.66\times10^{-11}$ for differences in reward interaction, p=0.036 for differences in reward impulsivity, and $p=3.06\times10^{-8}$ for differences in mean approach rates. To test the significance in differences in variance features across clusters, we used pairs of F-tests (f-test, MATLAB, https://www.mathworks.com/help/stats/vartest2.html ). We then chose the maximum of all pairs significance to report. This testing returned a significance level of $p<1\times10^{-68}$ for differences in within-subject variance.

#### Extended Data Figs. 3D, 3E, 3F-K, 3L, 3M, 3N, 3O

To re-create all of the features from scratch that will be used, follow these directions first. This will take several hours to run:

12. Run the following git clone command:

    git clone https://github.com/lrakocev/dec-making-app.git

13. Run "eye_tracker_feats.sh", located here: https://github.com/lrakocev/dec-making-app/blob/lr_analysis/eye_tracking_feats.sh
    a. This will also yield examples like in Extended Data Figs. 3C, 3D.
14. Run "eye_tracker_helper.py", located here: https://github.com/lrakocev/dec-making-app/blob/lr_analysis/eye_tracker_helper.py
15. Run "hr_feats.sh", located here: https://github.com/lrakocev/dec-making-app/blob/lr_analysis/hr_feats.sh
    a. This will also yield examples like in Extended Data Figs. 3E-J.
16. Run "hr_tracker_helper.py", located here: https://github.com/lrakocev/dec-making-app/blob/lr_analysis/hr_tracker_helper.py

To use the already-created features, continue with:

17. Open "run_corr_to_clusters.m", located here: https://github.com/lrakocev/human_dm/blob/permutations/physio/run_corr_to_clusters.m
18. Download the eye-tracking and heart-rate feature data from the following link: https://doi.org/10.7910/DVN/OZARPL.
19. Change locations in lines 2-6, 11-14 to point to data downloaded in step 7.
20. Change the location in line 19 to point to the following table: https://github.com/lrakocev/human_dm/blob/permutations/all_session_updated.xlsx
21. Change the location in line 24 to point to the following table: https://github.com/lrakocev/human_dm/blob/permutations/ingest%20helpers/human%20data.mat
22. Run sections "get psych data", "merging tables", and "3d plots for clusters". Line 69

produces Fig. 2K. Line 67 and 68 produce Extended Data Figs. 3K and 3L respectively.

23. Run section “3d plot for tasks” for Extended Data Figs. 3M and 3N.

#### Statistics for Extended Data Figs. 3L, 3M, 3N, 3O

To test the significance of differences in heart-rate features only across clusters, we ran a one-way ANOVA (anova, MATLAB, https://www.mathworks.com/help/stats/anova.html#mw_54c33315-7e4d-48cf-8174-8db71d907536 ) based on the feature of relative maximum heart rate. This yielded a significance level of p=0.028773. To test the significance of differences in eye-tracking features only across clusters, we ran a one-way ANOVA based on the feature average pupil diameter. This returned a significance level of p=0.00095. We also examined task-based differences and found that we could separate tasks based on eye-tracking and heart-rate features as well, using ANOVA. We performed a one-way ANOVA using relative maximum heart rate across tasks and found a significant differences with p=4.1839e-11. We performed a three-way ANOVA using average pupil diameter, number of local minima in pupil diameters over the trial duration, and number of local maxima in pupil diameter over the trial duration, and found significant differences across tasks with a significance level of p=1.1987e-67.

### Figure 3A

1. Open “hum_new_tasks_runme.m”, linked here: https://github.com/lrakocev/human_dm/blob/permutations/hum_new_tasks_runme.m
2. Run section Load the following data: https://github.com/lrakocev/human_dm/blob/permutations/ingest%20helpers/human%20data.mat
3. Run the section titled “overlapped for fig 6”, on lines 160-214. Change “path_to_save” to a local path to save the files to.

#### Statistics for Fig. 3A

To test the significance of differences in approach rates at varying reward and cost levels across tasks, we ran a 2-way ANOVA test (anova, MATLAB, https://www.mathworks.com/help/stats/anova.html#mw_54c33315-7e4d-48cf-8174-8db71d907536 ). This testing returned a significance level of $p= 3x10^{-6}$ for differences in cost levels across clusters and p= 0.002 for differences in reward levels across clusters.

### Figure 3B

1. Change the location in line 3 to point to the following table: https://github.com/lrakocev/human_dm/blob/permutations/clustering/create%20clusters/og_cluster_dir_10-28-2024/all%20human%20data.xlsx
2. Open the file “task_diffs_runme.m”, linked here: https://github.com/lrakocev/human_dm/blob/permutations/primitive%20building/task_dif

fs_runme.m

3. Run the section labeled “cluster proportions” on lines 20-25.

#### Figure 3C

To get the main figure of clustered data, follow these directions:

1. Open “session_cost_clustering_figures_threshold_0.m”, linked here: https://github.com/lddavila/human_dec_making/blob/main/session_cost_clustering_figures_threshold_0.m
2. Run the section titled “SETUP PART 1: add the Utility functions to my path”, located from lines 1-9.
3. Run the section titled “SET UP PART 5: put rat and human data into a table to be used”, located from lines 32-25.
4. Run the section titled “SET UP PART 6: get human and rat data information (i.e. # of subjects, # of data points)”, located from lines 36-38.
5. Run the section titled “FIG 1J: DISCRETE DECISION-Making STRATEGIES”, located in lines 43-65.
6. Lines 44-50 will create the 3D cluster plot.

To get the average sigmoid per cluster per task:

7. Using the same data and setup as above.
8. Open “per_cluster_runme.m”, located here: https://github.com/lrakocev/human_dm/blob/permutations/clustering/per%20cluster%20analysis/per_cluster_runme.m
9. Run line 37, setting “spectral_table” to the data from step 3 above.

#### Statistics for Fig. 3C

To find human clusters we split 982 sessions by reward and cost into 3830 sessions across 35 subjects. We fit these sessions with sigmoidal models (fit, MATLAB, https://www.mathworks.com/help/curvefit/fit.html ). 2500 sessions returned an r-squared value of at least 0.4. The fitting parameters were scaled, (log, MATLAB), (abs, MATLAB). These scaled parameters clustered into 5 groups, identified using Fuzzy C-Means (fcm, MATLAB, https://www.mathworks.com/help/fuzzy/fcm.html ). To measure the validity of this clustering we calculated the Modified Partition Coefficient (Friedman, https://github.com/lddavila/human_dec_making/blob/main/Utility%20Functions/calculate_mpc.m ), returning a score of 0.58335.  We then averaged and fit (fit, MATLAB) the original data of each cluster with a sigmoidal model which returned an r-squared rating of at least 0.4 to get the average sigmoid.

#### Extended Data Figures 4A

1. Open “incremental_sessions_3d_plot.m”, linked here: https://github.com/lddavila/human_dec_making/blob/main/incremental_sessions_3d_plot.m
2. Run the section titled “Set up part 1: add the Utility functions to my path”, located in lines 1-9. (Ensure that the data_dir variable is updated to point to the absolute path of the “session clustering” directory which you should have already cloned from GitHub.
3. Run the section titled “Set up part 2: copy the data (if necessary) and make plot” located from lines 10-49.

#### Statistics for Extended Data Fig. 4A

We start with 693 sessions. For each session/story, we asked participants to rate the relevance of the situation to themselves. We choose a threshold of 70 and apply it to the data, yielding 243 sessions in total across 31 subjects.   We then use Fuzzy C-Means (fcm, MATLAB, https://www.mathworks.com/help/fuzzy/fcm.html ) to identify the clusters within this set. To measure the validity of this clustering we calculated the Modified Partition Coefficient (Friedman, https://github.com/lddavila/human_dec_making/blob/main/Utility%20Functions/calculate_mpc.m ), returning a score of 0.70857.

### Extended Data Figures 4B

1. Open the file “create_threshold_3d_plots.m”, linked here: https://github.com/lddavila/human_dec_making/blob/main/create_threshold_3d_plots.m
2. Run the section titled “Supplemental Figure 3D”, located in lines 7-13.

#### Statistics for Extended Data Fig. 4B

We start with 693 sessions. For each session/story, we asked participants to rate the relevance of the situation to themselves. We choose a threshold of 70 and apply it to the data, yielding 243 sessions in total across 31 subjects. Once these sessions have been identified, we increase the level of granularity by looking at cost level sigmoids, so that each session is split into 4 cost levels as well. We then use Fuzzy C-Means (fcm, MATLAB, https://www.mathworks.com/help/fuzzy/fcm.html ) to identify the clusters within this set. To measure the validity of this clustering we calculated the Modified Partition Coefficient (Friedman, https://github.com/lddavila/human_dec_making/blob/main/Utility%20Functions/calculate_mpc.m ), returning a score of 0.58366.

### Extended Data Figures 4C

1. Open the file “session_cost_clustering_figures_threshold_0.m”

2. Run the section titled “SETUP PART 1: add the Utility functions to my path”, located in lines 1-9
3. Run the section titled “SET UP PART 5: put rat and human data into a table to be used”, located in lines 32-35
4. Run the section titled “SET UP PART 6: get human and rat data information (i.e. # of subjects, # of data points)”, located in lines 36-38
5. Run the section titled “Supplemental Figure 3E: cluster locations significantly differ”, located from lines 274-278.

#### Statistics for Extended Data Fig. 4C

To determine if there was significant difference between the human clusters, we measured the Bhattacharyya distance (bhattacharyyaDistance, MATLAB, https://www.mathworks.com/help/predmaint/ref/bhattacharyyadistance.html ) between each dimension of the clusters, which represent the fitting parameters of the sigmoidal models. Cluster 1 had 388 members, cluster 2 had 156 members, cluster 3 had 1550 members, cluster 4 had 95 members, and cluster 5 had 311 members.

#### Extended Data Figures 4D, 4E

1. Open the file “session_cost_clustering_figures_threshold_0.m”
2. Run the section titled “SETUP PART 1: add the Utility functions to my path”, located in lines 1-9
3. Run the section titled “SET UP PART 5: put rat and human data into a table to be used”, located in lines 32-35
4. Run the section titled “SET UP PART 6: get human and rat data information (i.e. # of subjects, # of data points)”, located in lines 36-38
5. Run the section titled “Supplemental Figure 3F: means”, located from lines 279-282. This produces Extended Data Fig. 4D.

#### Statistics for Extended Data Fig. 4D, 4F

To determine if there was significant difference between the human clusters’ means, we performed a 2 sample t-test (ttest2, MATLAB, https://www.mathworks.com/help/stats/ttest2.html ) between each dimension of each cluster. To determine if there was significant difference between the human clusters’ variances, we performed a 2 sample f-test (vartest2, MATLAB, https://www.mathworks.com/help/stats/vartest2.html) between each dimension of each cluster. Cluster 1 had 388 members, cluster 2 had 156 members, cluster 3 had 1550 members, cluster 4 had 95 members, and cluster 5 had 311 members.

#### Extended Data Figures 4F-K

1. Open the file “session_cost_clustering_figures_threshold_0.m”, linked here: https://github.com/lddavila/human_dec_making/blob/main/session_cost_clustering_figures_threshold_0.m
2. Run the section titled “SETUP PART 1: add the Utility functions to my path”, located in lines 1-9
3. Run the section titled “SET UP PART 5: put rat and human data into a table to be used”, located in lines 32-35
4. Run the section titled “SET UP PART 6: get human and rat data information (i.e. # of subjects, # of data points)”, located in lines 36-38
5. Run the section titled “Supplemental Figure 4A-4F: Task Comparisons” located in lines 141-143.

#### Statistics for Extended Data Figs. 4F-K

To determine if there was significant difference in cluster counts between tasks we split the clustered 2500 sessions across 35 subjects by their tasks resulting in 683 sessions across 25 subjects for approach avoid, 577 sessions across 29 subjects for moral, 623 sessions across 29 subjects for probability, and 617 sessions across 27 subjects for social. We then found each tasks’ cluster counts, and compared them to each other using a chi squared test (chi2test, MathWorks File Exchange, https://www.mathworks.com/matlabcentral/fileexchange/16177-chi2test ). The results of each comparison can be seen in Extended Data Figs 4F-K.

### Figure 3D

1. Open “session_cost_clustering_figures_threshold_0.m”, linked here: https://github.com/lddavila/human_dec_making/blob/main/session_cost_clustering_figures_threshold_0.m
2. Run the section titled “SETUP PART 1: add the Utility functions to my path”, located from lines 1-9.
3. Run the section titled “SET UP PART 5: put rat and human data into a table to be used”, located in lines 32-35.
4. Run the section titled “SET UP PART 6: get human and rat data information (i.e. # of subjects, # of data points)”, located in lines 36-38.
5. Run the section titled “FIG 1K: Strategy Utilization Differs across tasks”, located in lines 66-67.

#### Statistics for Fig. 3D

To define statistical difference between proportions of clusters in different tasks we used 2500 sessions across 4 cost levels and 4 reward levels: approach avoid contained 683 sessions across 25 subjects, moral had 577 sessions across 29 subjects, probability with 623 sessions across 29 subjects, and social with 617 sessions across 27 subjects. We used the chi squared test (chi2test, MathWorks File Exchange https://www.mathworks.com/matlabcentral/fileexchange/16177-chi2test ), inputting the number of points of each cluster for each task, and comparing 2 tasks at a time. The resulting levels of significance from these comparisons can be seen in figure 3D.

#### Figure 3E, 3F

1. Open "session_cost_clustering_figures_threshold_100.m", linked here: https://github.com/lddavila/human_dec_making/blob/main/session_cost_clustering_figures_threshold_100.m
2. Run the section titled "SETUP PART 1: add the Utility functions to my path", located in lines 1-9.
3. Run the section titled "SET UP PART 4: put rat and human data into a table to be used", located in lines 23-26.
4. Run the section titled "SET UP PART 5: get human data information (ie. # of subjects, # of data points )", located in lines 27-29.
5. Run the section titled "FIG 1L: create the average xyz plot", located in lines 30-33.
6. Run the section titled "FIG 1M: try to recreate xyz using data which is not significantly different data from each task (SF3Q) version 2", located in lines 33-38.

#### Statistics for Fig. 3E

To determine if there was a statistically significant difference between the tasks' means, we filtered 2500 sessions down to the 306 sessions that subjects defined as most relevant: approach avoid had 143 sessions across 16 subjects, moral had 53 sessions across 13 subjects, probability had 62 sessions across 11 subjects, and social had 48 sessions across 13 subjects. Then we ran a 1-way anova test (anova, MATLAB, https://www.mathworks.com/help/stats/anova.html#mw_54c33315-7e4d-48cf-8174-8db71d907536 ), inputting the fitting parameters. This returned a significance level of 0.0043289.

#### Figure 3G

1. Open file titled "session_cost_clustering_figures_threshold_100.m", linked here: https://github.com/lddavila/human_dec_making/blob/main/session_cost_clustering_figures_threshold_100.m
2. Run the section titled "SETUP PART 1: add the Utility functions to my path", located in lines 1-9
3. Run the section titled "SET UP PART 4: put rat and human data into a table to be used", located in lines 23-26
4. Run the section titled "SET UP PART 5: get human and rat data information (i.e. # of subjects, # of data points)", located in lines 27-29.
5. Run the section titled "Supplemental Figure 4M: recreate average xyz plots using explicitly wrong proportions", located in lines 46-52.

#### Extended Data Fig. 4L

1. Open the file “session_cost_clustering_figures_threshold_0.m”, linked here: https://github.com/lddavila/human_dec_making/blob/main/session_cost_clustering_figures_threshold_0.m
2. Run the section titled “SETUP PART 1: add the Utility functions to my path”, located in lines 1-9
3. Run the section titled “SET UP PART 5: put rat and human data into a table to be used”, located in lines 32-35
4. Run the section titled “SET UP PART 6: get human and rat data information (i.e. # of subjects, # of data points)”, located in lines 36-38
5. Run the section titled “SF4G, SF4H, SF4I: Clusters across task have same location”, located in lines 103-115.

#### Statistics for Extended Data Fig. 4L

To determine if there was significant difference in clusters between tasks, we split the clustered 2500 sessions across 35 subjects by their tasks resulting in 683 sessions across 25 subjects for approach avoid, 577 sessions across 29 subjects for moral, 623 sessions across 29 subjects for probability, and 617 sessions across 27 subjects for social. We then measured the Bhattacharyya Distance along each dimension for each of the clusters and experiments (bhattacharyyaDistance, MATLAB, https://www.mathworks.com/help/predmaint/ref/bhattacharyyadistance.html ). The resulting Bhattacharyya distance can be seen in Extended Data Fig. 5I.

#### Figure 4A, Extended Data Fig. 5A, 5B, 5C

1. Open “subjects_in_cluster_runme.m”, linked here: https://github.com/lrakocev/human_dm/blob/permutations/clustering/subjects%20in%20cluster/subjects_in_cluster_runme.m
2. Change the location in line 3 to point to the following table: https://github.com/lrakocev/human_dm/blob/permutations/clustering/create%20clusters/og_cluster_dir_10-28-2024/all%20human%20data.xlsx
3. Change the location in line 8 to point to the following table: https://github.com/lrakocev/human_dm/blob/permutations/ingest%20helpers/human%20data.mat
4. Run sections “get psych data” on lines 10-27 and section “individual subjects strategies across clusters” on lines 51-60. We show the “approach avoid” task in the main text.

#### Figure 4B

1. Open “subjects_in_cluster_runme.m”, linked here: https://github.com/lrakocev/human_dm/blob/permutations/clustering/create%20clusters/og_cluster_dir_10-28-2024/all%20human%20data.xlsx
2. Change the location in line 3 to point to the following table:

https://github.com/lrakocev/human_dm/blob/permutations/clustering/create%20clusters/og_cluster_dir_10-28-2024/all%20human%20data.xlsx

3. Change the location in line 8 to point to the following table: https://github.com/lrakocev/human_dm/blob/permutations/ingest%20helpers/human%20data.mat
4. Run sections "get psych data" on lines 10-27 and section "individual subjects strategies across clusters" on lines 40-49.

#### Figure 4C

1. Open "subjects_in_cluster_runme.m", linked here: https://github.com/lrakocev/human_dm/blob/permutations/clustering/subjects%20in%20cluster/subjects_in_cluster_runme.m
2. Change the location in line 3 to point to the following table: https://github.com/lrakocev/human_dm/blob/permutations/clustering/create%20clusters/og_cluster_dir_10-28-2024/all%20human%20data.xlsx
3. Change the location in line 8X to point to the following table: https://github.com/lrakocev/human_dm/blob/permutations/ingest%20helpers/human%20data.mat
4. Run sections "subjects in cluster", lines 29-38. We show the figure for "all" tasks.

#### Extended Data Fig. 5D

1. Open "prob_runme.m", linked here: https://github.com/lddavila/human_dec_making/blob/main/session_cost_clustering_figures_threshold_0.m
2. Change the location in line 4 to point to the following table: https://github.com/lrakocev/human_dm/blob/permutations/all_session_updated.xlsx
3. Change the location in line 10 to point to the following table: https://github.com/lrakocev/human_dm/blob/permutations/ingest%20helpers/human%20data.mat
4. Run the section runme. We show the figure for "all" tasks.

#### Extended Data Fig. 5E

1. Open "chain_behavior_runme.m", linked here: https://github.com/lrakocev/human_dm/blob/permutations/clustering/chaining/chain_behavior_runme.m
2. Change the location in line 4  to point to the following table: https://github.com/lrakocev/human_dm/blob/permutations/all_cost_5_clusters.xlsx
3. Load behavioral data, located here: https://github.com/lrakocev/human_dm/blob/permutations/ingest%20helpers/human%20data.mat

4. Change the location in lines 76-68 to point to the following table: https://github.com/lrakocev/human_dm/blob/permutations/figs/all_cost_5_clusters/chaining/total_table_all_cost_5_clusters.mat
5. Run the section titled “validation plots”, lines 74-87. Line 87 produces Extended Data Fig. 5E.

#### Figure 4D, 4E, Extended Data Fig. 5F, 5G

6. Open “chain_behavior_runme.m”, linked here: https://github.com/lrakocev/human_dm/blob/permutations/clustering/chaining/chain_behavior_runme.m
7. Change the location in line 4 to point to the following table: https://github.com/lrakocev/human_dm/blob/permutations/all_cost_5_clusters.xlsx
8. Load behavioral data, located here: https://github.com/lrakocev/human_dm/blob/permutations/ingest%20helpers/human%20data.mat
9. Change variable”type” in line 48 to “individual”.
10. Run sections “get spectral all_psych_data” in lines 11-27 and “calcs + plotting" on lines 46-70.
11. We pick subjects 11464 and 21188 as examples in Fig. 4D and 4E respectively.

#### Figure 4F, Extended Data Fig. 5H

1. Open “chain_behavior_runme.m”, linked here: https://github.com/lrakocev/human_dm/blob/permutations/clustering/chaining/chain_behavior_runme.m
2. Change the location in line 4 to point to the following table: https://github.com/lrakocev/human_dm/blob/permutations/all_cost_5_clusters.xlsx
3. Load behavioral data, located here: https://github.com/lrakocev/human_dm/blob/permutations/ingest%20helpers/human%20data.mat
4. Run sections “get spectral all_psych_data” in lines 11-27 and “calcs + plotting" on lines 29-44.

#### Figure 5A

1. Open “fig5a_rats.m”, linked here: https://github.com/lrakocev/human_dm/blob/permutations/psychometric%20analysis/fig5a_rats.m
2. Connect to the database for rat data by editing lines 3, 4, and 5 to point to the correct location as described in the **Database Guide** above.
3. Run the script. We use the example for rat “Johnny”.

#### Figure 5B

1. Open “fig5a_humans.m”, linked here: https://github.com/lrakocev/human_dm/blob/permutations/psychometric%20analysis/fig5a_rats.m
2. Load behavioral data, located here: https://github.com/lrakocev/human_dm/blob/permutations/psychometric%20analysis/fig5a_humans.m
3. Run the script.

#### Figure 5C

1. Open “session_cost_clustering_figures_threshold_0.m”, linked here: https://github.com/lddavila/human_dec_making/blob/main/session_cost_clustering_figures_threshold_0.m
2. Run the section titled “SETUP PART 1: add the Utility functions to my path”, located from lines 1-9.
3. Run the section titled “SET UP PART 5: put rat and human data into a table to be used”, located from lines 32-25.
4. Run the section titled “SET UP PART 6: get human and rat data information (i.e. # of subjects, # of data points)”, located from lines 36-38.
5. Run the section titled “FIG 2C: RAT DISCRETE DECISION STRATEGIES”, located in lines 68-76. This will create the 3D Cluster Plot.
6. Run the section titled “Fig 2C: Average Sigmoid Per Cluster”, located in lines 76-79. This will create the average sigmoidal plots.

#### Statistics for Fig. 5C

To identify clusters we fit 2106 rat sessions, across 42 subjects with sigmoidal models (fit, MATLAB, https://www.mathworks.com/help/curvefit/fit.html ). 1616 sessions returned an r-squared value of at least 0.4. T The fitting parameters were scaled, (log, MATLAB), (abs, MATLAB). These scaled parameters clustered into 5 groups, identified using Fuzzy C-Means (fcm, MATLAB, https://www.mathworks.com/help/fuzzy/fcm.html ). To measure the validity of this clustering we calculated the Modified Partition Coefficient (Friedman, https://github.com/lddavila/human_dec_making/blob/main/Utility%20Functions/calculate_mpc.m ), returning a score of 0.7061. In order to calculate the average sigmoidal shape per cluster we selected all points participating in a cluster, and averaged their psychometric functions across all sessions in each cluster.

#### Extended Data Figures 6A, 6B

1. Open the file “session_cost_clustering_figures_threshold_0.m”, linked here: https://github.com/lddavila/human_dec_making/blob/main/session_cost_clustering_figures_threshold_0.m
2. Run the section titled “SETUP PART 1: add the Utility functions to my path”, located in lines 1-9
3. Run the section titled “SET UP PART 5: put rat and human data into a table to be used”, located in lines 32-35
4. Run the section titled “SET UP PART 6: get human and rat data information (i.e. # of subjects, # of data points)”, located in lines 36-38
5. Run the section titled “Supplemental Figure 6A: rat vs human decision-making largely differs”, located in lines 292-298. This produces Extended Data Fig. 6A
6. Run the section titled “Supplemental Figure 2B: rat 1,3,4 overlap human 2,3,5”, located in lines 299-304. This produced Extended Data Fig. 6B.

#### Statistics for Extended Data Fig. 6B

To determine if there was overlap between rat and human clusters, we took the clustered 2500 human sessions across 35 subjects and the clustered 1616 rat sessions across 42 subjects then measured the Bhattacharyya Distance along each dimension between clusters and found the mean of them (bhattacharyyaDistance, MATLAB, https://www.mathworks.com/help/predmaint/ref/bhattacharyyadistance.html ). The resulting mean Bhattacharyya distances can be seen in Extended Data Fig. 6B.

### Figure 5D, 5E, 5F

1. Open “session_cost_clustering_figures_threshold_0.m”, linked here: https://github.com/lddavila/human_dec_making/blob/main/session_cost_clustering_figures_threshold_0.m
2. Run the section titled “SETUP PART 1: add the Utility functions to my path”, located in lines 1-9
3. Run the section titled “SET UP PART 5: put rat and human data into a table to be used”, located in lines 32-35
4. Run the section titled “SET UP PART 6: get human and rat data information (i.e. # of subjects, # of data points)”, located in lines 36-38
5. Run the section titled “Main Figures 2D, 2E, 2F”, located in lines 441-442.

#### Statistics for Fig. 5D, 5E

To define statistical difference between human male and female subjects’ cluster sizes, we used 2500 sessions across 4 cost levels and 4 reward levels. To match the number of male sessions (695 sessions), we randomly sampled the female sessions (1805 sessions) using function datasample (datasample, MATLAB, https://www.mathworks.com/help/stats/datasample.html ). To define statistical significance between male and female cluster proportions we used the chi squared test (chi2test, MathWorks File Exhchange https://www.mathworks.com/matlabcentral/fileexchange/16177-chi2test ), inputting the number of points of each cluster for each sex, returning a significance level of 0.0031349. To determine the strength of this test we ran a power analysis (sampsizepwr, MATLAB, https://www.mathworks.com/help/stats/sampsizepwr.html ), where we input the variance of the two groups, which returned a power level of 0.85536 as displayed on figures 5d, and 5e.

##### Statistics for Fig. 5F

To determine if hunger caused significant changes in cluster counts between sex, we took 695 male sessions across 11 subjects and 1805 female sessions across 24 subjects and split them by their hunger level. We took the cluster counts of the sessions which were greater than or equal to each hunger level and compared them to the cluster counts which were less than each hunger level using a chi squared test (chi2test, MathWorks File Exchange, https://www.mathworks.com/matlabcentral/fileexchange/16177-chi2test. The results show hunger has a significant change in males, but not in females. Figure 5F reports the significance levels.

#### Extended Data Figure 6C, 6D

1. Open the file “session_cost_clustering_figures_threshold_0.m”, linked here: https://github.com/lddavila/human_dec_making/blob/main/session_cost_clustering_figures_threshold_0.m
2. Run the section titled “SETUP PART 1: add the Utility functions to my path”, located in lines 1-9
3. Run the section titled “SET UP PART 5: put rat and human data into a table to be used”, located in lines 32-35
4. Run the section titled “SET UP PART 6: get human and rat data information (i.e. # of subjects, # of data points)”, located in lines 36-38
5. Run the section titled “Supplemental Figures 6C, 6D Sex Differences in human clusters do exist”, located in lines 414-417

##### Statistics for Extended Data Figs. 6C, 6D

To define statistical difference between human male and female subjects’ cluster sizes, we split session by gender (males had 695 sessions, females had 1805 sessions) and used the chi squared test (chi2test, MathWorks File Exhchange https://www.mathworks.com/matlabcentral/fileexchange/16177-chi2test ). We input the number of points of each cluster for each sex, returning a significance level of 0.0010587.

#### Figure 5G, 5H

1. Open “session_cost_clustering_figures_threshold_0.m”, linked here: https://github.com/lddavila/human_dec_making/blob/main/session_cost_clustering_figures_threshold_0.m

2. Run the section titled "SETUP PART 1: add the Utility functions to my path" located from lines 1-9.
3. Run the section titled "SET UP PART 5: put rat and human data into a table to be used", located in lines 32-35.
4. Run the section titled "SET UP PART 6: get human and rat data information (i.e. # of subjects, # of data points)", located from lines 36-38.
5. Run the section titled "Figure 2G, Figure 2H get male vs female bar plots for hunger", located from lines 468-485.

#### Figure 5I, 5J

1. Open "session_cost_clustering_figures_threshold_0.m", linked here: https://github.com/lddavila/human_dec_making/blob/main/session_cost_clustering_figures_threshold_0.m
2. Run the section titled "SETUP PART 1: add the Utility functions to my path" located from lines 1-9.
3. Run the section titled "SET UP PART 5: put rat and human data into a table to be used", located in lines 32-35.
4. Run the section titled "SET UP PART 6: get human and rat data information (i.e. # of subjects, # of data points)", located from lines 36-38.
5. Run the section titled "Figure 2I, Figure2J get male vs female bar plots for tiredness", located from lines 487-505.

#### Figure 5K, 5L, 5M, Extended Data Figure 6N, 6O, 6P

1. Open "session_cost_clustering_figures_threshold_0.m", linked here: https://github.com/lddavila/human_dec_making/blob/main/session_cost_clustering_figures_threshold_0.m
2. Run the section titled "SETUP PART 1: add the Utility functions to my path", located in lines 1-9
3. Run the section titled "SET UP PART 5: put rat and human data into a table to be used", located in lines 32-35
4. Run the section titled "SET UP PART 6: get human and rat data information (i.e. # of subjects, # of data points)", located in lines 36-38
5. Run the section titled "Main Figures 2K, 2L, 2M, Supplemental Figures 6n, 6o, 6p sex differences in rat clusters do not exist", located in lines 310-313.

#### Statistics for Figs. 5K, 5L, 5M

To define statistical difference between rat male and female subjects' cluster size we used 1616 sessions across 42 subjects. To match the number of male sessions (751 sessions) we randomly sampled the female sessions, (865 sessions) (datasample, MATLAB, https://www.mathworks.com/help/stats/datasample.html ). We used the chi squared test (chi2test, MathWorks File Exchange, https://www.mathworks.com/matlabcentral/fileexchange/16177-chi2test ), inputting the number of points of each cluster for each sex, returning a significance level of 0.63406. To determine the strength of this test we ran a power analysis (sampsizepwr, MATLAB, https://www.mathworks.com/help/stats/sampsizepwr.html ), where we input the variances of each group, and received a power level of 0.86087 as seen in figures 5h, 5i.

#### Extended Data Figure 6E

1. Open“session_cost_clustering_figures_threshold_0.m”, linked here: https://github.com/lddavila/human_dec_making/blob/main/session_cost_clustering_figures_threshold_0.m
2. Run the section titled “SETUP PART 1: add the Utility functions to my path”, located in lines 1-9.
3. Run the section titled “SET UP PART 5: put rat and human data into a table to be used”, located in lines 32-35,.
4. Run the section titled “SET UP PART 6: get human and rat data information (i.e. # of subjects, # of data points)”, located in lines 36-38.
5. Run the section titled “create histograms to measure the affect of differences between upper and lower half”, located in lines 342-403. Keep in mind that this section can take a while to run.
6. Run the section titled “Supplemental Figures 6G,6H: create a male and female first column heat map”, located in lines 433-436.

#### Extended Data Figure 6F

1. Open “session_cost_clustering_figures_threshold_0.m”, linked here: https://github.com/lddavila/human_dec_making/blob/main/session_cost_clustering_figures_threshold_0.m
2. Run the section titled “SETUP PART 1: add the Utility functions to my path”, located in lines 1-9.
3. Run the section titled “SET UP PART 5: put rat and human data into a table to be used”, located in lines 32-35.
4. Run the section titled “SET UP PART 6: get human and rat data information (i.e. # of subjects, # of data points)”, located in lines 36-38.
5. Run the section titled “create histograms to measure the affect of differences between upper and lower half”, located in lines 342-403. Keep in mind that this section can take a while to run.
6. Run the section titled “Fig 2F: create a male and female first column heat map”, located in lines 433-436.

##### Statistics for Extended Data Fig. 6E, 6F

To determine if tiredness caused significant changes in cluster counts between sex, we took 695 male sessions across 11 subjects and 1805 female sessions  across 24 subjects and split them by their tiredness level. We took the cluster counts of the sessions which were greater than or equal to each tiredness level and compared them to the cluster counts which were less than each tiredness level using a chi squared test (chi2test, MathWorks File Exchange, https://www.mathworks.com/matlabcentral/fileexchange/16177-chi2test. The results show hunger has a significant change in males, but not in females. Extended Data Fig. 6E, 6F report the significance levels.

#### Extended Data Fig. 6G, 6H

1. Open the file ]"session_cost_clustering_figures_threshold_0.m", linked here: https://github.com/lddavila/human_dec_making/blob/main/session_cost_clustering_figures_threshold_0.m
2. Run the section titled "SETUP PART 1: add the Utility functions to my path", located in lines 1-9
3. Run the section titled "SET UP PART 5: put rat and human data into a table to be used", located in lines 32-35
4. Run the section titled "SET UP PART 6: get human and rat data information (i.e. # of subjects, # of data points)", located in lines 36-38
5. Run the section titled "Supplemental Figures 6e, 6f: plot male/female hunger vs tiredness", located in lines 447-467.

#### Fig. 5N, 5O, 5P

1. Navigate into the file titled "session_cost_clustering_figures_threshold_0.m"
2. Run the section titled "SETUP PART 1: add the Utility functions to my path", located in lines 1-9
3. Run the section titled "SET UP PART 5: put rat and human data into a table to be used", located in lines 32-35
4. Run the section titled "SET UP PART 6: get human and rat data information (i.e. # of subjects, # of data points)", located in lines 36-38
5. Run the section titled "Figures 2J, 2K: Baseline Male and Female vs Food Dep Male and Female", located in lines 437-440.

#### Statistics for Figs. 5N, 5O

To determine if hunger caused significant changes in cluster counts between male rats, we randomly sampled the baseline sessions (751 sessions) to match the food deprived sessions(137 sessions)(MATLAB, datasample, https://www.mathworks.com/help/stats/datasample.html ). We then took the cluster counts of the food deprived and baseline groups then used a chi squared test (chi2test, MathWorks File Exchange, https://www.mathworks.com/matlabcentral/fileexchange/16177-chi2test. The results show hunger has a significant change in males, with a significance level of 0.011092. To determine the strength of this result we ran a power analysis (sampsizepwr, MATLAB, https://www.mathworks.com/help/stats/sampsizepwr.html ), where we input the variances of each group and received a power level of 0.98795 as seen in figures 5n, 5o.

#### Extended Data Fig. 6I, 6J

1. Open the file titled "check_chi_squared_between_rat_baseline_and_food_dep.m", linked here: ttps://github.com/lddavila/human_dec_making/blob/main/check_chi_squared_between_rat_baseline_and_food_dep.m
2. Run the file.

##### Statistics for Extended Data Fig. 6I, 6J

To determine if hunger caused significant changes in cluster counts between rats, we took the baseline sessions (1616 sessions) and the food deprivation sessions (297 sessions) and performed a chi squared test (chi2test, MathWorks File Exchange, https://www.mathworks.com/matlabcentral/fileexchange/16177-chi2test. We input the cluster counts of the food deprived and baseline groups, and the test returned a significance level of 0.0069.

#### Extended Data Fig. 6K, 6L, 6M

1. Open the file titled "session_cost_clustering_figures_threshold_0.m", linked here: https://github.com/lddavila/human_dec_making/blob/main/session_cost_clustering_figures_threshold_0.m
2. Run the section titled "SETUP PART 1: add the Utility functions to my path", located in lines 1-9
3. Run the section titled "SET UP PART 5: put rat and human data into a table to be used", located in lines 32-35
4. Run the section titled "Fig 2E: get food dep data", located in lines 169-174.
5. Run the section titled "Supplemental Figures 6G, 6H", located in lines 305-308.

##### Statistics for Extended Data Fig. 6K, 6L, 6M

To determine if hunger caused significant changes in cluster counts between male and female food deprived rats, we randomly sampled (datasample, MATLAB, https://www.mathworks.com/help/stats/datasample.html) female sessions (160 sessions) to match the male sessions (137 sessions). We then took the cluster counts of the male and female groups and used a chi squared test (chi2test, MathWorks File Exchange, https://www.mathworks.com/matlabcentral/fileexchange/16177-chi2test), which returned a significance level of 0.025637. To determine the strength of this result we ran a power analysis (sampsizepwr, MATLAB, https://www.mathworks.com/help/stats/sampsizepwr.html ), where we input the variance of the male and female groups and received a power level of 0.99905.

#### Extended Data Fig. 6N, 6O, 6P

1. Open the file titled “session_cost_clustering_figures_threshold_0.m”, linked here: https://github.com/lddavila/human_dec_making/blob/main/session_cost_clustering_figures_threshold_0.m
2. Run the section titled “SETUP PART 1: add the Utility functions to my path”, located in lines 1-9
3. Run the section titled “SET UP PART 5: put rat and human data into a table to be used”, located in lines 32-35
4. Run the section titled “SET UP PART 6: get human and rat data information (i.e. # of subjects, # of data points)”, located in lines 36-38
5. Run the section titled “Figures 2J, 2K: Baseline Male and Female vs Food Dep Male and Female”, located in lines 437-440.

#### Statistics for Extended Data Fig. 6N, 6O, 6P

To determine if hunger caused statistically significant changes in cluster counts between female rats, we randomly sampled the larger baseline female group which had 865 sessions to match the food deprived group which had 160 sessions (MATLAB, datasample, https://www.mathworks.com/help/stats/datasample.html ). We then took the cluster counts of the food deprived and baseline groups then used a chi squared test (chi2test, MathWorks File Exchange, https://www.mathworks.com/matlabcentral/fileexchange/16177-chi2test. The results show hunger has non-significant change in females, with a significance level of 0.1155. To determine the strength of this statistic we ran a power analysis (sampsizepwr, MATLAB, https://www.mathworks.com/help/stats/sampsizepwr.html ), where we input the variance of the food deprived and baseline groups and received a power level of 0.99715.

#### Extended Data Fig. 7A, 7B, 7C, 7D

1. Open file “rat_v_human_runme.m”, linked here: https://github.com/lrakocev/human_dm/blob/permutations/primitive%20building/rat_v_human_runme.m
2. Change the location in lines 20 and 32 to point to the following table: https://github.com/lrakocev/human_dm/blob/permutations/figs/all_session_updated/2d_clustering/human_2d_clusters.xlsx
3. Change the location in lines 21 and 33 to point to the following table:

https://github.com/lrakocev/human_dm/blob/permutations/figs/all_session_updated/2d_clustering/rat_2d_clusters.xlsx

4. Run the section titled “rat-human distance comparison”, lines 17-28. This produces Extended Data Fig. 7A.
5. Change the location in lines 37 to point to the following table: https://github.com/lrakocev/human_dm/blob/permutations/ingest%20helpers/human%20data.mat
6. Change the location in line 46 to point to the following table: https://github.com/lrakocev/human_dm/blob/permutations/baseline%20reward_choice%20psychometric%20functions%20table.xlsx
7. Run the section titled “prims histogram”, lines 29-73. This produces Extended Data Figs. 7C and 7D.
8. Run the section titled “2d homology”, lines 77-86. This produces Extended Data Fig. 7B.

#### Figure 6A

1. Open “ai_runme.m”, linked here: https://github.com/lrakocev/human_dm/blob/permutations/ai%20primitives/ai_runme.m
2. Change the “save_to” variable located on line 1 to point toward towards a directory located on your local machine.
3. Load the result of a simulation run linked here: https://github.com/lrakocev/human_dm/blob/permutations/ai%20primitives/mat%20files/full%20space%20w%20types.mat
   Or re-run the simulation using the following script: https://github.com/lrakocev/human_dm/blob/permutations/ai%20primitives/create_sigmoidal_space.m
4. Run “ai_runme.m”

#### Figure 6C, 6D, 6E

1. Open “dirk_helper_runme.m”, linked here: https://github.com/lrakocev/human_dm/blob/permutations/ai%20primitives/dirk_helper_runme.m
2. Change the location in line 3 to point to the following table: https://github.com/lrakocev/human_dm/blob/permutations/all_session_updated.xlsx
3. Load the following data: https://github.com/lrakocev/human_dm/blob/permutations/ai%20primitives/mat%20files/subset_dirk_space.mat
4. Run section “compare modeled data to human data”, lines 29-39 for Figs 6C, 6D
5. Run section “cluster config plot for human-adjacent clusters”, lines 41-43 for Fig. 6E

#### Extended Data Figure 8A

1. Open "fig_4_runme.m", linked here: https://github.com/lrakocev/human_dm/blob/permutations/ai%20primitives/fig_4_runme.m
2. Load the following table: https://github.com/lrakocev/human_dm/blob/permutations/ai%20primitives/mat%20files/full%20space%20w%20types.mat
3. Run the section titled "plot biased version of space", lines 32-39.

#### Extended Data Figure 8B, 8C, 8D

1. Open "space_v_hum_runme.m", linked here: https://github.com/lrakocev/human_dm/blob/permutations/ai%20primitives/space_vs_hum_runme.m
2. Load the following tables:
   a. https://github.com/lrakocev/human_dm/blob/permutations/ai%20primitives/mat%20files/full%20space%20w%20types.mat
   b. https://github.com/lrakocev/human_dm/blob/permutations/ai%20primitives/figs/orig_sampled/local_minima_clustered.xlsx
   c. https://github.com/lrakocev/human_dm/blob/permutations/all_cost_5_clusters.xlsx
3. Run section "compare local minima clustering to human", lines 23-38. This produces Extended Data. Fig. 8D.
4. Run section "compare human clusters to overall space", lines 40-49. This produces Extended Data Fig. 8B.
5. Run section "avg psychs from far-off clusters". This produces Extended Data Fig. 8C. Figs shown are examples selected from these results.

#### Extended Data Figure 8F

1. Open the file "example_activity.m", linked here: https://github.com/lrakocev/human_dm/blob/permutations/modeling/example_activity.m
2. Update the "save_to" variable located on line to a local directory where you'd like to save results.
3. Run the entire file, from lines 1-80.

#### Extended Data Figure 8G, 8H, 8I, 8J

1. Open the file “dirk_helper_runme.m”, linked here: https://github.com/lrakocev/human_dm/blob/permutations/ai%20primitives/dirk_helper_runme.m
2. Change the location in line 3 to point to the following table: https://github.com/lrakocev/human_dm/blob/permutations/all_session_updated.xlsx
3. Load the following two tables as done in section “load pre-calc'd clusters”, lines 19-25:
    a. https://github.com/lrakocev/human_dm/blob/permutations/ai%20primitives/mat%20files/subset_dirk_space.mat
    b. https://github.com/lrakocev/human_dm/blob/permutations/ai%20primitives/10_13_24/all_experiment_clustered_together.xlsx
4. Run the section titled “compare modeled data to human data”, located on lines 27-41.
5. Run the section titled “individual cluster config plots”, located on lines 45-47.

### Glossary

**Story**: A scenario. In our application, stories present a subject with a conflict resolution problem. A story encompasses the story context, the reward and cost preferences, and a series of trial questions. Each story belongs to a particular task type, and the context of each story surrounds a singular topic.

**Story order**: The list of stories assigned to a participant. The list is generated based on the participant topic preferences and the task types chosen, then it is randomised and recorded in the local participant file. The *story index* parameter dictates which story will be accessed next.

**Story index**: An integer that points to a particular position in the *story order* list. This parameter starts at 0 (pointing to the first element in the list) and is incremented by 1 after all 16 trial questions have been answered.

**Unicode**: A text encoding standard which allows the use of emoji and other regional characters for various writing systems.

**Demographic data**: All data taken from the user including but not limited to age range, sex, gender identity, education, hobbies, and visual media interests. Demographic information on the user is stored both locally in a file named ‘*demographic_info.txt*’ inside ‘/data/{SUBJECT_ID_NUMBER}’, and in the remote or local PostgreSQL database.

**Root directory**: The main working directory for the HUMANS application. The root directory will typically be represented with the leftmost ‘/’ in a file path throughout this document.

**Directory**: A 'folder' in the host computer's filesystem. Directories follow a hierarchy where the leftmost directory in a path represents the *root directory*. Any subsequent directories in the path are contained within each other. For example the file path '*C:\Windows\System32*' is a path that points to a Windows machine's system 32 directory.

**Command line**: A software program which provides direct communication between the user and the operating system or an application. In Windows, there are two command line shells, the Command shell (cmd) and the Power Shell.

**Console**: Can also be used as shorthand for *command line*. The console is a type of user interface which displays an application's output.

**ANT+ protocol**: The wireless communication protocol used for communicating heart rate information from the heart rate monitor to the host computer.

**HDMA (acronym)**: Human Decision-Making App

**HUMANS (acronym)**: Human Model for Analyzing Neuroeconomic Situations
**Host**: The computer on which an application runs.

**Client**: A computer which communicates with a *host* computer to request data.

**Integer (int, data type)**: A data type which includes whole numbers only. Integers are, simply put, numbers to do mathematical or logical operations on or with.

**Boolean (bool, data type)**: Is evaluated in conditional statements. A boolean can either be True (1), or False (0).

**List (list, data structure)**: An ordered succession of values in memory. Each element of a list can be accessed with an *integer* value denoting a value's position in the list. An index of 0 points to the first element in the list.

**Dictionary (dict, data structure)**: An ordered structure of values where each value is identified by a key. Dictionaries can easily be interpreted as tables where each key is a column name.

**String (str, data type)**: A data type containing numerical, ASCII, and/or *unicode* characters. Strings are typically used to represent non-numerical information and can be of variable length, spanning from a single character, to a whole book's worth of characters.

**ASCII (acronym)**: American Standard Code for Information Interchange. A standard data-encoding format for data exchange between computers. It assigns numerical values to letters, numbers, punctuation marks, and special characters.
